# Modeling Coronavirus Spike Protein Dynamics: Implications for Immunogenicity and Immune Escape

**DOI:** 10.1101/2021.08.19.456973

**Authors:** G. Kunkel, M. Madani, S. J. White, P. H. Verardi, A. Tarakanova

## Abstract

The ongoing COVID-19 pandemic is a global public health emergency requiring urgent development of efficacious vaccines. While concentrated research efforts are underway to develop antibody-based vaccines that would neutralize SARS-CoV-2, and several first-generation vaccine candidates are currently in Phase III clinical trials or have received emergency use authorization, it is forecasted that COVID-19 will become an endemic disease requiring second-generation vaccines. The SARS-CoV-2 surface Spike (S) glycoprotein represents a prime target for vaccine development because antibodies that block viral attachment and entry, i.e. neutralizing antibodies, bind almost exclusively to the receptor binding domain (RBD). Here, we develop computational models for a large subset of S proteins associated with SARS-CoV-2, implemented through coarse-grained elastic network models and normal mode analysis. We then analyze local protein domain dynamics of the S protein systems and their thermal stability to characterize structural and dynamical variability among them. These results are compared against existing experimental data, and used to elucidate the impact and mechanisms of SARS-CoV-2 S protein mutations and their associated antibody binding behavior. We construct a SARS-CoV-2 antigenic map and offer predictions about the neutralization capabilities of antibody and S mutant combinations based on protein dynamic signatures. We then compare SARS-CoV-2 S protein dynamics to SARS-CoV and MERS-CoV S proteins to investigate differing antibody binding and cellular fusion mechanisms that may explain the high transmissibility of SARS-CoV-2. The outbreaks associated with SARS-CoV, MERS-CoV, and SARS-CoV-2 over the last two decades suggest that the threat presented by coronaviruses is ever-changing and long-term. Our results provide insights into the dynamics-driven mechanisms of immunogenicity associated with coronavirus S proteins, and present a new approach to characterize and screen potential mutant candidates for immunogen design, as well as to characterize emerging natural variants that may escape vaccine-induced antibody responses.

**STATEMENT OF SIGNIFICANCE:** We present novel dynamic mechanisms of coronavirus S proteins that encode antibody binding and cellular fusion properties. These mechanisms may offer an explanation for the widespread nature of SARS-CoV-2 and more limited spread of SARS-CoV and MERS-CoV. A comprehensive computational characterization of SARS-CoV-2 S protein structures and dynamics provides insights into structural and thermal stability associated with a variety of S protein mutants. These findings allow us to make recommendations about the future mutant design of SARS-CoV-2 S protein variants that are optimized to elicit neutralizing antibodies, resist structural rearrangements that aid cellular fusion, and are thermally stabilized. The integrated computational approach can be applied to optimize vaccine immunogen design and predict escape of vaccine-induced antibody responses by SARS-CoV-2 variants.

## 1. INTRODUCTION

The recent COVID-19 global pandemic has highlighted that coronaviruses pose a dangerous threat to humans and animals. An important feature of coronaviruses is their ability to adapt to new hosts and environments through mutations (19). Thus, the threats that coronaviruses pose are ever-changing and long-term, and global health requires the quick characterization of SARS-CoV-2 related proteins and systematic design of treatment and prevention options. Coronaviruses are characterized by the crown-like Spike (S) glycoproteins on the surface of the virus particles (19). The coronavirus S protein is a member of the class I viral membrane fusion protein family present in SARS, MERS, Influenza (19,22,23), HIV(19,24), and Ebola (25) viruses. S proteins attach to cell-surface receptors, facilitating the viral membrane’s fusion with the host membrane and entry of the viral capsid into the cell cytoplasm (19,26,27). The S protein is a trimeric structure with each monomer comprised of two functional subunits: the N-terminal S1 subunit responsible for binding to the host cell receptor, and the C-terminal S2 subunit with machinery for fusion with the host cellular membrane (6,26,29,30). The critical first step in the fusion process occurs through the Receptor Binding Domain (RBD) on the N-terminal S1 domain of the S protein, which binds to the host cell receptor (6). The binding event is followed by proteolytic cleavage of the S protein by host proteases, resulting in significant conformational rearrangement of the S protein, shedding of the S1 domain, exposure of the S2 domain, and subsequent engagement of its fusion machinery, leading to host cell entry that leads to viral replication and cell death (31). The S protein is cleaved at the S1/S2 site between subunits and the proteolytic S2’ cleavage site, activating the membrane fusion cascade (14,32-34). In the case of SARS-CoV-2, S proteins recognize and bind to the human ACE2 receptor, triggering the viral fusion and replication cascade, leading to the spread of COVID-19 (6).

The structural orientation and dynamic behavior of the RBD is critical for host cell receptor binding (7,15,31,33). The RBD is a metastable domain that fluctuates between open and closed states in the prefusion conformation (15,31). It commonly adopts a single RBD open conformation, but multi-RBD open conformations have been observed upon receptor binding or in response to mutational design (7,15,17,35). The receptor-binding motif (RBM) is fully exposed and binds to cell-surface receptors to allow entry into the cell only in the open conformation of the RBD (36). Given that the RBD interaction with the ACE2 receptor is an essential viral mechanism, the S protein represents a prime target for immunogen design. Antibodies that block viral attachment and entry – neutralizing antibodies – bind almost exclusively to the receptor-binding domain of the S protein (6,26,30). While it may be possible that neutralizing antibodies bind to the S2 domain, the majority of studies show epitopes of neutralizing antibodies in S1 regions, mainly proximal to the RBD (3,30,35,37,38). While the main mechanism for viral neutralization occurs through antibody blocking of the receptor binding site, other mechanisms include prevention of ACE2 binding through steric clashes, and inducement of conformational shifts that prevent binding (2,4,5,35,39).

In recent years, structural biology has been instrumental in vaccine development, and in particular, atomic-level control of immunogens via structure-based design is increasingly feasible (17,39). There are multiple challenges and considerations for the informed design of immunogenic S protein variants. Prior mutagenesis studies of MERS-CoV, SARS-CoV, and SARS-CoV-2 S protein variants demonstrated that stability of prefusion structure plays a key role in viral fusion (7,17,33). A number of different mutations of the SARS-CoV-2 S protein have been designed in an effort to understand viral mechanisms and determine the best neutralizing variants. These include N-terminal domain (NTD) mutations (31,40), trimerization motif editing (5,9,15,31,33,40,41), proline mutations (7,17), and cleavage site mutations (7,9,31). Proline mutations in the S2 domain of the SARS-CoV-2 S protein, in particular, have been widely used for successful high-resolution cryo-EM structure determination and for generating structures with increased thermostability (7,17). Overall, evidence in the literature suggests the benefits of stabilizing mutations not only for prefusion state stabilization but also increased protein expression – both critical considerations for effective vaccine design. While stabilization of the prefusion conformation has been successfully implemented through a structure-based design approach for MERS-CoV and SARS-CoV-2, there is as of yet no highly effective immunogen – although S proteins are being used to develop first-generation vaccine candidates at this time (42). However, further characterization is needed to elucidate viral and neutralizing antibody mechanisms for effective vaccine design (42). To that end, we developed dynamic models for a large subset of S proteins associated with SARS-CoV, MERS-CoV, and SARS-CoV-2 implemented through coarse-grained elastic network models and normal mode analysis (NMA). The use of NMA in protein science is a standard method for generating protein dynamics by calculating vibrational modes (34,43,44). This method is useful for investigating protein motions around an equilibrium starting structure (43,44), where fluctuations obtained through NMA characterize a large fraction of the biologically-accessible movements experienced by structured proteins and proteins that contain flexible regions (43,44). This is a well-accepted method for describing biologically-relevant fluctuations of proteins, successfully applied to investigations of mechanically-driven deformations, energy transport properties, studies of large molecular complexes, and ligand-gated ion channels (34,45-47). We apply these models to systematically analyze local protein domain dynamics of S protein systems, as well as their thermal stability, to characterize structural and dynamical variability among different variants.

Traditionally, protein domains are associated with conserved regions of protein sequence and building blocks of multimeric structures (48). However, protein evolution does not always discretize dynamics over these domains (38, 39). Here, we consider rigid structural regions, termed dynamic domains, that behave in a quasi-independent manner around stabilized points, or hinges, and experience characteristic motion (15,49-52). The concretization of dynamic domains can highlight the functional roles of a localized area. For example, the RBD of the S protein is a qualitatively observed dynamic domain (7,15,31,53) whose fluctuation may provide a key viral mechanism for immune evasion. The identification of dynamic domains can give a measure of protein stability by pinpointing regions that are mobile and unstable compared to less dynamic and more stabilized regions. Throughout, we refer to structural stability as a characterization of a protein structure that resists deformation and reorganization. There are few computational analysis methods directed towards this task. Existing software use Gaussian Network Model methods to construct coarse-grained models, which may result in segmentation and accuracy artifacts (51,52). Other methods rely on the use of machine learning predictors that are trained on a limited set of NMR structures (52,54). Overall shortcomings associated with existing methods are ease of use, robust capture of accurate dynamic motions, domain differences between homologous structures, and quality of training data. To overcome these limitations, we designed a new algorithm that is applicable to biological structures in general, but is specifically developed for coronavirus S proteins.

We compare domain dynamics between SARS-CoV-2 mutants with SARS-CoV and MERS-CoV S proteins to establish the properties of various mutations and relate these to viral cellular fusion mechanisms. We then compare modeling results to available antibody-binding and epitope data to create a SARS-CoV-2 antigenic map and offer predictions for targeted molecular design of effective immunogens. Overall, this framework can be applied to the analysis and comparison of viral S proteins and associated mutants to determine structural and dynamic artifacts of mutations, as well as to link S protein dynamics patterns to antibody binding, toward more effective, computationally-driven immunogen design.

## 2. METHODS

### 2.1. Elastic network modeling and normal mode analysis

The use of normal mode analysis (NMA) in protein science is a standard method for generating protein dynamics by calculating vibrational modes (45). This approach uses a harmonic potential to compute protein movements. Although this approach is not as robust as, for example, molecular dynamics simulation using a more complex protein potential, it is able to produce accurate, large-scale protein motions around a starting structure (43,44). Fluctuations obtained via NMA can explore a small radius of movements around the equilibrium position within a protein’s free energy landscape (46). For structured proteins, or those with flexible regions, this radius can characterize representative, biologically-accessible protein motions (46). Indeed, studies show that the linear combination of low frequency modes is adequate to characterize collective motions and intrinsically favored dynamic patterns of functional units of membrane proteins and large systems like ion channels (43,47), receptors (43,55,56), and transporters (43,46,57). Additionally, other recent studies show that this is a valid method for describing realistic fluctuations of open and closed state S proteins, as well as for studying their nanomechanical properties (34,45).

In this study, anisotropic network models (ANMs) are constructed to coarse grain coronavirus S proteins. Normal mode analysis (NMA) is applied to ANMs to calculate vibrational normal modes and derive protein dynamics. Anisotropic network models are a variant of elastic network models that coarse grain the protein structure on a per-residue basis to construct a mass and spring system to dramatically speed up the NMA calculations (our method takes approximately 10 minutes to execute for each S protein). It differs from the simplified 1-D Gaussian Network Model (GNM) as each bead is represented as three points rather than as a single point, thus accounting for directionality and generating a more robust and accurate set of motions (50,51,58). The ANM construction represents each alpha carbon as a 3-point vibrational node, reducing computational cost and loss of accuracy in comparison to explicitly modeling every atom on each multi-atom amino acid (43,58). Our model implements connections between interacting nodes within a 15 Å cutoff distance. This includes interchain connections as the S protein is a multichain structure. If inter-protomer nodes are within the cutoff distance, then a connection between them is represented by a spring. These same criteria are applied for inter-subunit connections. Since the model construction is solely distance based, no additional springs or other refinements are included to account for or discriminate between specific intermolecular bonds (e.g. disulfide bonds). However, structural differences across variants that result from formation of new intermolecular bonds are reflected in the model construction. The node-spring composition for proteins is unique in the sense that mutation driven structural change, or PDB resolution, can influence the network model (results of this artifact are discussed in **Section 3.2**). When glycans are included in the PDB structures used for model construction, they are subject to the same modeling criteria.

We use the Python programming library ProDy (56) to construct ANM models along protein alpha carbons, form associated Hessian matrices (topological description), and conduct NMA (diagonalization of the Hessian) in Cartesian space. Our approach ensures that at least 98% of the total system dynamic response is captured in a collective motion, by first generating trajectories based on a linear combination of at least the first 15 normal modes weighted by their fractional variance (total contribution to motion). Trajectories are processed with MDtraj (59) and ProDy (56) programming libraries. All visualization is performed with VMD with the aid of the Normal Mode Wizard extension (60). All trajectory videos are included in the **Supplemental Videos**.

### 2.2. Dynamic domain analysis and calculations

We employ dynamic domain analysis (DDA) to characterize the specific cohesive, dynamic behavior of global domains in S protein systems considered here. Artifacts of a modal trajectory include vectoral and temporal data. Normal mode trajectories are parsed to find per-residue deformation vectors, deformation magnitude, coordinates of the starting structure, and coordinates of the deformed structure. The deformation data is obtained by comparing the starting structure to the most deformed structure in the normal mode trajectory. The most deformed structure is defined as the protein structure within the normal mode trajectory that has the highest root mean square displacement (RMSD) compared to the original. Deformation profiles show per-residue distances found by comparing corresponding node positions, between starting structure and most deformed structure:

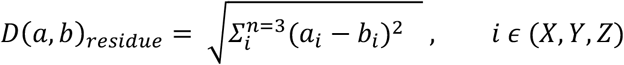

where *a* and *b* indicate a specific amino acid position, and *i* indicates their components in X,Y,Z Cartesian space.

Deformation profiles are de-noised to remove any outlier values that result from incomplete PDB structures. This data, in combination with starting structure coordinates, is used as input for a layered hierarchical agglomerative clustering algorithm that utilizes functions from the Scikit-learn programming library (61). Agglomerative clustering is the optimal choice because it avoids a specific cluster number constraint (unlike K-means or spectral clustering) and thus does not limit the number of identified dynamic domains (62). Each pass of agglomerative clustering uses a different linkage criterion— either Ward, complete, or single. These emphasize different properties to produce high-precision dynamic domain segmentation that can distinguish between small and large dynamical shifts while still respecting spatial barriers. The outputs of this analysis are locations of dynamic domains, local dynamics score (LDS), and global dynamics score (GDS). The LDS score is the average deformation that occurs over all the nodes in an identified dynamic domain. The GDS is the average deformation experienced by the whole structure, or root mean square deviation (RMSD). High GDS scores typically correspond to structures that experience large global rocking motions. A dynamic domain is defined as an identified cluster whose average deformation, LDS, is higher than the GDS. LDS scores that are lower and closer to the GDS score indicate a dynamic domain that is more stable relative to the deformation experienced by the entire structure. There is no ceiling for LDS scores. We note that a baseline for protein structural stability is measured by the GDS score and the level of deviation from structural stability is measured by the difference in GDS and LDS scores.

We also calculate the percentage overlap between dynamic domain residues across S protein variants and identified antibody binding zones. Percentages are calculated with respect to dynamic domains and antibody binding zones, by considering the number of common residues between the two groups. When finding the overlap with respect to dynamic domains, this number is scaled by the total number of residues within corresponding dynamic domains. When finding the overlap with respect to antibody binding zones, the number of common residues is scaled by the number of residues within the zones.

### 2.3. Sequence and structure analysis

One-to-One sequence comparisons are made using the BLAST Needleman-Wunsch Global Alignment software through the Blastp protein-protein webserver, where the wild type (WT) sequence is the subject sequence and the mutant is the query sequence (63). The BLAST tool gives an estimation of similarity between query and subject sequences. Multiple sequence alignment is performed on all presented sequences using the Clustal-Omega webserver on its default settings (64). The **Supplementary Information** contains the resulting multiple sequence alignment file and a description of all sequences used. Structural alignments and root mean square deviation (RMSD) calculations are performed using the “super” tool within the PyMol alignment software suite (65). Cryo-EM docking of crystal structures inside maps is done using Chimera fit in map functionality (66). Solvent accessible surface area (SASA) is computed using the Shrake-Rupley solvent accessible surface area function from the MDTraj programming library (59). Salt bridges are measured within a 3.2 cutoff radius (67,68) using the software VMD (60). In-house python scripts are used in all structural analysis.

### 2.4. Thermal stability prediction

Thermal stability is the ability of biological materials to resist degradation due to heat, pH change, and time evolution. It can be a determining factor for vaccine viability, and is therefore critical for experimental vaccine design. The Gibbs unfolding free energy (ΔG J/mol), and the difference between WT and mutant (ΔΔG J/mol), is considered as a measure for protein thermal stability. Here we create a thermal stability predictor to calculate ΔΔG and ΔG for protein sequences upon mutation as a measure of thermal stability. Existing computational methods for measurement of thermal stability rely on protein sequence or structure information and utilize machine learning or deep learning methods such as supportive vector machine (SVM) (69) or neural networks (NN) (50,70,71). We introduce a novel joint sequence- and structure-based thermal stability predictor to calculate single structure free energy and free energy change upon mutation. While both PDB structures and sequences are used in the training process, where PDB structures are used to derive many of the training features (see **SI Table A** in **Supplementary Information**), only the sequence is required for user input. So, our predictor returns the same outputs for both complete and incomplete PDB structures.

For training, we first employ a novel sequence embedding technique where embedding vectors are calculated with two different embedding approaches: Sequence Graph transform (SGT) (72) and Bidirectional LSTM (BiLSTM) models (71). Then, the total features are parsed through a convolutional neural network (CNN) model. The predictor is trained on combined biochemical features, biological features, structural properties, and energy terms (see **SI Table A** in **Supplementary Information**) that were extracted for each entry in our dataset. We use a combined dataset for training that includes the ProTherm dataset (73) and PoPMuSiC dataset (74), containing 1) PDB structure of wild-type protein, 2) mutation details such as location and residue type, 3) temperature, 4) pH, and 5) Gibbs free energy change upon mutation. Our CNN model includes hyperparameter tuning to increase performance and prediction accuracy. The optimal performance is found when hyperparameters including number of epochs, batch size, and learning rate are tuned. We found that 128 neurons per fully connected convolution layer and a 50% drop out rate prevent overfitting while still predicting optimal results. The total dataset consists of 16847 mutation points for 836 proteins, after removing redundancies. For testing, we use the independent I-mutant dataset (75). Our model also includes hyperparameters to tune the data and increase prediction accuracy. The close proximity of values in our binary output indicates that our predictor is well-trained and captures the most possible features from thermally stable and unstable classes. Importantly, we see no bias in contrast to other predictors. The 10-fold-cross-validation method is used to predict ΔΔG values and their associated standard error. The python libraries Keras (76), Tensorflow (70), Pandas (77) were used for algorithm construction and implementation.

## 3. RESULTS & DISCUSSION

### 3.1. Elastic network model dynamics qualitatively reflect cryo-EM data

We compare ANM models for S proteins and their corresponding NMA motions to true structures and their cryo-EM maps. Note that **Table 1** provides a description of all mutant SARS-CoV-2 PDB structures described in the following studies. The results from our analysis show that SARS-CoV-2 S protein models effectively capture the metastable nature of the RBD reflected in associated cryo-EM maps (**Figure 1**). Among available SARS-CoV-2 S protein structures, we consider two RBD-closed structures SC2.S1.TM1 (PDB ID 6VXX) and u1S2q (PDB ID 6×2C); three 1 RBD up structures BiPro (PDB ID 6VSB), SC2.S1.TM1 (PDB ID 6VYB), and u1S2q (PDB ID 6×2A); as well as a single 2 RBD up structure u1S2q (PDB ID 6×2B).

**Table 1:** PDB structures used as input to ANM models in this study. Thermal stability (**ΔΔG**) values are measured in comparison to WT thermal stability of ΔG=0.47 J/mol. Higher **ΔΔG** value denotes higher thermal stability. Thermal stability values less than 0.60 are considered to have a milder thermal stability improvement. Thermal stability values between 0.6 and 0.9 are considered to have moderate improvement. Thermal stability values greater than 0.9 are considered to have a significant improvement. Sequence length, percent of atomic coordinates unresolved within the PDB, and the similarity of each sequence to the WT sequence are presented. Supplementary Information contains additional sequence details. Note that furin cleavage mutations occur at the S1/S2 junction site.

| Mutant Name | PDB ID | Ref. | Sequence Length | Percent Unresolved | WT Similarity | Conformation | Expression Level | Mutation Description | Thermal Stability ($\Delta\Delta G$ J/mol) |
| --- | --- | --- | --- | --- | --- | --- | --- | --- | --- |
| SC2.S2.TM1-1 | 6ZGG | (40) | 1287 | 17% | 93% | 1 RBD up | Closed (34%)<br>Intermediate (39%)<br>Open (27%) | Signal Peptide, 2P, Trimerization Motif | $1.09 \pm 0.17$ |
| BiPro-1 | 6Z97 | (8) | 1273 | 22.9% | 94% | 1 RBD up | Unknown | 2P, GSAS furin cleavage, Trimerization Motif | $0.88 \pm 0.10$ |
| BiPro | 6VSB | (7) | 1288 | 25.4% | 94% | 1 RBD up | Unknown | 2P, GSAS furin cleavage, Trimerization Motif | $0.93 \pm 0.19$ |
| SC2.S1.TM1 | 6VXX/<br>6VYB | (6) | 1281 | 24.5% | 93% | All RBD down (6VXX)<br>1 RBD up (6VYB) | Unknown | Signal Peptide, 2P, GAGS furin cleavage, Trimerization Motif | $0.94 \pm 0.02$ |
| HexaPro | 6XKL | (17) | 1288 | 24.7% | 94% | 1 RBD up | Unknown | 6P, GSAS furin cleavage, Trimerization Motif | $1.21 \pm 0.10$ |
| SC2.N1.C1.2P.TM2 | 6XF6 | (21) | 1266 | 24.2% | 94% | 1 RBD up | Unknown | NTD clip, 2P, GSAS furin cleavage, Trimerization Motif | $0.99 \pm 0.11$ |
| u1S2q | 6X2B | (15) | 1273 | 24.6% | 93% | All RBD down (6X2C)<br>1 RBD up (6X2A)<br>2 RBD up (6X2B) | Open 67% | 2P, GSAS furin cleavage, Trimerization Motif, A570L T572I, F855Y, N856I | $0.32 \pm 0.11$ |
| SC2.C2.1P.TM3 | 7AD1 | (9) | 1297 | 26.5% | 93% | 1 RBD up | Closed (42%)<br>Intermediate (38%)<br>Open form (20%) | SRAG furin cleavage, 3P, Trimerization Motif | $0.82 \pm 0.14$ |
| SC2.C1.2P.TM4 | 6XM0 | (28) | 1288 | 18.7% | 93% | 1 RBD up | Unknown | NTD clip, 2P, GSAS furin cleavage, Trimerization Motif | $0.71 \pm 0.09$ |
| SC2.TM4-1 | 6XR8 | (14) | 1310 | Unknown | 97% | All RBD down | Unknown | Extended Trimerization Motif | $0.55 \pm 0.11$ |
| SC2.C1.2P | 7CN9 | (20) | 1127 | 13% | 88% | 1 RBD up | Unknown | NTD clip, 2P, GSAS furin cleavage, Trimerization Motif | $0.41 \pm 0.02$ |
| SC2.C1.TM4-2 | 7KDH | (78) | 1288 | 25.5% | 94% | 1 RBD up | Unknown | GSAS furin cleavage, Trimerization Motif | $0.93 \pm 0.14$ |
| BiPro-0 | 6ZP7 | (16) | 1273 | 22.7% | 99.7% | 1 RBD up | Unknown | 2P | $1.10 \pm 0.09$ |

**Figure 1:**
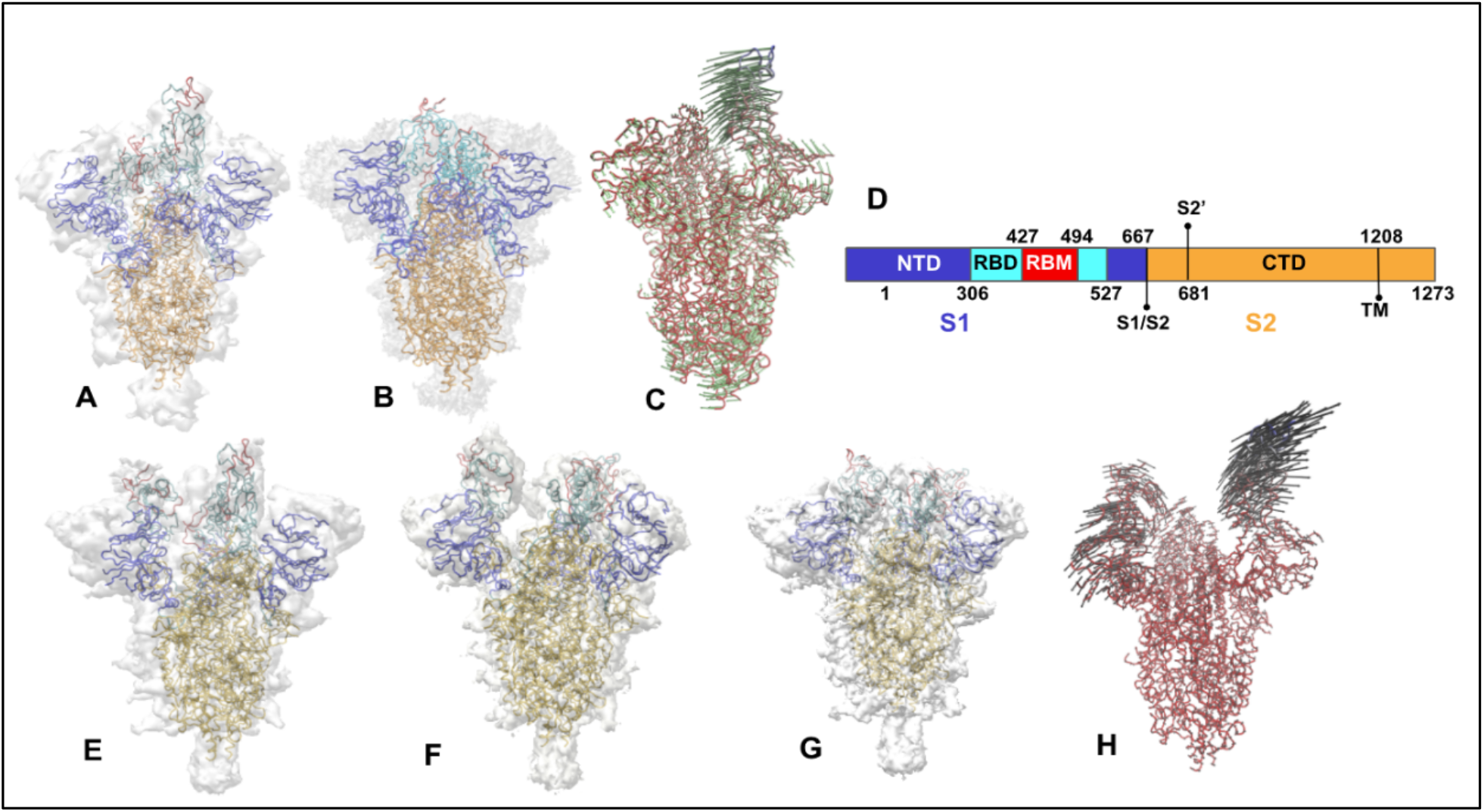
**(A)** 6VYB *(6)* starting structure inside its own cryo-EM map. **(B)** NMA derived 6VYB closed structure aligned inside 6VXX *(6)* cryo-EM map. **(C)** Path trace of 6VYB ANM modal trajectory with vectors indicating direction and degree of displacement. **(D)** SARS-CoV-2 sequence map showing protein domains as well as S1/S2 and S2’ cleavage sites and start point of trimerization motif (TM). **(E)** The 6×2B (15) 2 RBD up starting structure inside its cryo-EM map. **(F)** NMA derived 6×2B 1 RBD up structure aligned inside 6×2B (15) cryo-EM map. **(G)** NMA derived 6×2B closed structure aligned inside 6×2A (15) cryo-EM map. **(H)** Path trace of 6×2B ANM modal trajectory with vectors indicating direction and degree of displacement. Extra space between crystal structure and map can be attributed to the removal of surface glycans and/or regions of missing atomic structure. Proteins in **A-C** and **E-G** are colored according to the sequence map. TM = transmembrane domain.

First, ANM models based on BiPro (7) and SC2.S1.TM1 (6) are constructed using 1 RBD up starting structures to compare the match between 1 RBD up model trajectories (from 1 RBD up state to closed state) to corresponding cryo-EM maps. Qualitatively, the modal trajectory obtained from the BiPro ANM (see **6VSB Supplemental Video**) experiences the same characteristic hinge-like movements of the RBD between open and closed conformations identified in videos obtained by experimental cryo-EM studies on the BiPro sequence (*16*). SC2.S1.TM1 ANM dynamics similarly reflect the transitions between open (**Fig. 1A**) and closed (**Fig. 1B**) cryo-EM identified states (6) (see **6VYB Supplemental Video**). **Fig. 1A** shows the alignment of the SC2.S1.TM1 structure (PDB ID 6VYB) to its cryo-EM map. The closed RBD structure of SC2.S1.TM1 (PDB ID 6VYB) resulting from NMA is aligned inside the closed SC2.S1.TM1 (PDB ID 6VXX) cryo-EM map (**Fig. 1B**) to determine how well NMA can reproduce experimentally identified states. The root mean square deviation (RMSD) between the closed 6VYB model and the closed 6VXX structure is 1.42 Å, and 87.2% of the atoms in the closed 6VYB model fall within the 6VXX structural contour, as measured by Chimera fit in map function—establishing that open SC2.S1.TM1 structures can accurately capture large scale S protein movements.

Next, we compare the match between 2 RBD up model trajectories and dynamic domain analysis results of u1s2q (from 2 RBD up state, to 1 RBD up state, to closed state) to corresponding cryo-EM maps. Cryo-EM analysis of the u1S2q S protein reveals a 2 RBD up state (PDB ID 6×2B), 1 RBD up state (PDB ID 6×2A), and all RBD closed state (PDB ID 6×2C) (15). The propensity to adopt a 2 RBD up position is likely due to its unique set of A570L T572I, F855Y, N856I mutations (15). First, an ANM of the 2 RBD up u1S2q protein, 6×2B, was constructed and normal mode analysis was performed on the model to capture its dynamics. Frames that exhibit conformations which were the closest aligned with the 1 RBD up (PDB ID 6×2A) and closed (PDB ID 6×2C) crystal structures were extracted. The extracted conformations found via NMA were aligned with the crystal structures and their cryo-EM maps (**Fig. 1E-G**). The RMSD between the 1 RBD up model of u1S2q (PDB ID 6×2B) and 1 RBD up crystal structure (PDB ID 6×2A) is 0.66 Å and 89.4% alignment with the 6×2A map. The RMSD between the closed model of u1S2q (PDB ID 6×2B) and 6×2A is 0.72 Åand 85.9% aligned with the 6×2C map. In the NMA video of 6×2B (see **6×2B Supplemental Video**), the up RBDs are seen to fluctuate separately; one highly flexible RBD (LDS=1.51), closes while the other, less flexible RBD (LDS=0.83), flips upward (**Fig. 3C**). There is an intermediate state when both are in a mostly closed state. In this instance, the NMA method does not completely reproduce a down structure from the 2 RBD up structure. Overall, however, experimental studies show that the NTD and RBD domains undergo significant conformational change and exhibit flexibility, supporting the dynamic fluctuation observed in the dynamic network models (15).

### 3.2. Effect of glycans and structural resolution on WT anisotropic network model dynamics

All available experimental structures of SARS-CoV-2 S proteins contain unresolved structural regions. To verify that the anisotropic network models and normal mode analysis yield consistent characteristic motions, we consider the computationally refined WT S protein structure generated by Amaro et al (11). This structure is only missing data for the first 13 residues—the most resolved and accurate WT structure at the time of this study (11). NMA analysis and dynamic domain analysis is performed on a set of systematically reduced WT ANM models containing (1) intact glycans surrounding the protein, (2) removed glycans, (3) removed S2 subunit after residue 1146, and (4) removed commonly unresolved regions: 1-26, 67-81, 144-187, 243-262, 621-640, 672-689, 828-850, 1146-1273. Please see **Supplementary Information** for a full breakdown of missing regions for all PDB structures considered in this study.

First, NMA is conducted on the WT model that includes glycans surrounding the protein. The glycans exhibit independent, localized dynamic behavior while the protein’s mobility is significantly damped relative to the reduced structures (i.e., without glycans) (**Fig. 2A**). When the WT structure is analyzed without the glycans the general pattern of dynamics is preserved—as shown by deformation and solvent accessibility measurements (**Fig. 2B**). The dynamics are more pronounced and less damped, capturing more subtle local dynamics in the S2 subunit that would otherwise go undetected. Thus, glycan removal for NMA analysis can provide a more detailed breakdown of functional mechanisms. Additionally, glycosylation sites may differ from protein to protein so characterization of S protein dynamics without the presence of glycans can yield baseline motions that are consistent independent of glycosylation patterns (79). Still, including glycans in dynamics analysis may be useful to help identify the function of the glycans in different locations. For example, the glycans surrounding the location of the up-RBD (**Fig. 2A-red**) are predicted to be the most dynamic from NMA. Glycan studies by Amaro et al. note that glycans which surround the up-RBD help to stabilize it in the open conformation through hydrogen bonding (11). The high glycan flexibility exhibited in the models may provide further mechanisms for RBD stabilization. We also note that our analysis predicts dynamic domains within the extended region of the S2 subunit trimerization motif (**Fig. 2A & 2B**). However, under biological conditions, these are partially locked or stabilized within the virion membrane surface (19). Due to the geometry of the trimerization motif, a protruding structure that covers a large surface area, the NMA may bias towards predicting dynamics within this region rather than adjacent regions including the RBD. Thus, here we consider available PDB structures with unresolved trimerization motifs to evaluate realistic motions of RBD-adjacent domains.

**Figure 2:**
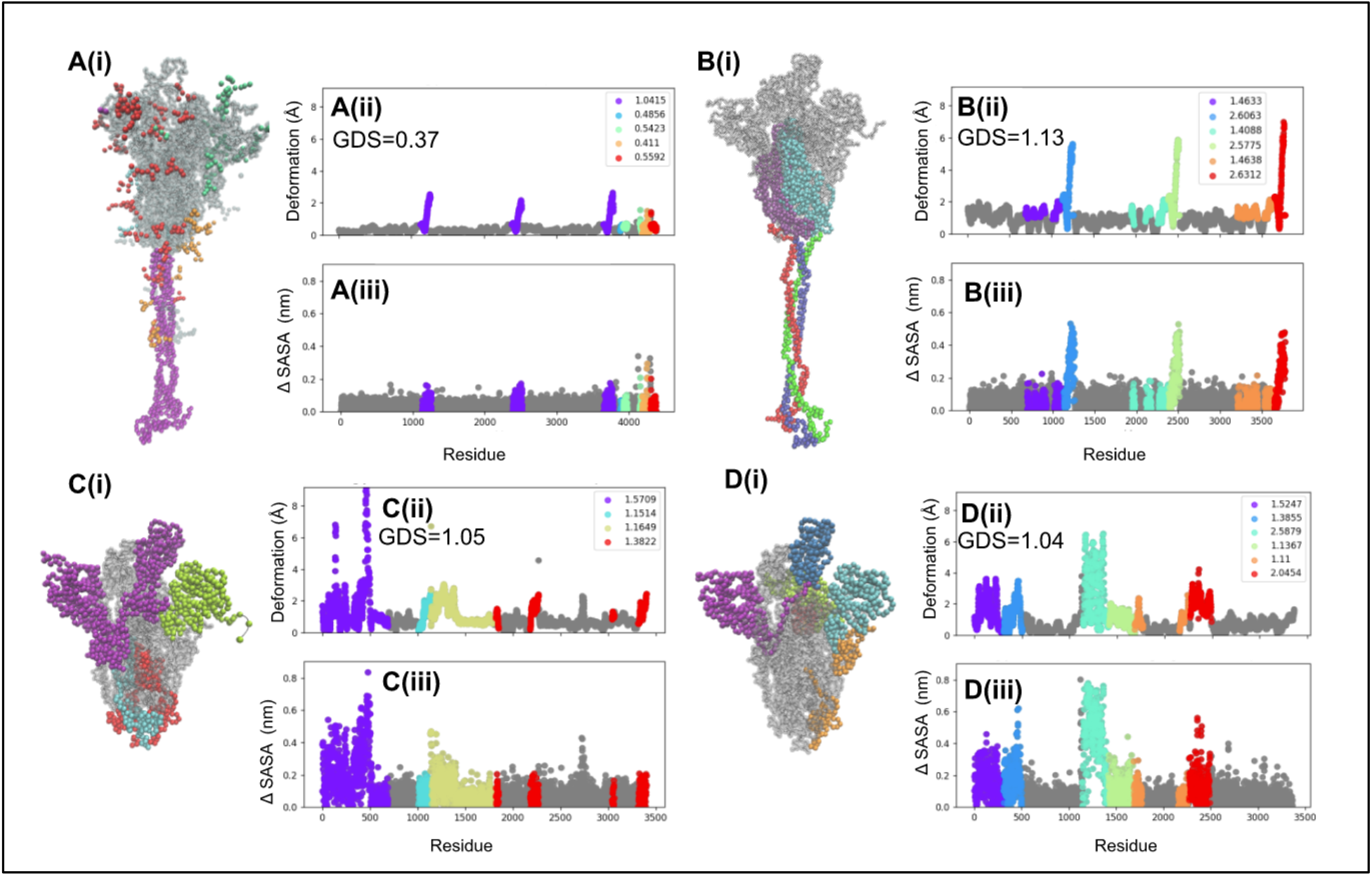
The evolution of NMA results and dynamic domain results for WT SARS-CoV-2 S protein models (11) with consecutive segments removed. The progression starts with **(A)** WT model with fully resolved trimerization motif and glycans, **(B)** removed glycans, **(C)** removed extended commonly unresolved S2 residues 1146-1273, and **(D)** removed additional commonly unresolved regions: 1-26, 67-81, 144-187, 243-262, 621-640, 672-689, 828-850. Each model is accompanied by local dynamics scores (LDS) in the legend in the upper right corner of per-residue deformation plots (ii) and solvent accessibility change plots (iii) to assess changes in protein movement calculations. Figure 2A is rotated to highlight all represented dynamic domains. The average distances traveled by the RBD oscillation are 3.84 Å and 3.50 Å for 2C and 2D, respectively.

**Figure 3:**
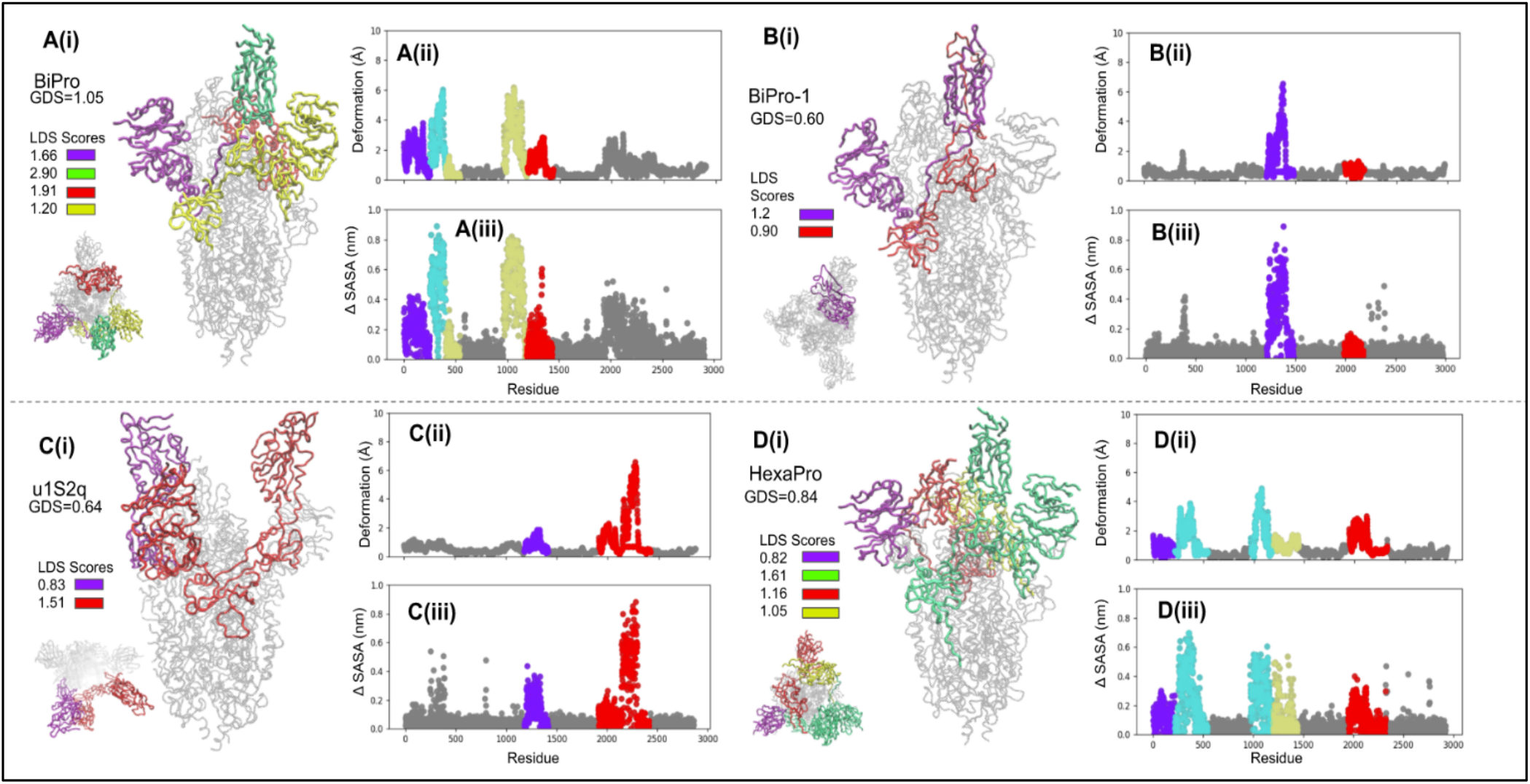
The domain dynamics associated with **(A)** BiPro (7), **(B)** BiPro-1 (8), **(C)** u1S2q (15), and **(D)** HexaPro (17) ANMs. The PDB ID, global dynamics score (GDS), local dynamics scores (LDSs), deformation profile **(ii)**, and Δ SASA profile **(iii)** is listed for each structure. On each 3-D structure and profile, identified dynamic domains are labeled in different colors and their LDSs are listed in each legend. On the profiles, the location of dynamic domains may look segmented, but these are often located in the same 3-D space. Dynamic domains are classified as regions which experience similar levels of deformation, but can also capture a small number of residues what are adjacent to dynamic regions and experience deformation in the same direction. Alternatively, residues that experience similar levels of deformation might be split into different dynamic domains because they are located in different regions of coordinate space or fluctuate in different directions. Associated dynamic videos are included in **Supplemental Information**.

We next consider the WT protein elastic network model with residues 1146-1273 removed and the WT elastic network model with other common unresolved regions removed (**Fig. 2C-D**). The deformation and solvent accessibility profiles suggest that these reduced models exhibit similar patterns of oscillation that are more consistent with what is seen experimentally for other S proteins with corresponding resolved regions. Dynamic domains are predicted around the location of WT S protein RBDs, where RBD oscillation typically occurs (6,33). In the WT model missing residues 1146-1273, the RBD moves a distance of 3.84 Å when alternating between open and closed states. The WT model missing both residues 1146-1273 and other commonly unresolved regions corresponds to a 3.5 Å displacement of the RBD. It also appears that removal of the NTD structure from positions 1-262 (approximately) encourages higher associated protein deformation. Additionally, removal of residues is hypothesized to accentuate weak regions and increased dynamic domain segmentation. This is also the case when we compared the dynamics results of BiPro and BiPro-1 mutant structures (see **Section 3.3**). This suggests that protein resolution levels can alter ANM-predicted dynamical patterns—although not dramatically. However, by confirming ANM dynamics with experimental data, even incomplete structures from cryo-EM may provide additional insights into S protein mechanisms.

### 3.3. Dynamics of S protein mutants and associated thermal stability predictions inform experimental observations

This section presents the dynamic domain patterns associated with different S protein mutants to compare with and confirm experimental findings, thereby further validating our approach. We also present the thermal stability results and discuss their implications. In the next section, 3.4, we explicitly synthesize this information to cluster families of mutations and to draw conclusions about their effect on dynamics and thermal stability, and the associated functional significance. We build anisotropic network models for SARS-CoV-2 mutants BiPro (7), SC2.S1.TM1 (open RBD, PDB ID 6VYB) (6), HexaPro (17), SC2.S1.TM1 (closed RBD, PDB ID 6VXX) (6), SC2.C2.1P. TM3 (9), SC2.N1.C1.2P.TM2 (21), BiPro-1 (8), SC2.C1.2P (20), SC2.C1.TM4-2 (78), SC2.TM4-1 (14), BiPro-0 (16), u1S2q (15), and SC2.C1.2P.TM4 (28) to first verify their agreement with experimental results and to gain insight into their associated immunogenic and mutation-related properties. These structures represent a comprehensive list of experimentally studied 1 RBD up S protein prefusion structures and 1 consensus model (**Table 1**). We consider primarily RBD up configurations as starting equilibrium structures for building network models because these can effectively sample both open (RBD up) and closed (RBD down) conformations. Dynamic analysis of closed structures, like 6VXX (**Fig. 5A**), did not produce RBD-up conformations but explores bending and twisting motions experienced in the closed state (see **6VXX Supplemental Video**). Generally, in our analysis, models that contain significant region(s) of deformation include dominant and auxiliary dynamic domains. Highly stable structures may only contain auxiliary domains and local dynamics scores (LDS) do not differ significantly from the global dynamics score (GDS). In addition to identifying the location of dynamic domains on 3-D S protein structures, we also present these domains on protein deformation and solvent accessibility plots, to quantify patterns in protein dynamics (**Fig. 3-5**). Dynamic domains typically correspond to regions that exhibit high deformation or solvent accessibility as compared to the rest of the structure. Finally, to complement our dynamic models, we evaluate the effects of mutations on the thermal stability of each sequence through our novel thermal stability predictor (**Table 1**).

**Figure 4:**
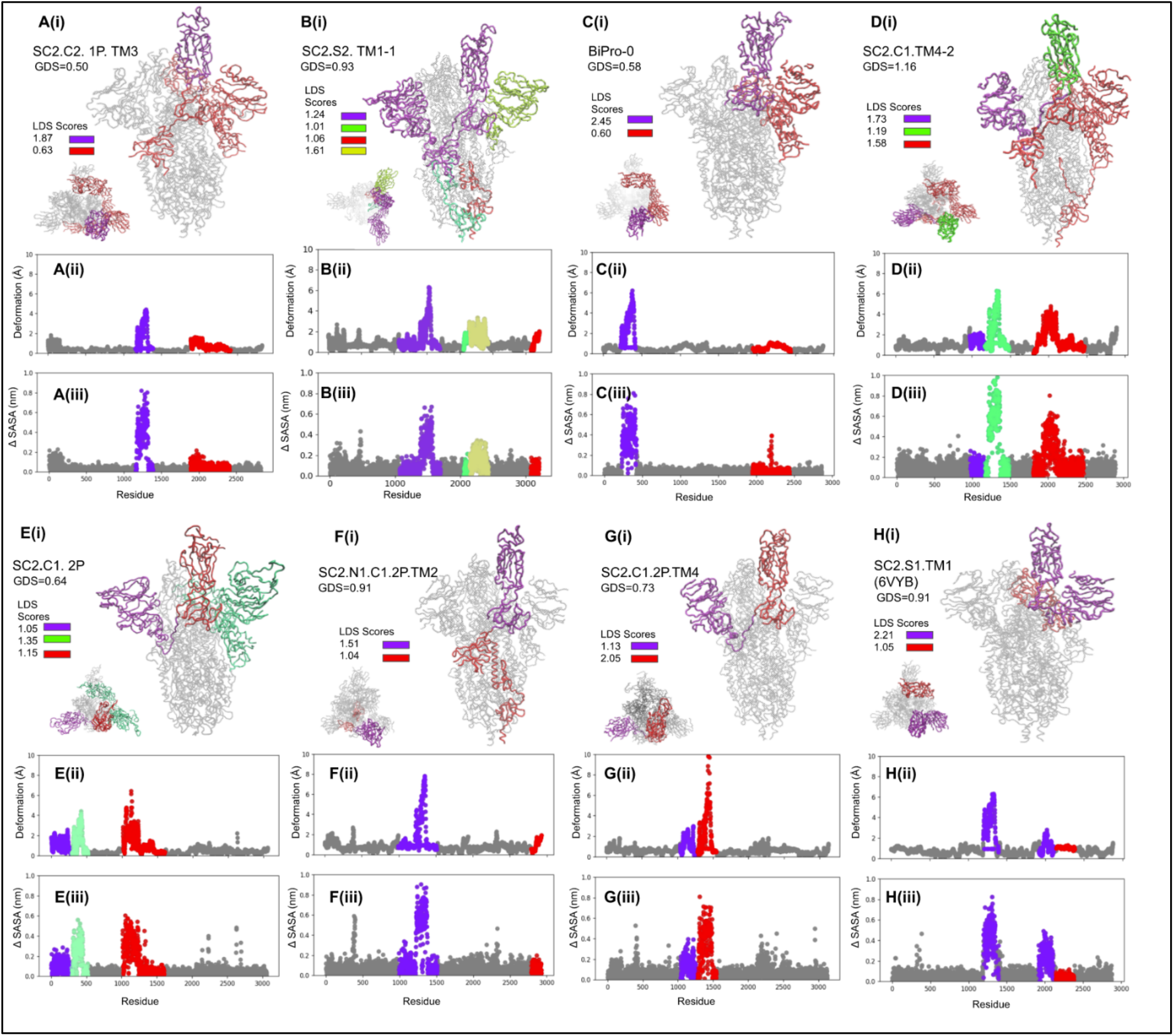
The domain dynamics associated with **(A)** SC2.C2.P1.TM3 (9), **(B)** SC2.S2.TM1-1 (13), **(C)** BiPro-0 (16), **(D)** SC2.C2.TM4-1 (14), **(E)** SC2.C1.2P (20), **(F)** SC2.N1.C1.2P.TM2 (21), **(G)** SC2.C1.2P.TM4 (28), and **(H)** SC2.S1.TM1 (6) ANMs. The analysis results for the open RBD structure of SC2.S1.TM1, 6VYB, are shown in **Figure 4H** and the closed structure in **Figure 5A**. The PDB ID, global dynamics score (GDS), local dynamics scores (LDSs), deformation profile **(ii)**, and Δ SASA profile **(iii)** is listed for each structure. Dynamic domains are color coded on 3-D structures according to LDS. Associated dynamic videos are included in **Supplemental Information**.

**Figure 5:**
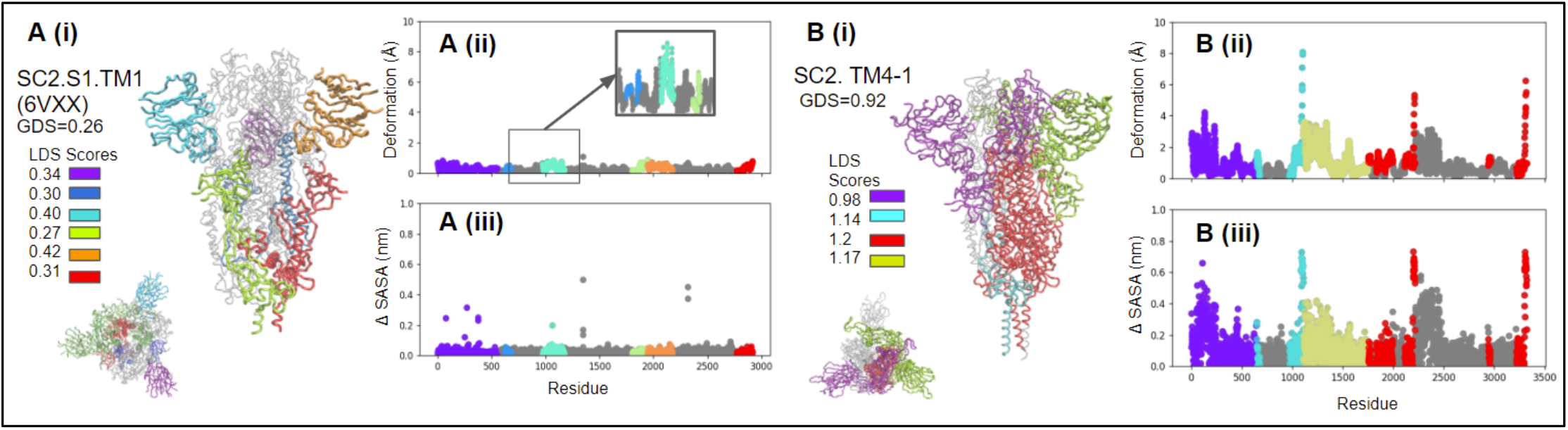
The domain dynamics associated with the ANMs of closed structure SC2.S1.TM1 (6) **(A)** and SC2.TM4-1 (14) **(B)**. The PDB ID, global dynamics score (GDS), local dynamics scores (LDSs), deformation profile **(ii)**, and Δ SASA profile **(iii)** is listed for each structure. Dynamic domains are color coded on 3-D structures according to LDS. Associated dynamic videos are included in **Supplemental Information**.

Analysis of the BiPro ANM trajectory **(Fig. 3A)** shows a dominant dynamic domain surrounding the up RBD with auxiliary domains around the NTD and an additional down RBD. Experimental studies confirm that thermal stability displayed by the BiPro RBD promotes an unstable three-RBD up conformation upon binding to the ACE2 receptor (7). In our analysis, the range of thermal stability across all mutants is ΔΔG=[0.32 J/mol, 1.21 J/mol]. The thermal stability increase of BiPro compared to the WT protein, at ΔΔG= 0.93 j/mol, represents a significant increase in thermal stability, correlating well with experimental observations. Our analysis shows that dynamic domains cover significant surface area in the S1 subunit. Their flexibility predicts the reorganization to the three-RBD up structure. However, there is large variability among the local domain dynamics in the S1 subunit, ranging from stable regions (LDS=GDS=1.05) to highly unstable, dynamic regions (LDS=2.9). The heterogeneity in dynamic behavior may contribute to the transience of the multi-RBD up configuration. Additionally, experimental flexibility analyses show that the resolution propensity of the S1 domains is decreased compared to the rest of the structure (7,16). The 6VSB ANM dynamic domain pattern confirms the mobility of these regions.

Next, we analyze BiPro-1(PDB ID 6ZP7) to further emphasize the contribution of protein resolution to observed dynamics. Both BiPro and BiPro-1 mutants contain the same family of mutations and have a sequence alignment of 99.8% to each other (**Table 1**; sequences are available in **SI**). Overall, we found that both proteins experience the same deformation range, contain a dynamic domain around the up RBD, and contain an auxiliary domain adjacent to a more dominant domain (see **Fig. 2A** and **Fig. 2B**). However, BiPro contains additional dynamic domains in the S1 subunit. This discrepancy may be due to the structural changes caused by differing trimerization motif sequences, but more likely, it is due to their differing levels of structural resolution (**Table 1**). The BiPro PBD structure contains missing regions in positions 330-334, 444-490, and 501-502, whereas BiPro PDB does not (7,8). This further confirms that mutation-caused structural shifts and protein resolution levels can alter elastic network model construction and thus predicted dynamics or protein stability – making experimental validation essential to NMA-derived dynamics. By confirming domain dynamics with experimental data, NMA can provide additional insight on incomplete crystal structures.

Local domain dynamics of the HexaPro (17) based model (**Fig. 3D**) are similar to BiPro. Considering that the sequences are similar (the BiPro sequence is mutated with four prolines in the S2 subunit to create HexaPro), it is expected that their associated dynamics would be similar. We discuss effects of proline mutations further in **Section 3.4**. The BiPro model does result in a dynamic domain around the RBD, whereas this is not the case in the HexaPro model. HexaPro shows a single dynamic domain surrounding the RBD and adjacent S1 regions. These dynamics are confirmed by cryo-EM studies that suggest that the S1 subunit is further secured (17). The HexaPro sequence also has the highest predicted thermal stability of the SARS-CoV-2 S proteins studied, ΔΔG=1.21 J/mol. Experimental thermal stability assessments show that the A942P mutation, not included in BiPro, was particularly powerful in increasing thermal stability (17).

The SC2.C1.TM4-2 (PDB ID 7KDH) (78) sequence contains GSAS in the furin cleavage site, trimerization motif mutations, and is missing proline mutations compared to the structures discussed thus far, although it is predicted to have higher thermal stability, ΔΔG=0.93 J/mol. Dynamics analysis of the ANM trajectory shows a dynamic domain extending into the S2 subunit (**Fig. 4D**). Aside from this difference, the S1 subunit local domain dynamics resemble BiPro most closely. Experimentally, the RBD experiences increased rigid body movement and the surrounding NTD regions experience smaller shifts (78)—this is consistent with our identified dynamic domain locations and LDS values. The RBD has a high LDS score of 1.73, relative to its GDS of 1.16, and the surrounding NTDs are identified as dynamic domains that have LDSs of 1.58 and 1.19.

The SC2.S1.TM1 (6) sequence contains the most diverse set of mutations associated with RBD up structures in this study (**Fig. 4H** and **Table 1**) containing a signal peptide, 2P, GAGS in the furin cleavage site, and trimerization motif mutations. Notably, the ANM of open SC2.S1.TM1 (corresponding to the 6VYB PDB structure) produces a lower number of dynamic domains than many of the other structures explored. These cover a smaller surface area; one is located around the up RBD and its adjacent NTD, the other covers one of the down RBDs. The up RBD is the dominant domain and displays the highest level of instability (LDS=2.21). The auxiliary domain around the down RBD is only slightly unstable, meaning that there is a small difference between the LDS (1.05) and GDS (0.91). Together these results indicate that that structure is mostly stable. Walls et al. note that the SC2.S1.TM1 S protein not only adopts 1 RBD up and down conformations but does not display a propensity to reorganize into multi-RBD up structures upon binding (6). Cryo-EM identified structures demonstrate that the closed RBDs lock down more firmly than in structures like BiPro (6)—this is consistent with our observations from the dynamic domain analysis. The majority of the S1 subunit structure is stable, and the dominant dynamic domain switches between the experimentally observed open and closed states. Since the majority of S1 regions are located in dynamically stable zones, they may not easily reorganize like the RBDs of the BiPro and HexaPro structures. Additionally, SC2.S1.TM1 is predicted to have moderate thermal stability, ΔΔG=0.94, compared to the WT, which is only slightly greater than the BiPro sequence (ΔΔG=0.93) that does not contain the signal peptide mutation.

Like SC2.S1.TM1, the SC2.S2.TM1-1 (PDB ID 6ZGG) sequence also contains a signal peptide, 2P, and trimerization motif (40). Unlike SC2.S1.TM1, it does not contain the furin cleavage mutation and its modal trajectory produces different domain dynamics (**Fig. 4B**). In SC2.S2.TM1-1, there are larger dynamic domains within the S1 subunits, especially around the RBD, as well as dynamic domains around the S1/S2 junction that extend into both S1 and S2 subunits. This is in contrast to SC2.S1.TM1 that presents smaller and more restricted domains only in the S1 subunit. Given that SC2.S1.TM1 contains the furin cleavage mutation and SC2.S2.TM1-1 does not, it may be the cause of these downstream dynamic changes. In fact, prior studies on SC2.S2.TM1-1 suggest that the absence of the GAGS furin cleavage mutation promotes disorder between the S domains and lowers thermal stability (6,12). However, we predict that its thermal stability, ΔΔG=1.09, is increased in comparison to SC2.S1.TM1, ΔΔG=0.94, which contains the same family of mutations plus the GAGS furin cleavage mutation. In comparison to the WT protein, which is also missing the furin cleavage mutation, SC2.S2.TM1-1 does indeed display an increased thermal stability. The experimental study also notes the structure is able to sample open, closed, and intermediate states with similar frequency (40). The domain dynamics analysis also supports this observation; the RBD adjacent S1 domains are unstable (exhibiting a dynamic domain) and may modulate intermediate states.

Like the BiPro and SC2.S1.TM1 sequences, the SC2.N1.C1.2P.TM2 (PDB ID 6XF6) (21) equence contains a GSAS furin cleavage mutation, 2P, and trimerization motif mutations (**Table 1**). Unlike these other sequences, it contains an NTD clip mutation. ANM dynamics analysis of the SC2.N1.C1.2P.TM2 model (**Fig. 4F**) shows a decreased number of dynamic domains that cover less surface area. Like the other sequences, it contains a highly dynamic up RBD and an auxiliary dynamic domain. This auxiliary (red) domain, however, is located in the S2 domain. We suspect that the different sequence architecture of the NTD causes structural artifacts and changes in auxiliary domain location.

Unfortunately, the corresponding experimental study to SC2.N1.C1.2P.TM2 was not yet published at the time of this research and, thus, these observations are not available for additional insights. Our thermal stability predictions show a moderately high improvement over the WT, ΔΔG=0.99 J/mol. Interestingly, this value is higher than that of BiPro (ΔΔG=0.93 J/mol) which contains the same family of mutations except for the NTD clip. Their trimerization motif substitutions differ. Thus, the NTD mutation and/or trimerization motif mutation may increase thermal stability in this case.

Unlike most of the other sequences considered here, SC2.C2.1P.TM3 (PDB ID 7AD1) possesses an SRAG furin cleavage mutation as opposed to the more commonly used GSAS mutation (9). From the domain dynamics (see **Fig. 4A**), the SRAG mutation in combination with 3 proline mutations provides an S2 stabilizing effect since there are no independent dynamic domains located in the S2 subunit. The addition of the proline mutation may also enhance the stabilization provided by SRAG. The corresponding experimental study notes that the mutations in the SC2.C2.1P.TM3 structure resulted in open, closed, and intermediate states with slightly more preference towards closed structures (9). This is supported by our dynamics analysis which predicts that most of the S1 subunit domains are stabilized (**Fig. 4A**). Experimental thermal stability analysis shows that the collective mutations performed on SC2.C2.1P.TM3 increase protein thermal stability, and the trimerization motif mutation in position 614 contributes greatly towards protein fusogenicity (9). Likewise, our results predict that these mutations moderately improve thermal stability, ΔΔG=0.82 J/mol, as compared to the rest of the mutants.

The BiPro-0 (PDB ID 6ZP7) (**Fig. 4C**) sequence is the most similar to the WT and only contains the 2P mutation (16). We note that the structure was produced by consensus computational modeling rather than experimental methods or molecular dynamics (16). Dynamics analysis identifies dynamic domains in the S1 subunit, the dominant domain being the RBD and the auxiliary domain covering the adjacent RBD and NTD. Interestingly, dynamic domains are not identified in the S2 subunit despite the limited number of mutations and no missing structural regions. This may be attributed to its mutations or, perhaps, an artifact of the computational modeling that caused some structural change. In a separate study, the principal component analysis of BiPro-0 structure showed that the NTD and RBD both fluctuate together, but the RBD shows a much more complex movement pattern (16). This further confirms our analysis, which predicts the RBD to have a higher degree of movement (LDS=1.89) than the auxiliary domain (LDS=0.6). Thermal stability predictions show BiPro-0 to have very high thermal stability, ΔΔG=1.10, possibly demonstrating the power of coupled proline mutations.

Lastly, we note the effect of the trimerization motif in PDB structures and resulting dynamic domain output. All of the structures listed in **Table 1** have large unresolved portions in the trimerization motifs—however, the SC2.TM4-1 closed structure, 6XR8 (14), has approximately 60 additional amino acids resolved in the trimerization motif compared to most structures. When comparing the SC2.TM4-1 (**Fig. 5B**) ANM dynamics to another closed structure, 6VXX (**Fig. 5A**), the general deformation trend and dynamic domain signature looks similar with the exception of the trimerization motif region. However, the overall level of deformation is increased for SC2.TM4-1 (GDS=0.92) compared to SC2.TM4-1 (GDS=0.26). In synthesis with the structural resolution insights gained from the WT structure, this result points to partially resolved trimerization regions contributing to more pronounced global dynamics and fully-extended, resolved trimerization motifs damping S protein global dynamics. Overall, we find that there are specific dynamic signatures associated with each presented S protein mutant. Both predicted dynamics and thermal stability predictions closely agree with experimental observations. These results provide the motivation and points of comparison to understand the effect of each family of mutations: proline mutation, NTD editing, and furin cleavage editing.

### 3.4. Mutations in SARS-CoV-2 S protein sequence induce variability in protein dynamics

The patterns in dynamic signatures computed for the presented protein mutant models suggest that there may be specific protein behaviors associated with each family of mutations. Therefore, we first compared the dynamical differences between S protein models associated with certain mutations to form an initial set of hypotheses to investigate. Then, because different structures contain unique sets of unresolved regions, we create artificial control(s) where the aggregates of all unresolved regions within the structures being compared are removed. This provides a baseline by which to understand the dynamics of compared models and gives an estimation of the error associated with each of the presented predictions.

#### Effects of the furin cleavage mutation

The presence of the furin cleavage mutation at the S1/S2 subunit is common among all S protein mutants surveyed in the literature. Our analysis shows that this mutation may have an effect on S protein dynamic patterns, structural stability, and thermal stability. A key motivation for additional analysis of these mechanisms stems from common patterns in the presented S protein mutants and their associated dynamics. We first consider all proteins that sample both open and closed conformations—all of which contain furin cleavage mutations (see **Table 1**) at the S1/S2 subunit junction except SC2.S2.TM1-1, BiPro-0, and the WT sequence. The remaining structures have SRAG (SC2.C1.2P.TM4), GAGS (SC2.S1.TM1), or GSAS—the most common furin cleavage mutation. We do not include BiPro-0 in this analysis considering that the structure was resolved by consensus modeling, which may affect its structure and corresponding dynamics. Interestingly, the WT and SC2.S2.TM1-1 structures do not contain the furin cleavage mutation and present dynamic domains in the S2 subunit. This motivates the hypothesis that RBD fluctuations transmit forces to the rest of the structure, including the S2 subunit. Stabilizing furin cleavage mutations can secure the S1/S2 junction, redistributing forces to S1 subunit regions to create a more stable S2 subunit. In the absence of the furin cleavage mutation, the S2 subunit may act as a shock absorber by presenting dynamic domains. For example, SC2.S2.TM1-1 lacks a furin cleavage mutation and has lower LDSs in the S1 domain, closer to the GSS, than structures whose dynamic domains are restricted to the S1 subunit. Most S proteins exhibit LDS scores in dominant dynamic domains that are at least 0.70 greater than their GDS scores, while the dominant dynamic domain in SC2.S2.TM1-1 displays an LDS that is lower. In fact, SC2.C2.TM3, BiPro-0, BiPro, SC2.C1.2P.TM4, and SC2.S1.TM1 all exhibit dominant dynamic domains with LDS scores exceeding GDS by at least 1.30. This suggests that the additional S2 subunit dynamic domains absorb some of the force that is transmitted to S1 subunit domains thereby reducing the level of deformation exhibited by S1 dynamic domains. This hypothesis is strengthened by experimental observations that note furin cleavage mutations can control RBD allosteric effects through S2 domain changes (78). We note that while other models, including SC2.C1.TM4-2 and SC2.N1.C1.2P.TM2, present dynamic domains in their S2 subunits, this behavior may be attributed to multiple factors, such as the NTD clip mutations and higher levels of unresolved structure.

To further understand the possible implications of having a furin cleavage mutation, we first compare the structure and dynamics of SC2.S2.TM1-1 (PDB ID 6ZGG) with SC2.S1.TM1 (PDB ID 6VYB). The SC2.S2.TM1-1 sequence contains the same family of mutations as the SC2.S1.TM1 sequence, minus the furin cleavage mutation. There is a large difference in the amount of unresolved regions between the two structures, 7.5% higher in SC2.S1.TM1 (**Table 1)**. To characterize the effect of unresolved regions, the common unresolved regions between SC2.S2.TM1-1 and SC2.S1.TM1 (for which there is overlap, see breakdown in **SI Section 7)** are removed from SC2.S2.TM1-1 to create control SC2.S2.TM1-1’ (**SI Fig. 1A)**. SC2.S2.TM1-1’ exhibits dynamic domains around the furin cleavage sites in the lower S1 regions and upper S2 regions similar to SC2.S2.TM1-1, but dynamics are damped in other regions of the S2 subunit. Although SC2.S2.TM1-1 and its control display some differences in their dynamic patterns, there are apparent commonalities when compared to SC2.S1.TM1. Both present increased dynamic domains in the S1 subunit and around the furin cleavage sites compared to the SC2.S1.TM1 model whose dynamic domains cover a smaller surface area only in the S1 subunit. These results suggest that the furin cleavage mutation may help mitigate the dynamics seen by S1 regions and provide mild stabilization to S2.

We further investigate the effect that the furin cleavage mutation has on the S2 subunit by comparing the dynamic patterns of SC2.S1.TM1 to the WT protein. The unresolved regions in SC2.S1.TM1 are removed from the WT to create WT’-A (**SI Fig. 1B**). The results indicate that the general pattern of dynamics is preserved in WT’-A. Specifically, dynamic domains within WT’-A cover the same S1 regions as in WT (**SI Fig. 1B, Fig. 2C-D**). Additionally, two dynamic domains exist at furin cleavage sites and extend into the S2 subunit, similar to the other WT models presented in **Section 3.2**. Although SC2.S1.TM1 contains additional mutations, it is clear that the WT protein displays a propensity for naturally flexible regions in the S2 subunit and an unstable S1/S2 junction.

Together, the comparison of dynamical patterns between SC2.S2.TM1-1, WT, and their controls demonstrates that structures without stabilizing furin cleavage mutations may have a propensity to exhibit flexibility in the S2 subunit. Furin cleavage mutations, individually, may provide a mild stabilizing effect to the S2 subunit by modulating the response at the S1/S2 junction. However, this action may be highly sensitive to the presence of other mutations since we see more consistent and pronounced dynamic domains in the S2 subunit of the WT than of SC2.S2.TM1-1. Furthermore, experimental observations of SC2.S2.TM1-1 note that GAGS furin cleavage mutation promotes disorder between the S domains (28). It may be possible that disorder-related flexibly of the furin cleavage site is a mechanism that controls the S1-S2 junction stability and force transmissibility from RBD motion of S1 to the S2 subunit. Lastly, we note that most S protein sequences containing furin cleavage mutations in this study display a moderate to high increase in thermal stability. However, given the variability in protein sequence, it is difficult to identify if this is only due to the presence of the furin cleavage mutation or multiple mutational factors.

#### Effects of NTD editing

Insertions and deletions of amino acids in the NTD of the S protein are also common mutations in the presented variants. In this section, we consider SC2.N1.C1.2P.TM2 (PDB ID 6XF6), SC2.C1.2P.TM4 (PDB ID 6XM0), and SC2.C1.2P (PDB ID 7CN9) which all contain NTD clip mutations. We do not identify a unifying defining feature among proteins that contain the NTD clip mutation—which is likely due to their sequence and structural variability. Signal peptide mutations are one subset of NTD mutations, occurring in S protein mutants SC2.S2.TM1-1 and SC2.S1.TM1. These present differing S2 subunit dynamics, which is likely attributed to the presence of a furin cleavage mutation in SC2.S1.TM1. Interestingly, SC2.N1.C1.2P.TM2 displays an auxiliary dynamic domain in the S2 region. This is not a common pattern associated with proteins that have the GSAS furin cleavage and 2P mutations. This result suggests that NTD-related mutations may augment the level of stability provided by the furin cleavage region. Removal of the NTD amino acids may disrupt protein stability, causing stabilizing furin cleavage mutations to be less effective. Addition of signal peptides may reinforce the furin cleavage site or may cause significant dynamical changes in S1 regions. Given the variability that results from introducing different signal peptides, or NTD deletions, it remains difficult to definitively predict the dynamical and functional implications of a certain class of NTD editing.

We find that the location of an NTD mutation has an effect on resulting dynamics. The first 32 residues are most commonly changed via signal peptide or deletion (see sequence breakdown in **Supplementary Information**). Within the WT protein, residues 13-32 are located on the underside of the NTD in close proximity to the S1/S2 junction (**Fig. 6**). By contrast, within the SC2. TM4-1 structure, for example, residue 14 (1-13 is unresolved) is located around the top portion of the NTD (15), suggesting a variability in organization of this region among S protein structures. The position of the first 32 residues may be critical for determining S1 dynamic stability. Structural change may indirectly drive dynamical and functional mechanisms through protein geometry or directly by changing the architecture of critical bonds. In particular, LYS77 and ARG80 form a salt bridge in the NTD of the WT structure. This pair exists in close proximity to residues 13-26, with residue 77 as close as 9 Å to residue 24. Alteration of the first amino acid positions by mutation may subvert critical bonds like the observed salt bridge. Additionally, any mutation to position 15 may result in the direct destabilization of a disulfide bond between CYS15 and CYS136 observed by Cai et al. (14).

**Figure 6:**
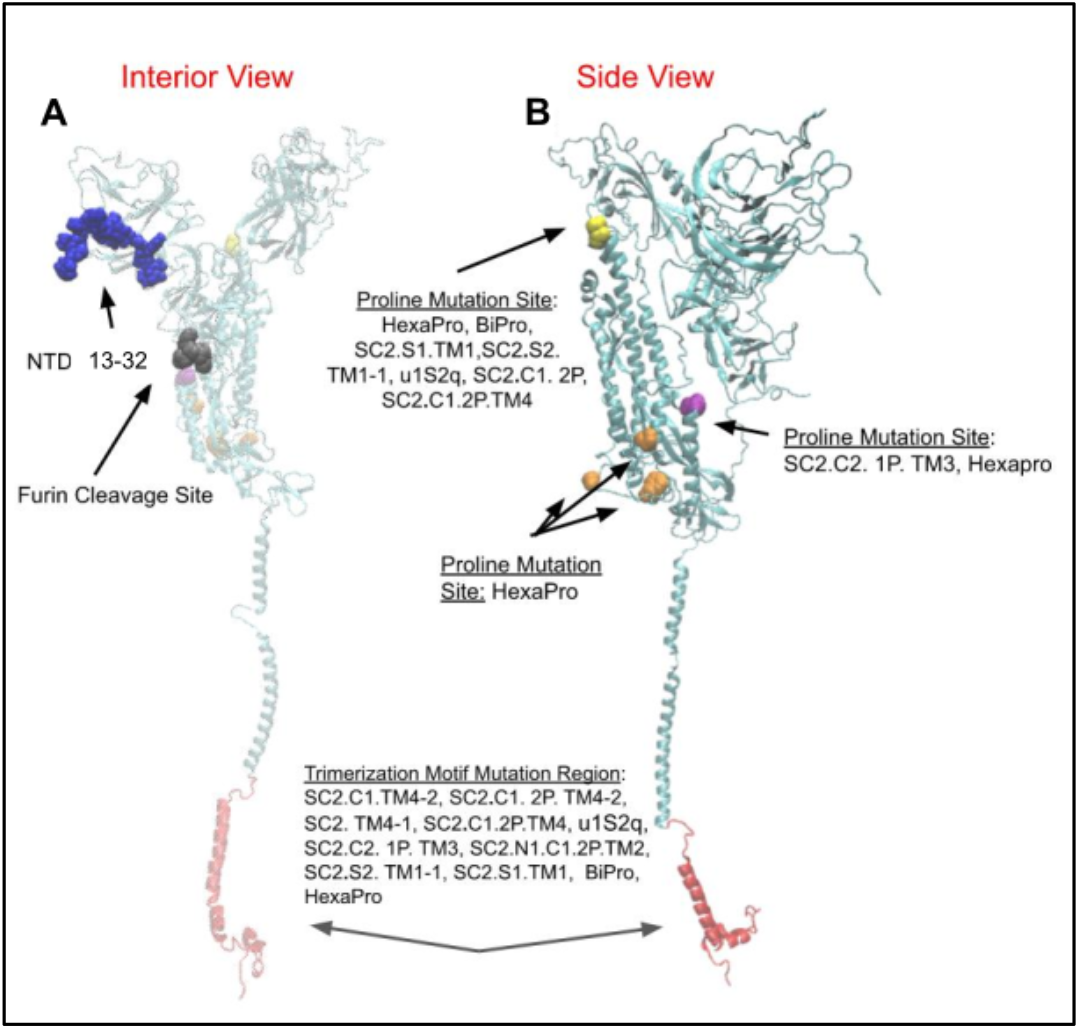
Representation of the WT S protein (67) structure interior view (A) and side view (B). All mutation sites in the S protein shown are listed in **Table 1**

To further investigate the effect of NTD clip mutations, we remove all unresolved regions between SC2.N1.C1.2P.TM2, SC2.C1.2P.TM4, and SC2.C1.2P from the SC2.C1.2P structure to create the control SC2.C1.2P’ and analyze resulting modal trajectories. We remove unresolved regions from SC2.C1.2P because it has the highest resolution, unresolved regions within only 13% of the structure, in comparison to SC2.N1.C1.2P.TM2 and SC2.C1.2P.TM4, with unresolved regions amounting to 24.5% and 18.7%, respectively. The dynamics of SC2.C1.2P’ show an increased number of dynamic domains in the S1 subunit compared to SC2.C1.2P (**SI Fig. 2A, Fig. 4E**). Next, the same unresolved regions are removed from SC2.C1.2P.TM4 to create the control SC2.C1.2P.TM4’ and dynamic domain analysis is conducted (**SI Fig. 2B**). The dynamic domain breakdown of SC2.C1.2P.TM4’ shows an additional dynamic domain in the S2 subunit that is not present in SC2.C1.2P.TM4 and the dynamic domain around the RBD covers a larger surface area (**SI Fig. 2B, Fig. 4G**). Overall, however, the general pattern of dynamics is not disrupted when SC2.C1.2P and SC2.C1.2P.TM4 are compared to their controls. Protein deformation is, however, emphasized in the controls, which accounts for the additional dynamic domains in regions where deformation is more damped in the original structures. For example, there is some observed protein deformation in SC2.C1.2P close to residue index positions 2250 and 2750 (**Fig. 3E**), while in SC2.C1.2P’ this motion is more pronounced (**SI Figure 2A**, in orange). Comparing the original S proteins to their controls, in addition to the analysis that we perform on the WT protein, shows how removal of regions can encourage instabilities by emphasizing already present dynamic patterns. Thus, the outcome of mutations that remove portions of protein structure may likely result in a similar phenomena. Comparison of proteins that have NTD clips does not elucidate any other apparent dynamic patterns given the other variations within the sequences. Thus, the outcome of this mutation is hypothesized to be highly sensitive to the location and magnitude of the deletion, which may be further influenced by other present mutations.

Overall our observations highlight NTD insertion and deletion effects on structural stability and protein mobility levels. Location of the mutations and other protein sequence artifacts are likely to impact the nature of dynamical and functional mechanisms that occur due to NTD mutations. Analysis of protein structure surrounding the location of critical NTD residues highlights how mutational changes may disrupt local bonding and/or supplement S1/S2 junction stability. The comparison of SC2.C1.2P.TM4 and SC2.C1.2P to their control structures in conjunction with the WT analysis in Section 3.2 demonstrates how the removal of protein structure can emphasize protein dynamics and encourage instabilities by providing less structural support.

#### Effects of proline mutations

Proline mutations are common among the S protein mutants we considered in this study, occurring in all mutants, except for SC2.C1.TM4-2 (78), SC2.TM4-1 (14), and the WT (11). In this section, we analyze thermal stability patterns and compare the dynamic domain composition of S protein mutants to investigate the mechanisms of S protein proline mutations. Experimental studies indicate that proline mutations increase protein thermal stability and may aid in S protein resistance against reorganization, especially when prolines are added to the backbone and/or loop positions (17,33,39,78). Our thermal stability predictions confirm that prolines increase thermal stability. There is a 0.28 J/mol thermal stability increase in the BiPro (7) sequence when prolines are added to create HexaPro (17), which has the highest measured thermal stability in our set, ΔΔG=1.21 J/mol. Also, the BiPro-0 sequence (16) displays a high thermal stability value, ΔΔG=1.10 J/mol, with just the 2P mutation. Of the structures with proline mutations, u1S2q has the lowest thermal stability value ΔΔG=0.32 J/mol. Its unique quadruple mutation results in a 2 RBD up structure but may also decrease thermal stability compared to other S protein mutants (**Table 1)**. This structure may benefit from additional prolines, e.g. as introduced in HexaPro.

While the effects of proline mutations on thermal stability have been investigated previously (6,7,17), the impact of proline mutations on dynamic domain decomposition is largely unknown. Of the structures that sample both open and closed conformation, only the SC2.C1.TM4-2 and WT sequence do not contain any proline mutations. All structures with proline mutations, except SC2.S2.TM1-1, contain the upper interior proline mutation (**Fig. 6C**, in yellow). The HexaPro mutant contains many unique proline mutations (**Fig. 6C**, in orange) and shares one proline site with SC2.C2.1P.TM3 (**Fig. 6C**, in purple). To investigate the role of proline mutations, we first compare the general dynamic patterns in S protein mutants to make generalizations about proline contributions in specific sequences. Since most of the structures contain proline mutations and there is variability in sequence and structural resolution among them, it is difficult to establish a basis of comparison for a more accurate and direct assessment of individual proline mutations. However, we do compare the effects of the HexaPro 6P mutation and BiPro 2P mutation to gain more direct insight into the effect of proline mutations on structural dynamics patterns (**Fig. 3A**,**D**). These structures have 100% alignment in their unresolved regions and thus can be directly compared without artificial controls. HexaPro presents an increased number of dynamic domains compared to BiPro, and its dynamic domains cover a wider surface area in the S1 subunit. It is possible that in the absence of other S1 stabilizing mutations, such as a signal peptide, the higher proline content of HexaPro stabilizes the S2 domain further and redistributes forces that contribute to the motion of S1 subunit domains—propelling these domains to sweep a larger surface area. Since the dynamic domains in the S1 subunit cover a larger surface area, the force per area may be lower and may contribute to decreased LDS scores in HexaPro. Comparing S2 subunit dynamics between BiPro and HexaPro, the GDSs for these regions are 1.05 and 0.94, respectively—suggesting increased S2 stabilization within HexaPro. Experimentally, it has also been suggested that the 6P mutation triggers further S1 instability compared to other structures (17).

### 3.5. S1 subunit antigenic map and mechanisms of virus neutralization

The variability in S protein dynamics suggests that domain accessibility and mobility patterns associated with S protein mutants may influence the number and positioning of neutralizing antibodies that can bind to S proteins. To investigate the relationship between solvent accessibility and the location of dynamic domains, we calculated the change in solvent accessibility over the course of modal trajectories (**Fig. 3-5(iii)**). Solvent accessible surface area (SASA) deformation closely correlates with NMA deformation patterns, and dynamic domains are characterized by larger variability in solvent accessibility, as expected.

To investigate whether and how the location and flexibility of dynamic domains influence antibody binding, we created an S1 subunit antigenic map and characterized the neutralizing mechanisms associated with different antibody-binding regions on the S protein (**Fig. 7**). The antigenic map was created from an exhaustive literature review of SARS-CoV-2 related antibodies and their epitope data (**SI Table 1)**. All epitope positions were mapped onto our WT model. Their location on this model, combined with binding characteristics from all available antibodies, formed the basis of the defined zones. Epitope positions in zone 1 are 392, 403-421, 428-430, 444-458, 472-486, and 515-517; zone 2 are 439, 470, 487-498, and 505-505; zone 3 are 440-445, 343-346, and 368-374; zone 4 are 347-360, 370-390, 405-418, and 376-380; and zone 5 are 145-150 and 246-250 (**Fig. 7**). We note that residues in each zone may shift due to RBD refolding in response to mutations, binding, or other structural modifications. The zones are limited to the S1 subunit since there is more literature characterizing antibodies that bind to S1, and in association, possible competition with ACE2 provides a direct neutralizing action (14,15). There is less literature characterizing SARS-CoV-2 antibodies that bind to the S2 region; it is presently unclear if this is due to research bias or because antibodies bind dominantly to the S1 domain. These may stabilize the structure into a neutralizing configuration or compete with ACE2 receptor binding directly (35,37). While antibodies can bind to other regions of the SARS-CoV-2 S protein, these are not well documented (80). The S protein antigenic map is suspected to exist as a continuum rather than as discrete zones. However, based on our current understanding of S1 epitopes, defining them by zones informs the differentiation between binding mechanisms to the S protein. The SC2.S1.TM1, BiPro, and HexaPro sequences are studied most commonly in antibody binding studies. In **SI Table 1** and **Fig. 7**, the binding zone, neutralizing effect, and related prefusion trimer structures are categorized. If an antibody is studied in relation to a freely expressed RBD that is associated with a prefusion trimer, or docked to one computationally, then it is marked as unknown. This analysis shows that potent neutralizing antibodies bind to the S protein in all zones. However, the mechanisms for neutralization are different and contact with any binding zone does not guarantee potent neutralization. All epitope positions were mapped onto our WT model, including the location of each antibody and binding characteristics (**Fig. 7B**).

**Figure 7:**
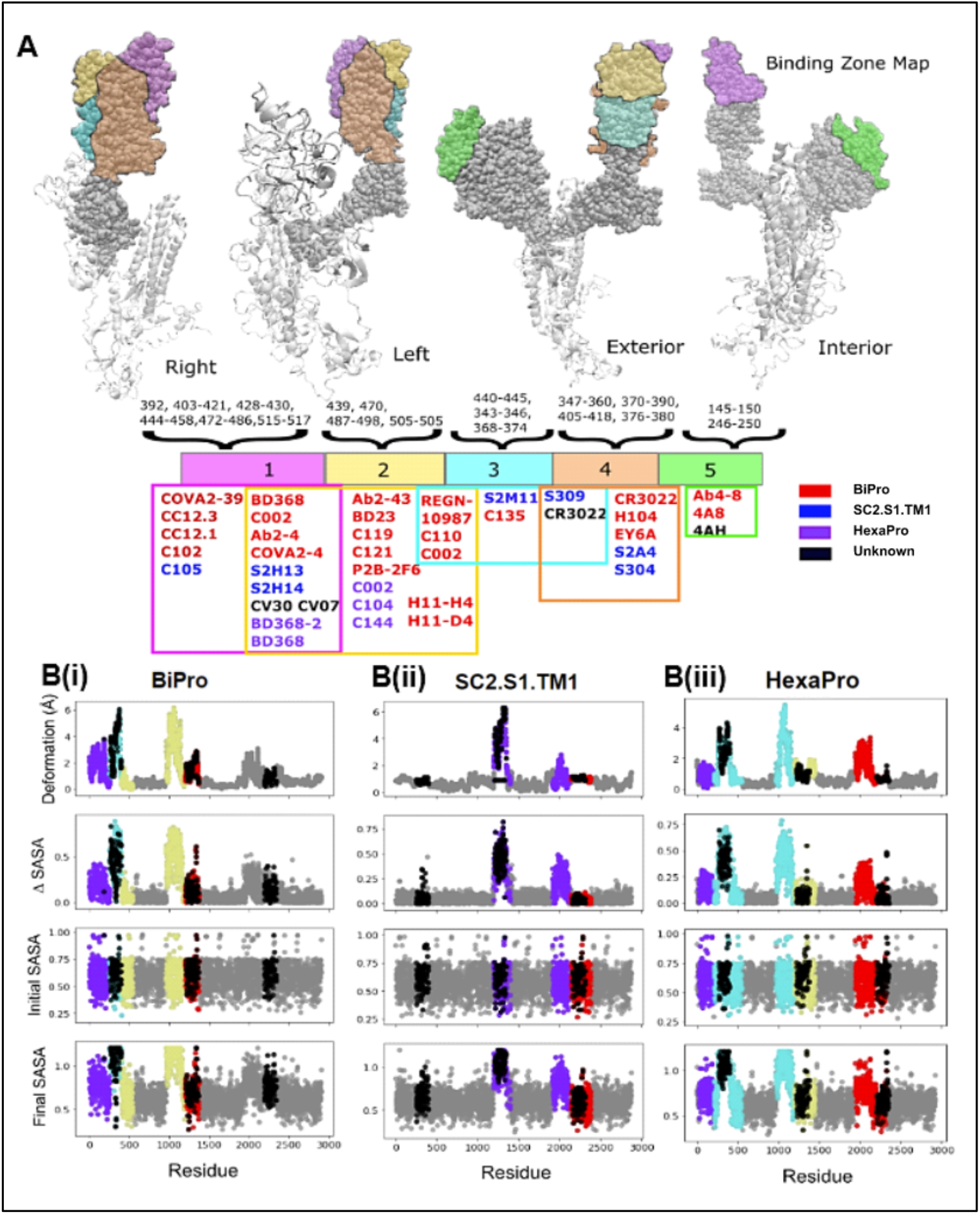
**(A)** S protein antigenic map with epitope zones labeled in different colors. The color bar provides a Venn-diagram showing the zone associated with each of the studied antibodies (see **Supplemental Table 1**). Above the color map sequence positions of defined epitopes are labeled in association with each zone. **(B)** The location of epitopes (black regions) from all zones are mapped onto deformation and solvent accessibility plots for **(i)** BiPro (7), **(ii)** SC2.S1.TM1(6), and **(iii)** HexaPro (17) to show the overlap between epitope zones and dynamic domains.

Zone 1 (**Fig. 7A**) exists on the inside and top of the RBD—it is largely hidden when an RBD is closed and is fully exposed when an RBD is rotated vertically by way of hinge fluctuations. Thus, antibodies can only bind to zone 1 fully when the RBD is in the up conformation. For many of the antibodies that bind to the RBD in zone 1, their mode of action is direct blocking of the ACE2 binding site (5,35,36,81,82). These antibodies fully or partially overlap with ACE2 binding positions (35). The antibody can simultaneously support neutralization by producing steric clashes with ACE2 such that it cannot bind to other exposed binding regions (35).

The neutralization action provided by antibodies that bind, or partially bind, to zone 2 (**Fig. 7A**) is varied. Zone 2 exists around the top and exterior of the RBD and it can be recognized while the RBD is in up and down configurations. Some antibodies, such as REGN10987, are suspected of shifting zones as the RBD fluctuates and moves into ACE2 competing positions (35,83). Many antibodies that target zone 2 act as bridges between adjacent RBDs or between other antibodies (35). This may influence the RBD’s ability to lock on to ACE2 by inducing conformational changes or by blocking ACE2 sterically (35). For example, C144 is able to attach to adjacent domains and lock the trimer into a closed position so that it cannot interact with ACE2 (35).

Zone 3 (**Fig. 7A**) is located underneath zone 2 on the exterior of the RBD, making it easily accessible in both open and closed states. Since it does not overlap with the ACE2 binding site, nor exist in close proximity, the neutralizing effects are suspected to be caused by conformational changes or blocking of ACE2 by steric clashes (5,35,40). In cases of weak neutralization, the conformational changes needed for potent neutralization may not be accessible or the ability to clash with ACE2 is mild (5). In cases of higher potency, this may not be the case. Importantly, antibodies that bind in zone 3 may allow space for other neutralizing antibodies to bind to RBD regions and work together to create a neutralizing cocktail (83). For example, S309 and S2E12 work together to stabilize RBDs in the down position and hide receptor binding sites (2).

Zone 4 (**Fig. 7A**) is located on the side regions of the RBD. These epitopes have been labeled as ‘cryptic epitopes’ in other studies (35,36). Zone 4 is only fully accessed in the RBD up configuration. However, this region may be partially accessed in the down position if an antibody is bound elsewhere. In most cases where this cryptic epitope is accessed, a multi-RBD up structure would be optimal as this would create space for more stable binding (10, 25). Unless the up RBD is fully extended, binding in zone 4 may cause further conformational shifts to an unstable multi-RBD up structure or a stabilized open structure (5,15,30,35). These scenarios may again inhibit the ability of the RBD to lock into ACE2 and form stable interactions.

Lastly, zone 5 (**Fig. 7A**) is located around the tip of the NTD. There have been fewer documented cases of antibodies binding to this region, but examples include 4AH (12,80) and Ab4-8 (37). Interestingly, antibodies that bind in this zone have been seen to produce potent, neutralizing effects. They can bind to the NTD in both up and down positions and do not clash with ACE2 binding regions. The neutralization mechanisms of these antibodies remain largely unclear, although Chi et al. note that antibodies which bind to the NTD may provide some stabilizing effect (12). Our dynamics analysis of S proteins with signal peptide mutations show that additions made to the NTD can provide a stabilizing effect (**Fig. 2** and **Table 1**). Thus, based on our analysis it is plausible that antibodies targeting the NTD can provide additional neutralization support.

Each S protein mutant has a different family of mutations (**Table 1**) and produces unique dynamic patterns associated with a specific level of stability. We find that these patterns directly correlate with the antibody binding propensity and binding mechanisms for different S protein mutants. This is quantified in **Fig. 7B** where the locations of known epitopes (in black) are mapped onto deformation and solvent accessibility profiles for BiPro, SC2.S1.TM1, and HexaPro. The locations of known neutralizing epitopes directly overlap with the locations of dynamic domains. At the same time, there are certain areas where dynamic domains do not overlap with an identified epitope location. For example, the yellow and purple dynamic domains on the BiPro structure (**Fig. 7B (i)**) cover entire NTDs and do not overlap with black epitope regions. Given that known epitopes and dynamic domains correlate significantly, these regions may indicate additional antibody targeting sites outside of those already found experimentally.

BiPro and HexaPro structures, containing epitopes in all of the defined zones, are recognized by a variety of antibodies. Interestingly, the epitopes of these structures may be predicted from their dynamic domain patterns (**Fig. 3A**,**D**). BiPro and HexaPro present dynamic domains that cover a larger surface area in the S1 subunit domains. The up RBDs are considered dynamic domains, or regions of a dynamic domain in the case of HexaPro, corresponding to zones 1 and 2. In some cases, binding in zone 2 may require structural reorganization or binding to adjacent RBDs. The presence of dynamic domains in these same regions highlights the structure’s ability to adjust and be made available to secondary antibody contacts or structural manipulations. Also, the presence of dynamic domains surrounding the NTD shows that this area may be receptive to binding with other proteins.

Unlike BiPro and HexaPro, the SC2.S1.TM1 S protein has epitopes concentrated in zones 1 and 4, with some overlapping with zone 3. Due to its location (**Fig. 7A**), zone 3 is easily accessed in both open and closed states. Zone 3 can likely be recognized regardless of S protein state or stability. The presence of a fluctuating RBD exposes zone 1. Thus, we would expect to identify epitopes associated with zone 1 since SC2.S1.TM1 samples open and closed states, and this is confirmed in our dynamic domain analysis (**Fig. 4H**). It may be the case that the structural stability of the closed RBDs and mobility experienced by the open RBD create an accessible space in zone 4 and allow for recognition by antibodies. The SC2.S1.TM1 structure is associated with more epitopes in zone 4 than BiPro and HexaPro structures by percentage. The SC2.S1.TM1 structure is also not associated with epitopes in zone 2. Antibodies that bind in zone 2 commonly create bridges with adjacent RBDs and present other conformational changes. In our dynamics analysis, we find that the down RBDs are more stabilized and inclined to adopt the down position over the up position; thereby reducing RBD movement and exposure of additional antibody binding zones. This observation suggests that secondary antibody contacts may be essential for the longevity and stability of neutralizing antibodies. Secondly, the stability associated with the closed RBDs within the S1 subunit, as in the SC2.S1.TM1 structure, hinders the ability of antibodies to stably connect and reorganize S1 subunit domains.

We next calculate the dynamic domain overlap percentage (**Fig. 8)** with the defined antibody zones for each SARS-CoV-2 S protein mutant listed in **Table 1**. The percent overlap is calculated with respect to the number of residues within dynamic domains, giving a measure of the total dynamic domain space that overlaps with known antibody binding zones. High percent overlap, scaled by the dynamic domains, indicates that the protein may not present additional antibody binding areas outside of the defined zones. Percent overlap is also calculated with respect to the number of residues within antibody binding zones, giving a measure of the total zone space overlapping with dynamic domains. High percent overlap, scaled by antibody zones, indicates that protein dynamic domains overlap significantly with the defined binding zones. Thus, the most desired combination, in consideration for S protein design, could target low overlap with respect to dynamic domains (which would indicate the potential for additional antibodies) and high overlap with respect to antibody binding zones. Among the structures considered here, this result is best exemplified by the u1S2q mutant.

**Figure 8:**
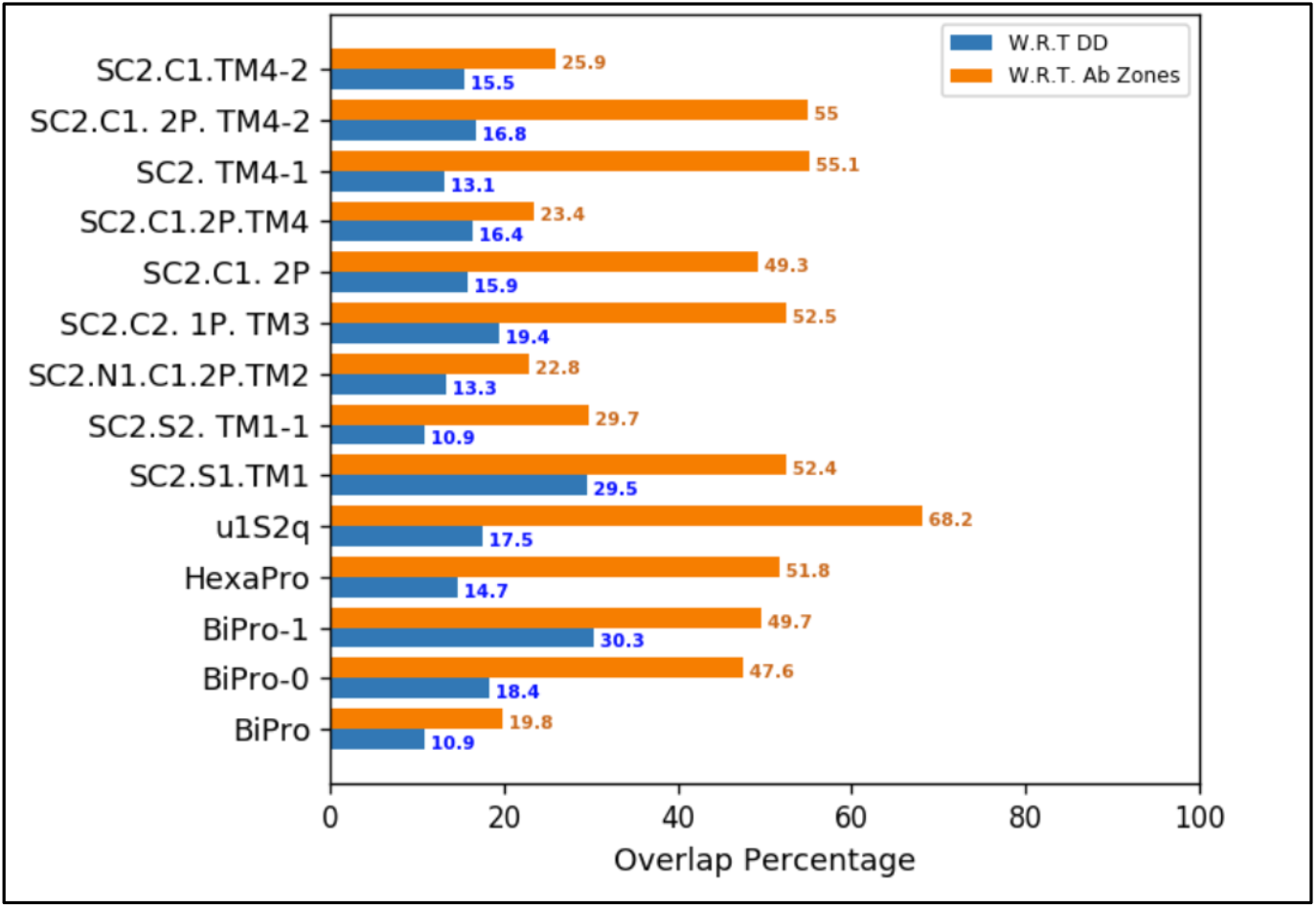
Percentage overlap between dynamic domains and antibody binding zones for different spike protein mutants. W.R.T. : “with respect to”. The blue bar indicates how much of the dynamic domain space overlaps with known antibody binding regions. The orange bar indicates the amount of antibody zones overlapping with dynamic domains.

We find that the u1S2q mutant has the highest overlap between dynamic domains and antibody binding zones, at 68.2% and 17.5% overlaps, respectively (**Fig. 8**). These values indicate that the protein dynamics result in high exploration of antibody binding zones, and simultaneously regions outside of these zones. Based on this result, we hypothesize that 2 RBD up S proteins garner increased neutralizing activity as compared to, for example, single RBD up S proteins. Results from the dynamics analysis of the 2 RBD up structures (**Fig. 1**,**3**) suggest that one RBD is highly dynamic and retains a higher propensity to flip between open and closed orientations, while the other RBD is less mobile. Dynamic domains are also present at all three NTDs. Based on our findings and results of experimental studies (15), the flexibility of the NTDs is essential for the structure to accommodate 2 up RBDs. Multi-RBD up structures expose a greater number of epitope zones. Therefore, the existence of additional dynamic domains around these sites may confer sufficient flexibility to allow for several possible antibodies binding at the same time. We note that alternatively, excessive mobility of these RBDs may hinder antibody binding and destabilize critical bonds. We expect that future design of S protein mutants will benefit from these considerations.

Based on our observations of 1 RBD up S protein dynamics and analysis of the percentage overlap results, we find that structures which present dynamic domains covering a large surface area in the S1 subunit are more likely to expose a variety of epitope zones and possess the mobility needed for conformational change in response to antibody binding. Out of the structures we have analyzed, u1S2q, BiPro, HexaPro, BiPro-1, and SC2.C2.1P.TM3 are predicted to elicit the most varied and neutralizing antibody response while remaining stabilized at the S2 subunit. The SC2.S2.TM1-1 S protein presents dynamic domains that cover significant surface area and a low percentage overlap with respect to dynamic domains, indicating varied antibody response as well. However, this structure also presents dynamic domains in the S2 subunit. These dynamic domains may represent an additional region for antibody targeting but we predict that this instability can also increase virus-cell fusion efficacy through significant structural shifts. The SC2.S2.TM1-1 S protein may be a candidate for a cocktail of antibodies that target both S1 and S2 subunits in the manner that is described by Chi et al. (12) and Pinto et al. (41).

### 3.6. Antibodies bound to SARS-CoV-2 S protein influence local domain dynamics

We next constructed ANMs for SARS-CoV-2 S proteins bound to neutralizing antibodies in each of the defined zones and found their associated modal trajectories. Dynamics analysis was performed to characterize neutralizing mechanisms further and determine how antibody contacts may influence S protein stability. The starting PDB structures used for analysis are PDB IDs 7K4N (2), 7BYR (3), 7K43 (2), 7JW0 (5), and 7C2L (12), and include antibodies cover all zones (**Fig. 8**). We note that the structural and dynamical properties associated with each zone may not hold for all antibodies that bind to that zone. We also compute deformation and solvent accessibility profiles for each antibody bound S protein as in Section 3.2, to highlight new dynamical patterns associated with antibody bound structures (**Fig. 9**).

**Figure 9:**
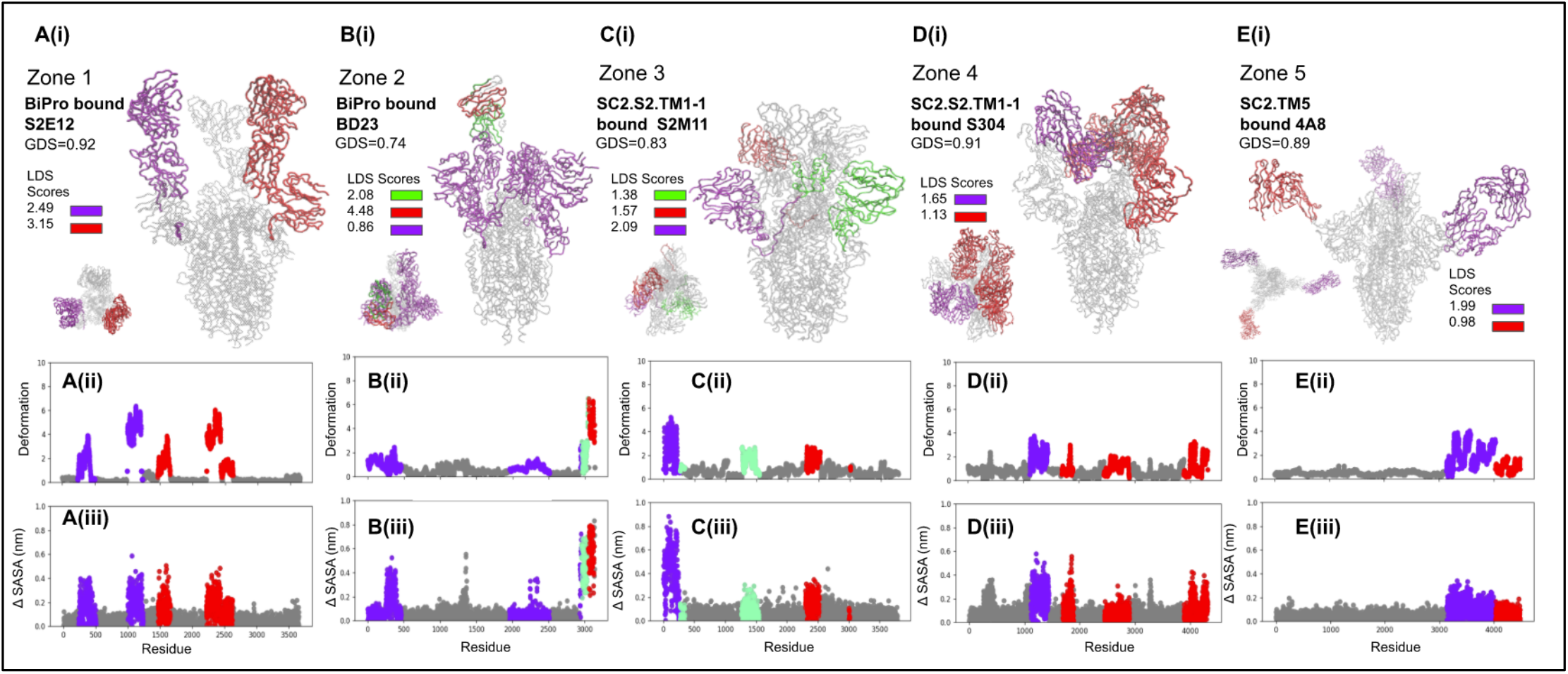
Results of dynamic domain analysis for antibody bound S proteins with PBD IDs **(A)** 7K4N (2), **(B)** 7BYR (3), **(C)** 7K43 (2), **(D)** 7JW0 (5), and **(E)** 7C2L (12). The PDB ID, global dynamics score (GDS), local dynamics scores (LDSs), deformation profile **(ii)**, and Δ SASA profile **(iii)** is listed for each structure.

The SARS-CoV-2 BiPro S protein mutant binds to antibody S2E12 in zone 1 (PDB ID 7K4N) (2). This combination leads to highly potent neutralizing action by the antibody overlapping with the ACE2 binding site (2). As with most zone 1 antibodies, S2E12 can only bind to the S protein when its RBD is in the up conformation (2). ELISA assays confirm this, showing solutions containing closed trimer structures with very low neutralizing action compared to solutions containing RBD up S proteins (2). The BiPro sequence displays a propensity to reorganize into a multi-RBD up structure upon binding, as confirmed by our analysis of its local domain dynamics (**Fig. 3A**) (2,7). Such reorganization is believed to take place in the BiPro protein, enabling it to adopt a 3 RBD up conformation where each RBD is bound to an antibody (2). Dynamic domain analysis of BiPro bound to S2E12 ANM (**Fig. 9A**) reveals dynamic domains around two of the up RBDs bound to antibodies, while the third is stabilized. Thus, as expected, antibodies provide some structural stabilization to the S protein in addition to neutralization by ACE2 competition. LDS and GDS scores for this structure are indeed lower than that of BiPro—pointing to S protein stabilization. Cryo-EM characterization of the complex confirms that antibody binding has a stabilization effect noting that it dramatically improves protein resolution (2).

The BD23 antibody binds to the BiPro S protein mutant in zone 2 (PDB ID 7BYR) (3,7). The potent neutralizing action provided by BD23 is through direct overlap with the RBM on one RBD and overlap with an opposite RBD—blocking ACE2 binding on both RBDs. BD23, like many antibodies that bind in zone 2, can bind to the RBD in the up or down position. Dynamic domain analysis of the ANM model of BD23 bound to BiPro (**Fig. 9B**) reveals a slightly mobile S1 domain (LDS=0.84 compared to GDS=0.74). The BD23 protein is determined to have 2 highly mobile domains (LDS=4.48 and LDS=2.08). The NMA trajectory (see **7BYR Supplemental Video**) shows that BD23 stabilizes the RBD and fluctuates in a hinge-like motion, enabling it to interact with the neighboring closed RBDs. In experimental studies, BD23 is suspected to act similarly to S2E12 to stabilize S1 subunit domains while providing neutralizing activity (3). Thus, zone 2 presents a prime antibody target since it is accessible in both open and closed states but can still provide potent neutralization through ACE2 competition and structural stabilization.

The S2M11 antibody binds to the SC2.S2.TM1-1 at all three RBDs through a quaternary epitope in zone 3 (PDB ID 7K43) (2,6). Cryo-EM data shows that S2M11 can achieve its peak neutralizing ability when attaching to 1 RBD up structures due to its ability to make contacts with zone 4 as the RBD changes position (2). Upon reaching the closed position, S2M11 can lock down all RBDs while burying the RBM and paratope (2). For the SC2.S2.TM1-1 structure, the fully closed conformation may be more accessible since most of the S1 subunit region is stabilized with two of the three RBDs in the down conformation (**Fig. 4H**) (6). The domain dynamics for the ANM model of S2M11 and S protein complex reveal dynamic domains around two of the NTDs and one of the S2M11 antibodies (**Fig. 9C**). In the case of the two NTDs, these may need to adopt enough mobility to compensate for the locked position of the RBDs. Cryo-EM observed states also show that S2M11 antibodies bridge neighboring RBDs (2). The dynamic domain identified around one of the antibodies (**Fig. 9C)** may indicate weaker binding to neighboring RBDs.

The SC2.S2.TM1-1 S protein also binds to the S304 antibody (PDB ID 7K43) in zone 4 and makes contacts at all RBDs which reorganize into the 3 RBD up conformation (5). This combination leads to weak neutralization through partial ACE2 competition via steric clashing (5). The exact binding mechanisms for S304 are unclear, but based on the behavior of the other zone 4 antibody, S2A4, S304 may act like a molecular ratchet to wedge open RBDs (5). The NMA trajectory of the SC2.S2.TM1-1 and S304 model shows dynamic domains covering large portions of the S1 subunit and antibody (**Fig. 9D**). The larger and less mobile (LDS=1.13) dynamic domain (**Fig. 9D (i)**) comprises two of the up RBDs and all three antibody fragments, and the dominant domain (purple) fluctuates independently of the rest of the structure (see **7K43 Supplemental Video**). This suggests that binding to this zone 4 epitope may not stabilize the RBD position in all cases. This fluctuation may provide a means to bind to ACE2 and evade steric clashes with S304.

Lastly, S proteins also bind to the antibody 4AD in zone 5 (PDB ID 72CL) at the tip of each NTD (12) Thus, the potent neutralizing action provided by this epitope involves no direct contact with ACE2 or the RBDs (12). Cryo-EM analysis shows that the 4AD association stabilizes the NTDs, quantified by dramatically improved structural resolution (12). Analysis of the S protein bound to 4AD modal trajectory confirms this observation. It further shows stabilization of the whole S protein in the 1 RBD up confirmation and dynamic domains covering the 4AD antibody (**Fig. 9E**). This highlights a possible neutralization mechanism since ACE2 is not able to bind to the S protein even in the presence of an up RBD. The stabilized RBD and hindered flexibility of surrounding regions may prevent it from fitting onto ACE2 and structurally reorganizing in response to binding. This is an example of the neutralizing potential of antibodies through indirect structural stabilization mechanisms rather than ACE2 competition. Additionally, this result suggests that the NTD is a key area to target when designing a structurally stabilized prefusion S protein. Presented findings in our study and in others demonstrate that increased stability can be achieved by making mutations at the S1/S2 junction and hinge region that controls RBD fluctuation. Stabilized prefusion structures that prevent RBD fluctuation may have increased immunogenic properties by inhibiting ACE2 binding mechanisms.

### 3.7. Cellular fusion and receptor binding mechanisms differ across SARS and MERS family coronaviruses

#### Cleavage sites inform differing cellular fusion properties

SARS and MERS family coronaviruses are similar in their viral architecture but can be widely varied in sequence and biological virus effects (6,84). A key difference between SARS-CoV-2 and other coronaviruses is its widespread and pernicious nature—a key characteristic believed to be rooted in the S protein’s function (84). While S proteins from SARS-CoV-2, SARS-CoV, and MERS-CoV all adopt the characteristic trimer structure, they differ in sequence, specifically, at their cleavage sites (32). Cleavage of S proteins is required for fusion of viral and cell membranes. Cleavage site sequence encodes which cell receptors can be recognized by the S protein and can therefore modulate cell-virus fusion efficacy (32,83). SARS family S proteins recognize ACE2 cell surface receptors, whereas the MERS-CoV S protein recognizes DPP4 (12,19,83,85). The combination of R|SV and R|SF cleavage sites within the SARS-CoV-2 S protein is unique to SARS-CoV-2 S proteins in comparison to other coronavirus proteins (see **Table 2**) and is suspected to render SARS-CoV-2 WT S proteins highly recognizable by furin, an abundant enzyme in respiratory environments (32). Unlike SARS-CoV-2, SARS-CoV contains an R|ST S1/S2 cleavage site and MERS-CoV contains an R|SA cleavage site at S2’, which may provide an additional immune mechanism that accounts for their less-widespread nature.

**Table 2:** Comparison of SARS and MERS cleavage sites and sequence similarity to SARS-CoV-2.

| Virus | S1/S2 site | S2' site | Similarity with SARS-CoV-2 | References |
| --- | --- | --- | --- | --- |
| SARS-CoV-2 | SPRRAR\|SV | KR\|SF | 100% | Coutard et al. (32) |
| SARS-CoV | TVSLLR\|ST | KR\|SF | 73% | Coutard et al. (32), Yuan et al. (10) |
| MERS-CoV | TPRSCR\|SV | AR\|SA | 31% | Coutard et al. (32), Yuan et al. (10), Li et al (85) |

#### Comparison of SARS-CoV-2 and MERS S protein dynamics suggests potential immune escape mechanisms

We also investigate the dynamic and mechanistic differences between SARS-CoV-2 and MERS-CoV S proteins. Specifically, we examine the MC.SD.TM1 MERS-CoV S protein that adopts both 1 RBD up (PDB ID 6×2F) and 2 RBD up (PDB ID 6×2C) conformations (10). This sequence has stabilizing S1/S2 cleavage mutations, NTD clip, and trimerization motif mutations as compared to the Uniprot ID K9N5Q8 MERS-CoV sequence (10). We measure a 94% alignment between the two sequences. Our model of the 1 RBD up structure (PDB ID 6×2F) shows a dominant dynamic domain surrounding the up RBD and auxiliary dynamic domains presented on either side of the RBD (**Fig. 10A**). This breakdown is similar to the dynamic patterns seen in SARS-CoV-2 models (**Fig. 3-5**). The 2 RBD up model of MC.SD.TM1 (PDB ID 6×2C) presents dynamic domains surrounding both up RBDs and their adjacent NTDs (**Fig. 10B**) in the same manner as u1s2q. Also the dynamic domain signatures of the 2 RBD up MC.SD.TM1 MERS-CoV S protein model and 2 RBD up u1S2q SARS-CoV-2 S protein model (**Fig. 10B, Fig. 3C**) resemble one another and the only divergence between them is in an additional dynamic domain around MC.SD.TM1’s down RBD. Interestingly, the MERS-CoV 2 RBD up configuration exhibits these behaviors without the presence of the A570L, T572I, F855Y, and N856I mutations observed in the SARS-CoV-2 u1S2q sequence. These dynamics in the MERS-CoV S protein suggest that it may be naturally more flexible in the S1 domain compared to SARS-CoV-2. Additionally, the increased solvent exposure and level of RBD fluctuation presented by MERS-CoV S proteins with up RBDs may increase neutralizing antibody activity, as shown by the experimental immunogenic studies on u1S2q (15) . These RBD-linked mechanisms likely contribute to MERS-CoV localized spread in contrast to the SARS-CoV-2 global spread.

**Figure 10:**
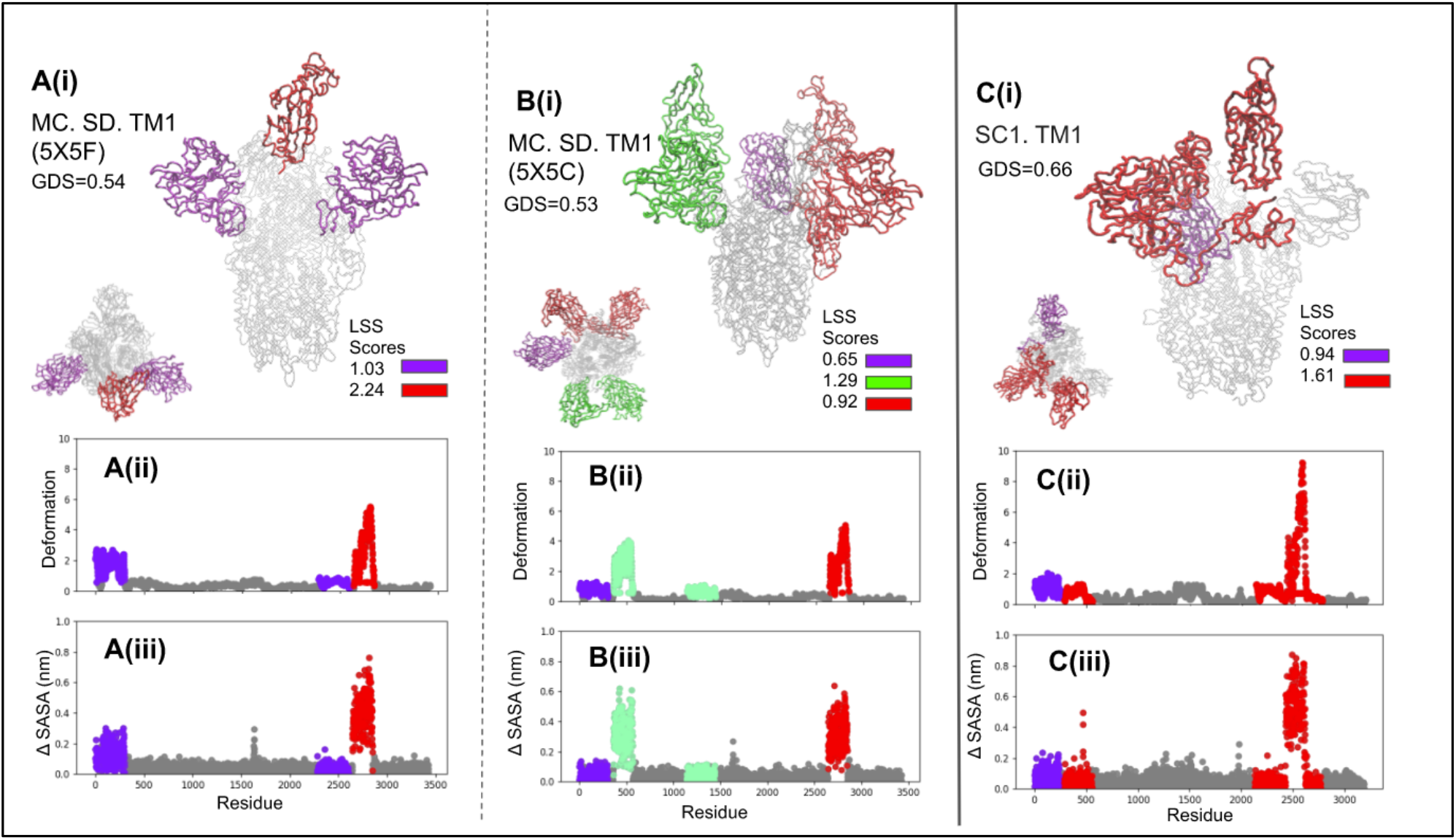
Comparison of dynamic domain analysis results for 1 RBD up MERS-CoV (10) **(A)** 2 RBD up MERS-CoV (10) **(B)** and 1 RBD up SARS-CoV (18) **(C)** S proteins. The PDB ID, global dynamics score (GDS), local dynamics scores (LDSs), deformation profile **(ii)**, and Δ SASA profile **(iii)** is listed for each structure. Dynamic domains are highlighted in different colors.

#### Comparison of SARS-CoV and SARS-CoV-2 S protein dynamics suggests potential immune escape mechanism

We characterize the structural and dynamical differences between SARS-CoV-2 and SARS-CoV by constructing anisotropic network models and performing dynamic domain analysis on resulting NMA trajectories (**Fig. 10C, Fig. 3-5**). Multi RBD up prefusion SARS-CoV S proteins are not currently available in the PDB; therefore we analyze the 1 RBD up structure, SC1.TM1 (PDB ID 6ACD) (18). This S protein contains an S1/S2 alanine mutation compared to the WT, NCBI Accession NP_828851.1 (18). An earlier study shows that this mutation does not significantly affect structural orientation but does impact ACE2 fusion properties (18). Dynamics analysis of SC1.TM1 ANM (**Fig. 10C**) shows a dominant dynamic domain around the up RBD and adjacent NTD, and another auxiliary domain opposite to the up RBD. This breakdown is similar to the dynamic domain patterns seen in SARS-CoV-2 models (**Fig. 3-5**). However, the maximum deformation for SC1.TM1 ANM is approximately 10 Å around its RBD, whereas the maximum deformation for 1 RBD up SARS-CoV-2 S protein mutants typically ranges between 6 Å and 8.5 Å. This spurs the hypothesis that there may be an RBD fluctuation window that optimizes its ability to lock onto ACE2. Our results from SARS-CoV-2 antibody binding studies suggest that static, or slightly fluctuating, single up RBDs may hinder ACE2 binding. The results of the SARS-CoV structure demonstrates that a highly fluctuating RBD may also hinder ACE2 binding—possibly providing a reason for its localization.

#### Analysis of SARS and MERS family antibody-bound S proteins informs SARS-CoV-2 antibody targets

Lastly, we constructed ANMs for SARS-CoV and MERS-CoV S proteins bound to neutralizing antibodies and performed dynamic domain analysis on their resulting modal trajectories to see if these presented any key differences to SARS-CoV-2 results. Full antibody bound trimer structures for these coronaviruses are not as prevalent as those associated with SARS-CoV-2, so we performed analysis on MERS-CoV complex (PDB ID 5W9K (1)) and SARS-CoV complex (PDB ID 6NB6 (4)) as case studies (**Fig. 11**).

**Figure 11:**
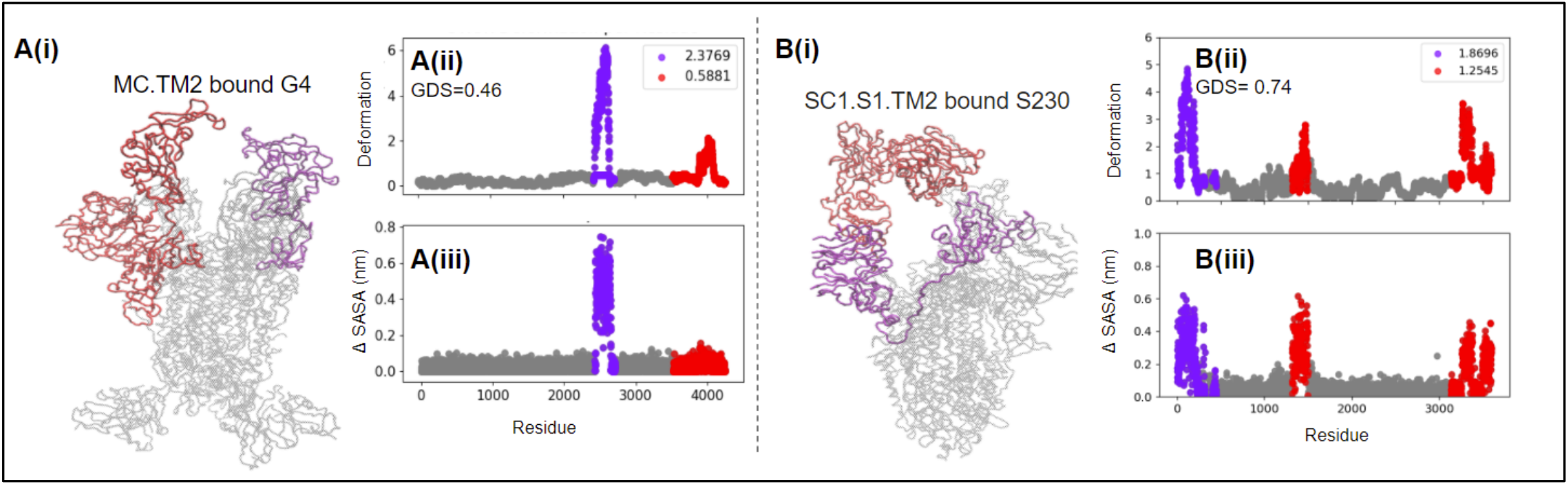
Results of dynamic domain analysis for antibody bound MERS-CoV S protein PDB ID 5W9K (1) **(A)** and SARS-CoV S protein PDB ID 6NB6 (4) **(B)**. The PDB ID, global dynamics score (GDS), local dynamics scores (LDSs), deformation profile **(ii)**, and Δ SASA profile **(iii)** is listed for each structure. Dynamic domains are highlighted in different colors.

MERS-CoV S protein MC.TM1 structure bound to the neutralizing antibody G4 (PDB ID 5W9K) (1) deviates in behavior from all other complexes we have examined because G4 is bound to the S2 domain near the trimer base. The mode of neutralization is extreme stabilization to prevent S2 structural reorganization upon receptor binding and thus prevention of cellular fusion (39). Based on the dynamic patterns of the G4 bound MC.TM1 model (**Fig. 11A**), the S2 domain of MC.TM1 and connecting antibody regions are structurally stabilized. Dynamic domains exist in the S1 subunit around the up RBDs. The G4 antibody may not only stabilize local dynamics, but global oscillations as well. From our analysis of SC2.TM4-1 and the SARS-CoV-2 WT structure, we find that the portion of the trimerization motif and C-terminal domain (CTD) that makes contact with the virion surface may contribute to global rocking motions experienced by the trimer around a hinge site. The MERS-CoV trimerization motif may generate a similar effect on global motions. Even without a resolved trimerization motif, MC.TM1 does present the lowest GDS score seen among all other structures considered here. The neutralization capacity of S2 binding MERS-CoV antibodies presents another mechanism that may not yet be available to SARS-CoV-2 and contributes to the localized spread of MERS-CoV. Targeting antibodies that attach to the base of the trimer may be particularly effective for SARS-CoV-2 S proteins that have a propensity to exhibit unstable S2 dynamics such as the WT protein or SC2.S2.TM1-1 mutant.

The SARS-CoV S protein SC1.S1.TM2 structure bound to an S230 antibody fragment (PDB ID 6NB6) shows that S230 makes contact with the tip of the up RBD and ratchets open the opposite RBD to create a bridge between the two RBDs (4). The neutralization action is generated by a direct overlap of the ACE2 binding site by the antibody. Dynamics analysis of the S230 bound SC1.S1.TM2 ANM modal trajectory (**Fig. 11B**) reveals dynamic domains around the S230, SC1.S1.TM2 RBD group, and another around the neighboring NTD. Since the RBDs and antibodies form a single dynamic domain, this interaction between the proteins is suspected to be firm and resistant to deformation—providing an auxiliary neutralization mechanism. This trait is also exhibited by SARS-CoV-2 structures (see **Supplementary Table 1**). This result provides further evidence that the antibody binding characteristics of the SARS-CoV S protein are similar to SARS-CoV-2.

## 4. CONCLUSIONS

The results presented herein, synthesized together with prior experimental data, highlight the viability of elastic network modeling, integrated normal mode analysis, thermal stability predictions, and dynamic domain analysis to characterize the structure, dynamics, and associated functions of coronavirus S proteins. We provide a review of the local dynamic patterns and associated implications of common SARS-CoV-2 mutations. We find that alterations in trimerization motifs affect trimer thermal stability and contribute to the overall level of global dynamics experienced by the structure. Changes to the first 32 residues through added signal peptides or deletions of residues may alter the stability of the NTD region by disrupting critical bonds. Indirect destabilization may occur through changes in protein dynamic patterns within domains adjacent to critical bond sites and the S1/S2 junction. We show that SARS-CoV-2 S proteins may also be structurally and dynamically sensitive to S1/S2 furin cleavage mutations, which have the potential to augment the level and distribution of force transmitted to either subunit by the oscillation of the RBD. For example, the GAGS mutation may provide a mild stabilizing effect to the S2 domain and restrict dynamic domains to the S1 subunit—although this may not always be the case if destabilizing NTD mutations are present. Proline mutations in the S2 subunit may not dramatically affect local dynamics, although they may help to partially stabilize S1 subunit dynamic regions by providing a more heterogeneous force distribution across the structure. Our thermal stability predictions, however, did show that proline mutations improve thermal stability, as expected. We also found that thermal stability may be improved through specific furin cleavage mutations. Additionally, mutations that support multi RBD up structures—such as those introduced in the u1S2q sequence—may decrease thermal stability, so these structures may benefit from additional proline mutations, for example.

We synthesize available experimental SARS-CoV-2 antibody binding data to create a SARS-CoV-2 antigenic map and label known antibody binding zones. By comparing local dynamics of S protein mutants we find that it is possible, using our models, to predict the accessibility of known epitope zones and thereby predict the binding properties of SARS-CoV-2 S protein mutants. This can be valuable information for determining which S protein variants to use for immunogen design. Thus far, we predict u1S2q, BiPro, HexaPro, BiPro-1, and SC2.C2.1P.TM3 to elicit the most varied antibody response. We presented case studies of SARS-CoV-2 trimers bound to antibodies in each zone, showing that antibodies affect protein dynamics which can influence mechanisms of neutralization. Some directly overlap with ACE2 binding sites (only accessed in the RBD up conformation): in this case there is a direct neutralizing mechanism. When antibodies are bound to other zones, they can block ACE2 binding directly and/or induce dynamic perturbations that shift S proteins into a neutralizing configuration. Alternatively, antibodies can also initiate neutralizing conformational changes such as bridging a multi RBD up structure or stabilizing NTD regions to prevent RBD-ACE2 binding shifts. Overall, our models can predict new regions of viable epitopes as these show a strong correlation with the location of dynamic domains.

We also presented analysis of free SARS-CoV and MERS-CoV trimers and trimers bound to antibodies. We found dynamic mechanisms through which multi RBD up structures may impair binding to cell receptors and elicit a more varied antibody response. MERS-CoV S proteins show a higher propensity to adopt multi RBD up structures than SARS-CoV-2 S proteins, which may account for their more limited and localized infection. SARS-CoV S proteins do not adopt multi-RBD up conformations as frequently but their RBD fluctuations may induce larger deformations than in SARS-CoV-2 S proteins, which may impair their ability to lock on to cell receptors. Additionally, the S1/S2 junction for SARS-CoV proteins presents a different furin cleavage motif than MERS-CoV and SARS-CoV-2 proteins which further impairs cellular fusion. By comparing local domain dynamics associated with all S proteins, we found that the arginine residues surrounding the furin cleavage site in SARS-CoV-2 WT S protein may account for the mechanism for S2 subunit destabilization. S2 subunit destabilization and the presence of a dynamic domain in the S2 subunit point to significant structural shifts upon binding and, thus, better virus-cell fusion for the SARS-CoV-2 S protein.

Dynamics analysis of SARS-CoV and MERS-CoV trimers bound to antibodies suggests that the antibody binding mechanisms of SARS-CoV S proteins may not differ significantly from SARS-CoV-2 S proteins. The MERS-CoV S protein structure presents a unique S2 epitope not seen in cryo-EM studies of SARS-CoV-2. The G4 antibody binds to the base of the trimer close to the start of the trimerization motif. This antibody provides strong S2 stabilization and is suspected to decrease the level of global dynamics by stabilizing the trimer against the virion surface. Based on these results, we suspect that S2 subunit targeting antibodies may present strong neutralizing and stabilizing effects in the SARS-CoV-2 S protein as well. Extensive studies have not been performed on S2 targeting antibodies for SARS-CoV-2, so this could be a new avenue for research (86).

Our recommendation for the molecular design of a SARS-CoV-2 S protein immunogen is to create a multi RBD up structure whose RBD dynamics fall on either end of a spectrum of flexibility: either significant fluctuation or by contrast, enhanced stability. The latter may be more easily designed. Key mutations for creating such a structure would be: GSAS furin cleavage site; 6P in S2; A570L, T572I, F855Y, N856I; and stabilizing the trimerization motif. This design is predicted to increase the number of accessible neutralizing epitopes, stabilize S2 subunits, enhance the multi RBD up structure, and increase thermal stability. Increased proline mutations may also inhibit structural reorganization upon ACE2 binding and more evenly distribute any dynamic domains presented in the S1 subunit. The addition of a signal peptide may create an additional bridge near the S1/S2 junction and promote stabilization of NTDs and S2 subunits. However, it is unclear if this will improve antibody binding properties or render the structure unable to adjust to antibodies that require conformational change.

Methods for quick characterization of coronavirus systems are essential to investigate future mutations of the SARS-CoV-2 virus and other coronavirus infectious disease outbreaks, and to develop treatment and prevention options. Our study proposes an integrated framework to characterize such proteins and evaluate their functional regions. The findings presented in this study can be used to further SARS-CoV-2 immunogen design and vaccine applications. SARS-CoV-2 is mutating at an alarming rate and unique variants are emerging in different regions of the world. For example, a new common variant has emerged in the UK that is becoming progressively prevalent in other areas of the world (87,88). Thus, future directions of this work include the comparison of emerging SARS-CoV-2 S protein variants to predict their effects and recommend molecular design solutions.

## ACKNOWLEDGEMENTS

This work was supported by the University of Connecticut OVPR Covid-19 Rapid Start Funding Program. This work utilized the Extreme Science and Engineering Discovery Environment (XSEDE), which is supported by National Science Foundation grant number ACI-1548562 (83). XSEDE resources Stampede 2 and Ranch at the Texas Advanced Computing Center and Bridges at the Pittsburg Supercomputing Center through allocation TG-MCB180008 were used.

## SUPPORTING CITATIONS

References (2,3,6-9,12-17,20,21,28,35,41,78,81,83,89-100) are also listed in the Supplementary Information

