## Supplementary Information for "Modeling Coronavirus Spike Protein Dynamics: Implications for Immunogenicity and Immune Escape"

### 1. SARS-CoV-2 Protein Sequences

*Mutations are in yellow
** Spike protein names correspond to what is listed in the literature through experimental studies or by database identifier. If no name exists, then the spike protein mutant is assigned a name. For proteins that are assigned a name, the naming conventions of its associated experimental study are used. If this information is not listed, then different families of mutations are separated by a period and mutation is assigned a numerical identifier. For example, the 6VXX SARS-CoV-2 mutant sequence contains a signal peptide and trimerization motif, thus its name convention is SC2.S1.TM1. Should another spike protein have the same, for example, trimerization motif mutation, it will also have the ".TM1" identifier. Similar families of mutations include dashes.

**Uniport P0DTC2 (Wild Type Sequence) (1)**
MFVFLVLLPLVSSQCVNLTTRTQLPPAYTNSFTRGVYYPDKVFRSSVLHSTQDLFLPFFSNVTWFH
AIHVSGTNGTKRFDNPVLPFNDGVYFASTEKSNIIRGWIFGTTLDSKTQSLLIVNNATNVVIKVCEFQFCNDP
FLGVYYHKNNKSWMESEFRVYSSANNCTFEYVSQPFLMDLEGKQGNFKNLREFVFKNIDGYFKIYSKHTPI
NLVRDLPQGFSALEPLVDLPIGINITRFQTLLALHRSYLTPGDSSSGWTAGAAAYYVGYLQPRTFLLKYNEN
GTITDAVDCALDPLSETKCTLKSFTVEKGIYQTSNFRVQPTESIVRFPNITNLCPFGEVFNATRFASVYAWNR
KRISNCVADYSVLYNSASFSTFKCYGVSPTKLNDLCFTNVYADSFVIRGDEVRQIAPGQTGKIADYNYKLPD
DFTGCVIAWNSNNLDSKVGGNYNYLYRLFRKSNLKPFERDISTEIYQAGSTPCNGVEGFNCYFPLQSYGFQ
PTNGVGYQPYRVVVLSFELLHAPATVCGPKKSTNLVKNKCVNFNFNGLTGTGVLTESNKKFLPFQQFGRDI
ADTTDAVRDPQTLEILDITPCSFGGVSVITPGTNTSNQVAVLYQDVNCTEVPVAIHADQLTPTWRVYSTGSN
VFQTRAGCLIGAEHVNNSYECDIPIGAGICASYQTQTNSPRRARSVASQSIIAYTMSLGAENSVAYSNNSIAIP
TNFTISVTTEILPVSMTKTSVDCTMYICGDSTECSNLLLQYGSFCTQLNRALTGIAVEQDKNTQEVFAQVKQ
IYKTPPIKDFGGFNFSQILPDPSKPSKRSFIEDLLFNKVTLADAGFIKQYGDCLGDIAARDLICAQKFNGLTVL
PPLLTDEMIAQYTSALLAGTITSGWTFGAGAALQIPFAMQMAYRFNGIGVTQNVLYENQKLIANQFNSAIG
KIQDSLSSTASALGKLQDVVNQNAQALNTLVKQLSSNFGAISSVLNDILSRLDKVEAEVQIDRLITGRLQSL
QTYVTQQLIRAAEIRASANLAATKMSECVLGQSKRVDFCGKGYHLMSFPQSAPHGVVFLHVTYVPAQEKN
FTTAPAICHDGKAHFPREGVFVSNGTHWFVTQRNFYEPQIITTDNTFVSGNCDVVIGIVNNTVYDPLQPELD
SFKEELDKYFKNHTSPDVDLGDISGINASVVNIQKEIDRLNEVAKNLNESLIDLQELGKYEQYIKWPWYIWL
GFIAGLIAIVMVTIMLCCMTSCCSCLKGCCSCGSCCKFDEDDSEPVLKGVKLHYT

**Name**: BiPro
**PDB:** 6VSB (2)
MFVFLVLLPLVSSQCVNLTTRTQLPPAYTNSFTRGVYYPDKVFRSSVLHSTQDLFLPFFSNVTWFHAIHVSG
TNGTKRFDNPVLPFNDGVYFASTEKSNIIRGWIFGTTLDSKTQSLLIVNNATNVVIKVCEFQFCNDPFLGVY

YHKNNKSWMESEFRVYSSANNCTFEYVSQPFLMDLEGKQGNFKNLREFVFKNIDGYFKIYSKHTPINLVRD
LPQGFSALEPLVDLPIGINITRFQTLLALHRSYLTPGDSSSGWTAGAAAYYVGYLQPRTFLLKYNENGTITD
AVDCALDPLSETKCTLKSFTVEKGIYQTSNFRVQPTESIVRFPNITNLCPFGEVFNATRFASVYAWNRKRISN
CVADYSVLYNSASFSTFKCYGVSPTKLNDLCFTNVYADSFVIRGDEVRQIAPGQTGKIADYNYKLPDDFTG
CVIAWNSNNLDSKVGGNYNYLYRLFRKSNLKPFERDISTEIYQAGSTPCNGVEGFNCYFPLQSYGFQPTNG
VGYQPYRVVVLSFELLHAPATVCGPKKSTNLVKNKCVNFNFNGLTGTGVLTESNKKFLPFQQFGRDIADTT
DAVRDPQTLEILDITPCSFGGVSVITPGTNTSNQVAVLYQDVNCTEVPVAIHADQLTPTWRVYSTGSNVFQT
RAGCLIGAEHVNNSYECDIPIGAGICASYQTQTNSPGSASSVASQSIIAYTMSLGAENSVAYSNNSIAIPTNFT
ISVTTEILPVSMTKTSVDCTMYICGDSTECSNLLLQYGSFCTQLNRALTGIAVEQDKNTQEVFAQVKQIYKT
PPIKDFGGFNFSQILPDPSKPSKRSFIEDLLFNKVTLADAGFIKQYGDCLGDIAARDLICAQKFNGLTVLPPLL
TDEMIAQYTSALLAGTITSGWTFGAGAALQIPFAMQMAYRFNGIGVTQNVLYENQKLIANQFNSAIGKIQD
SLSSTASALGKLQDVVNQNAQALNTLVKQLSSNFGAISSVLNDILSRLDPPEAEVQIDRLITGRLQSLQTYVT
QQLIRAAEIRASANLAATKMSECVLGQSKRVDFCGKGYHLMSFPQSAPHGVVFLHVTYVPAQEKNFTTAP
AICHDGKAHFPREGVFVSNGTHWFVTQRNFYEPQIITTDNTFVSGNCDVVIGIVNNTVYDPLQPELDSFKEE
LDKYFKNHTSPDVDLGDISGINASVVNIQKEIDRLNEVAKNLNESLIDLQELGKYEQGSGYIPEAPRDGQAY
VRKDGEWVLLSTFLGRSLEVLFQGPGHHHHHHHHSAWSHPQFEKGGGSGGGGSGGSAWSHPQFEK

Unresolved: 1-26, 67-78, 96-98, 143-155,177-186, 247-260, 330-334, 444-490, 501-502, 621-640, 673-686, 812-814, 829-850, 1147-1288

**Name:** HexaPro
**PDB:** 6XKL (3)
MFVFLVLLPLVSSQCVNLTTRTQLPPAYTNSFTRGVYYPDKVFRSSVLHSTQDLFLPFFSNVTWFHAIHVSG
TNGTKRFDNPVLPFNDGVYFASTEKSNIIRGWIFGTTLDSKTQSLLIVNNATNVVIKVCEFQFCNDPFLGVY
YHKNNKSWMESEFRVYSSANNCTFEYVSQPFLMDLEGKQGNFKNLREFVFKNIDGYFKIYSKHTPINLVRD
LPQGFSALEPLVDLPIGINITRFQTLLALHRSYLTPGDSSSGWTAGAAAYYVGYLQPRTFLLKYNENGTITD
AVDCALDPLSETKCTLKSFTVEKGIYQTSNFRVQPTESIVRFPNITNLCPFGEVFNATRFASVYAWNRKRISN
CVADYSVLYNSASFSTFKCYGVSPTKLNDLCFTNVYADSFVIRGDEVRQIAPGQTGKIADYNYKLPDDFTG
CVIAWNSNNLDSKVGGNYNYLYRLFRKSNLKPFERDISTEIYQAGSTPCNGVEGFNCYFPLQSYGFQPTNG
VGYQPYRVVVLSFELLHAPATVCGPKKSTNLVKNKCVNFNFNGLTGTGVLTESNKKFLPFQQFGRDIADTT
DAVRDPQTLEILDITPCSFGGVSVITPGTNTSNQVAVLYQDVNCTEVPVAIHADQLTPTWRVYSTGSNVFQT
RAGCLIGAEHVNNSYECDIPIGAGICASYQTQTNSPGSASSVASQSIIAYTMSLGAENSVAYSNNSIAIPTNFT
ISVTTEILPVSMTKTSVDCTMYICGDSTECSNLLLQYGSFCTQLNRALTGIAVEQDKNTQEVFAQVKQIYKT
PPIKDFGGFNFSQILPDPSKPSKRSPIEDLLFNKVTLADAGFIKQYGDCLGDIAARDLICAQKFNGLTVLPPLL
TDEMIAQYTSALLAGTITSGWTFGAGPALQIPFPMQMAYRFNGIGVTQNVLYENQKLIANQFNSAIGKIQDS
LSSTPSALGKLQDVVNQNAQALNTLVKQLSSNFGAISSVLNDILSRLDPPEAEVQIDRLITGRLQSLQTYVTQ
QLIRAAEIRASANLAATKMSECVLGQSKRVDFCGKGYHLMSFPQSAPHGVVFLHVTYVPAQEKNFTTAPAI
CHDGKAHFPREGVFVSNGTHWFVTQRNFYEPQIITTDNTFVSGNCDVVIGIVNNTVYDPLQPELDSFKEELD
KYFKNHTSPDVDLGDISGINASVVNIQKEIDRLNEVAKNLNESLIDLQELGKYEQGSGYIPEAPRDGQAYVR
KDGEWVLLSTFLGRSLEVLFQGPGHHHHHHHHSAWSHPQFEKGGGSGGGGSGGSAWSHPQFEK

Unresolved: 1-26, 67-78, 96-98, 143-155,177-186, 247-260, 330-334, 444-490, 501-502, 621-640, 673-686, 812-814, 829-850, 1147-1288

**Name**: BiPro-1
**PDB:**6Z97 (4)
MFVFLVLLPLVSSQCVNLTTRTQLPPAYTNSFTRGVYYPDKVFRSSVLHSTQDLFLPFFSNVTWFHAIHVSG
TNGTKRFDNPVLPFNDGVYFASTEKSNIIRGWIFGTTLDSKTQSLLIVNNATNVVIKVCEFQFCNDPFLGVY
YHKNNKSWMESEFRVYSSANNCTFEYVSQPFLMDLEGKQGNFKNLREFVFKNIDGYFKIYSKHTPINLVRD
LPQGFSALEPLVDLPIGINITRFQTLLALHRSYLTPGDSSSGWTAGAAAYYVGYLQPRTFLLKYNENGTITD
AVDCALDPLSETKCTLKSFTVEKGIYQTSNFRVQPTESIVRFPNITNLCPFGEVFNATRFASVYAWNRKRISN
CVADYSVLYNSASFSTFKCYGVSPTKLNDLCFTNVYADSFVIRGDEVRQIAPGQTGKIADYNYKLPDDFTG
CVIAWNSNNLDSKVGGNYNYLYRLFRKSNLKPFERDISTEIYQAGSTPCNGVEGFNCYFPLQSYGFQPTNG
VGYQPYRVVVLSFELLHAPATVCGPKKSTNLVKNKCVNFNFNGLTGTGVLTESNKKFLPFQQFGRDIADTT
DAVRDPQTLEILDITPCSFGGVSVITPGTNTSNQVAVLYQDVNCTEVPVAIHADQLTPTWRVYSTGSNVFQT
RAGCLIGAEHVNNSYECDIPIGAGICASYQTQTNSPGSASSVASQSIIAYTMSLGAENSVAYSNNSIAIPTNFT

ISVTTEILPVSMTKTSVDCTMYICGDSTECSNLLLQYGSFCTQLNRALTGIAVEQDKNTQEVFAQVKQIYKT
PPIKDFGGFNFSQILPDPSKPSKRSFIEDLLFNKVTLADAGFIKQYGDCLGDIAARDLICAQKFNGLTVLPPLL
TDEMIAQYTSALLAGTITSGWTFGAGAALQIPFAMQMAYRFNGIGVTQNVLYENQKLIANQFNSAIGKIQD
SLSSTASALGKLQDVVNQNAQALNTLVKQLSSNFGAISSVLNDILSRLDPPEAEVQIDRLITGRLQSLQTYVT
QQLIRAAEIRASANLAATKMSECVLGQSKRVDFCGKGYHLMSFPQSAPHGVVFLHVTYVPAQEKNFTTAP
AICHDGKAHFPREGVFVSNGTHWFVTQRNFYEPQIITTDNTFVSGNCDVVIGIVNNTVYDPLQPELDSFKEE
LDKYFKNHTSPDVDLGDISGINASVVNIQKEIDRLNEVAKNLNESLIDLQELGKYEQGSGYIPEAPRDGQAY
VRKDGEWVLLSTFLGRSLEVLFQGPGHHHHHHHHGSAWSHPQFEKGGGSGGGSGGSAWSHPQFEK

Unresolved: 1-26, 70-81,114-115, 144-187, 243-262, 621-640, 677-689, 828-850, 1148-1288

**Name:** SC2.S1.TM1
**PDB:** 6VYB & 6VXX (5)
MGILPSPGMPALLSLVSLLSVLLMGCVAETGTQCVNLTTRTQLPPAYTNSFTRGVYYPDKVFRSSVLHSTQ
DLFLPFFSNVTWFHAIHVSGTNGTKRFDNPVLPFNDGVYFASTEKSNIIRGWIFGTTLDSKTQSLLIVNNATN
VVIKVCEFQFCNDPFLGVYYHKNNKSWMESEFRVYSSANNCTFEYVSQPFLMDLEGKQGNFKNLREFVFK
NIDGYFKIYSKHTPINLVRDLPQGFSALEPLVDLPIGINITRFQTLLALHRSYLTPGDSSSGWTAGAAAYYVG
YLQPRTFLLKYNENGTITDAVDCALDPLSETKCTLKSFTVEKGIYQTSNFRVQPTESIVRFPNITNLCPFGEVF
NATRFASVYAWNRKRISNCVADYSVLYNSASFSTFKCYGVSPTKLNDLCFTNVYADSFVIRGDEVRQIAPG
QTGKIADYNYKLPDDFTGCVIAWNSNNLDSKVGGNYNYLYRLFRKSNLKPFERDISTEIYQAGSTPCNGVE
GFNCYFPLQSYGFQPTNGVGYQPYRVVVLSFELLHAPATVCGPKKSTNLVKNKCVNFNFNGLTGTGVLTE
SNKKFLPFQQFGRDIADTTDAVRDPQTLEILDITPCSFGGVSVITPGTNTSNEVAVLYQDVNCTEVPVAIHAD
QLTPTWRVYSTGSNVFQTRAGCLIGAEHVNNSYECDIPIGAGICASYQTQTNSPSGAGSVASQSIIAYTMSL
GAENSVAYSNNSIAIPTNFTISVTTEILPVSMTKTSVDCTMYICGDSTECSNLLLQYGSFCTQLNRALTGIAVE
QDKNTQEVFAQVKQIYKTPPIKDFGGFNFSQILPDPSKPSKRSFIEDLLFNKVTLADAGFIKQYGDCLGDIAA
RDLICAQKFNGLTVLPPLLTDEMIAQYTSALLAGTITSGWTFGAGAALQIPFAMQMAYRFNGIGVTQNVLY
ENQKLIANQFNSAIGKIQDSLSSTASALGKLQDVVNQNAQALNTLVKQLSSNFGAISSVLNDILSRLDPPEAE
VQIDRLITGRLQSLQTYVTQQLIRAAEIRASANLAATKMSECVLGQSKRVDFCGKGYHLMSFPQSAPHGVV
FLHVTYVPAQEKNFTTAPAICHDGKAHFPREGVFVSNGTHWFVTQRNFYEPQIITTDNTFVSGNCDVVIGIV
NNTVYDPLQPELDSFKEELDKYFKNHTSPDVDLGDISGINASVVNIQKEIDRLNEVAKNLNESLIDLQELGK
YEQYIKGSGRENLYFQGGGGSGYIPEAPRDGQAYVRKDGEWVLLSTFLGHHHHHHHH

Unresolved: 1-26, 70-81, 114-115, 144-185, 243-262, 443-489, 502, 621-640, 677-689, 812, 828-854, 1148-1281

**Name:** SC2.S2. TM1-1
**PDB:** 6ZGG (6)
MGILPSPGMPALLSLVSLLSVLLMGCVAETGMFVFLVLLPLVSSQCVNLTTRTQLPPAYTNSFTRGVYYPD
KVFRSSVLHSTQDLFLPFFSNVTWFHAIHVSGTNGTKRFDNPVLPFNDGVYFASTEKSNIIRGWIFGTTLDSK
TQSLLIVNNATNVVIKVCEFQFCNDPFLGVYYHKNNKSWMESEFRVYSSANNCTFEYVSQPFLMDLEGKQ
GNFKNLREFVFKNIDGYFKIYSKHTPINLVRDLPQGFSALEPLVDLPIGINITRFQTLLALHRSYLTPGDSSSG
WTAGAAAYYVGYLQPRTFLLKYNENGTITDAVDCALDPLSETKCTLKSFTVEKGIYQTSNFRVQPTESIVR
FPNITNLCPFGEVFNATRFASVYAWNRKRISNCVADYSVLYNSASFSTFKCYGVSPTKLNDLCFTNVYADSF
VIRGDEVRQIAPGQTGKIADYNYKLPDDFTGCVIAWNSNNLDSKVGGNYNYLYRLFRKSNLKPFERDISTEI
YQAGSTPCNGVEGFNCYFPLQSYGFQPTNGVGYQPYRVVVLSFELLHAPATVCGPKKSTNLVKNKCVNFN
FNGLTGTGVLTESNKKFLPFQQFGRDIADTTDAVRDPQTLEILDITPCSFGGVSVITPGTNTSNQVAVLYQD
VNCTEVPVAIHADQLTPTWRVYSTGSNVFQTRAGCLIGAEHVNNSYECDIPIGAGICASYQTQTNSPRRARS
VASQSIIAYTMSLGAENSVAYSNNSIAIPTNFTISVTTEILPVSMTKTSVDCTMYICGDSTECSNLLLQYGSFC
TQLNRALTGIAVEQDKNTQEVFAQVKQIYKTPPIKDFGGFNFSQILPDPSKPSKRSFIEDLLFNKVTLADAGFI
KQYGDCLGDIAARDLICAQKFNGLTVLPPLLTDEMIAQYTSALLAGTITSGWTFGAGAALQIPFAMQMAYR
FNGIGVTQNVLYENQKLIANQFNSAIGKIQDSLSSTASALGKLQDVVNQNAQALNTLVKQLSSNFGAISSVL
NDILSRLDPPEAEVQIDRLITGRLQSLQTYVTQQLIRAAEIRASANLAATKMSECVLGQSKRVDFCGKGYHL
MSFPQSAPHGVVFLHVTYVPAQEKNFTTAPAICHDGKAHFPREGVFVSNGTHWFVTQRNFYEPQIITTDNT
FVSGNCDVVIGIVNNTVYDPLQPELDSFKEELDKYFKNHTSPDVDLGDISGINASVVNIQKEIDRLNEVAKN
LNESLIDLQELGKYEQSGRENLYFQGGGGSGYIPEAPRDGQAYVRKDGEWVLLSTFLGHHHHHH

Unresolved: 1-13, 71-75, 618-640, 677-688, 828-848, 941-943, 1147-1287

**Name:** SC2.N1.C1.2P.TM2
**PDB:** 6XF6 (7)
GPQCVNLTTRTQLPPAYTNSFTRGVYYPDKVFRSSVLHSTQDLFLPFFSNVTWFHAIHVSGTNGTKRFDNP
VLPFNDGVYFASTEKSNIIRGWIFGTTLDSKTQSLLIVNNATNVVIKVCEFQFCNDPFLGVYYHKNNKSWM
ESEFRVYSSANNCTFEYVSQPFLMDLEGKQGNFKNLREFVFKNIDGYFKIYSKHTPINLVRDLPQGFSALEP
LVDLPIGINITRFQTLLALHRSYLTPGDSSSGWTAGAAAYYVGYLQPRTFLLKYNENGTITDAVDCALDPLS
ETKCTLKSFTVEKGIYQTSNFRVQPTESIVRFPNITNLCPFGEVFNATRFASVYAWNRKRISNCVADYSVLY
NSASFSTFKCYGVSPTKLNDLCFTNVYADSFVIRGDEVRQIAPGQTGKIADYNYKLPDDFTGCVIAWNSNN
LDSKVGGNYNYLYRLFRKSNLKPFERDISTEIYQAGSTPCNGVEGFNCYFPLQSYGFQPTNGVGYQPYRVV
VLSFELLHAPATVCGPKKSTNLVKNKCVNFNFNGLTGTGVLTESNKKFLPFQQFGRDIADTTDAVRDPQTL
EILDITPCSFGGVSVITPGTNTSNQVAVLYQDVNCTEVPVAIHADQLTPTWRVYSTGSNVFQTRAGCLIGAE
HVNNSYECDIPIGAGICASYQTQTNSPGSASSVASQSIIAYTMSLGAENSVAYSNNSIAIPTNFTISVTTEILPV
SMTKTSVDCTMYICGDSTECSNLLLQYGSFCTQLNRALTGIAVEQDKNTQEVFAQVKQIYKTPPIKDFGGF
NFSQILPDPSKPSKRSFIEDLLFNKVTLADAGFIKQYGDCLGDIAARDLICAQKFNGLTVLPPLLTDEMIAQY
TSALLAGTITSGWTFGAGAALQIPFAMQMAYRFNGIGVTQNVLYENQKLIANQFNSAIGKIQDSLSSTASAL
GKLQDVVNQNAQALNTLVKQLSSNFGAISSVLNDILSRLDPPEAEVQIDRLITGRLQSLQTYVTQQLIRAAEI
RASANLAATKMSECVLGQSKRVDFCGKGYHLMSFPQSAPHGVVFLHVTYVPAQEKNFTTAPAICHDGKA
HFPREGVFVSNGTHWFVTQRNFYEPQIITTDNTFVSGNCDVVIGIVNNTVYDPLQPELDSFKEELDKYFKNH
TSPDVDLGDISGINASVVNIQKEIDRLNEVAKNLNESLIDLQELGKYEQGSGYIPEAPRDGQAYVRKDGEW
VLLSTFLGRSGGGLVPQQSGGLNDIFEAQKIEWHEG

Unresolved: 12-26, 70-81, 114-115, 144-165, 173-185, 243-262, 621-640, 677-690, 828-854, 1148-1266

**Name**: SC2.C2. 1P. TM3
**PDB:** 7AD1 (8)
MFVFLVLLPLVSSQCVNLTTRTQLPPAYTNSFTRGVYYPDKVFRSSVLHSTQDLFLPFFSNVTWFHAIHVSG
TNGTKRFDNPVLPFNDGVYFASTEKSNIIRGWIFGTTLDSKTQSLLIVNNATNVVIKVCEFQFCNDPFLGVY
YHKNNKSWMESEFRVYSSANNCTFEYVSQPFLMDLEGKQGNFKNLREFVFKNIDGYFKIYSKHTPINLVRD
LPQGFSALEPLVDLPIGINITRFQTLLALHRSYLTPGDSSSGWTAGAAAYYVGYLQPRTFLLKYNENGTITD
AVDCALDPLSETKCTLKSFTVEKGIYQTSNFRVQPTESIVRFPNITNLCPFGEVFNATRFASVYAWNRKRISN
CVADYSVLYNSASFSTFKCYGVSPTKLNDLCFTNVYADSFVIRGDEVRQIAPGQTGKIADYNYKLPDDFTG
CVIAWNSNNLDSKVGGNYNYLYRLFRKSNLKPFERDISTEIYQAGSTPCNGVEGFNCYFPLQSYGFQPTNG
VGYQPYRVVVLSFELLHAPATVCGPKKSTNLVKNKCVNFNFNGLTGTGVLTESNKKFLPFQQFGRDIADTT
DAVRDPQTLEILDITPCSFGGVSVITPGTNTSNQVAVLYQNVNCTEVPVAIHADQLTPTWRVYSTGSNVFQT
RAGCLIGAEHVNNSYECDIPIGAGICASYQTQTNSPSRAGSVASQSIIAYTMSLGAENSVAYSNNSIAIPTNFT
ISVTTEILPVSMTKTSVDCTMYICGDSTECSNLLLQYGSFCTQLNRALTGIAVEQDKNTQEVFAQVKQIYKT
PPIKDFGGFNFSQILPDPSKPSKRSFIEDLLFNKVTLADAGFIKQYGDCLGDIAARDLICAQKFNGLTVLPPLL
TDEMIAQYTSALLAGTITSGWTFGAGPALQIPFAMQMAYRFNGIGVTQNVLYENQKLIANQFNSAIGKIQD
SLSSTPSALGKLQDVVNQNAQALNTLVKQLSSNFGAISSVLNDILSRLDKPEAEVQIDRLITGRLQSLQTYVT
QQLIRAAEIRASANLAATKMSECVLGQSKRVDFCGKGYHLMSFPQSAPHGVVFLHVTYVPAQEKNFTTAP
AICHDGKAHFPREGVFVSNGTHWFVTQRNFYEPQIITTDNTFVSGNCDVVIGIVNNTVYDPLQPELDSFKEE
LDKYFKNHTSPDVDLGDISGINASVVNIQKEIDRLNEVAKNLNESLIDLQELGKYEQGSGYIPEAPRDGQAY
VRKDGEWVLLSTFLGRSLEVLFQGPGSLPETGGGSDYKDDDDKGGGGSGGGGSGGGGSGGGGSGGGGSH
HHHHH

Unresolved: 1-26, 70-87, 114-115, 132-165, 173-185, 243-262,443-448, 477-489, 502-503, 621-640, 677-689, 812, 828-854, 1148-1297

**Name:** u1S2q
**PDB:** 6X2B (9)
VNLTTRTQLPPAYTNSFTRGVYYPDKVFRSSVLHSTQDLFLPFFSNVTWFHAIHVSGTNGTKRFDNPVLPFN
DGVYFASTEKSNIIRGWIFGTTLDSKTQSLLIVNNATNVVIKVCEFQFCNDPFLGVYYHKNNKSWMESEFR
VYSSANNCTFEYVSQPFLMDLEGKQGNFKNLREFVFKNIDGYFKIYSKHTPINLVRDLPQGFSALEPLVDLPI
GINITRFQTLLALHRSYLTPGDSSSGWTAGAAAYYVGYLQPRTFLLKYNENGTITDAVDCALDPLSETKCTL
KSFTVEKGIYQTSNFRVQPTESIVRFPNITNLCPFGEVFNATRFASVYAWNRKRISNCVADYSVLYNSASFST

FKCYGVSPTKLNDLCFTNVYADSFVIRGDEVRQIAPGQTGKIADYNYKLPDDFTGCVIAWNSNNLDSKVG
GNYNYLYRLFRKSNLKPFERDISTEIYQAGSTPCNGVEGFNCYFPLQSYGFQPTNGVGYQPYRVVVLSFELL
HAPATVCGPKKSTNLVKNKCVNFNFNGLTGTGVLTESNKKFLPFQQFGRDILDITDAVRDPQTLEILDITPC
SFGGVSVITPGTNTSNEVAVLYQDVNCTEVPVAIHADQLTPTWRVYSTGSNVFQTRAGCLIGAEHVNNSYE
CDIPIGAGICASYQTQTNSPGSASSVASQSIIAYTMSLGAENSVAYSNNSIAIPTNFTISVTTEILPVSMTKTSV
DCTMYICGDSTECSNLLLQYGSFCTQLNRALTGIAVEQDKNTQEVFAQVKQIYKTPPIKDFGGFNFSQILPD
PSKPSKRSFIEDLLFNKVTLADAGFIKQYGDCLGDIAARDLICAQKYIGLTVLPPLLTDEMIAQYTSALLAGT
ITSGWTFGAGAALQIPFAMQMAYRFNGIGVTQNVLYENQKLIANQFNSAIGKIQDSLSSTASALGKLQDVV
NQNAQALNTLVKQLSSNFGAISSVLNDILSRLDPPEAEVQIDRLITGRLQSLQTYVTQQLIRAAEIRASANLA
ATKMSECVLGQSKRVDFCGKGYHLMSFPQSAPHGVVFLHVTYVPAQEKNFTTAPAICHDGKAHFPREGVF
VSNGTHWFVTQRNFYEPQIITTDNTFVSGNCDVVIGIVNNTVYDPLQPELDSFKEELDKYFKNHTSPDVDLG
DISGINASVVNIQKEIDRLNEVAKNLNESLIDLQELGKYEQGSGYIPEAPRDGQAYVRKDGEWVLLSTFLGR
SLEVLFQGPGHHHHHHHHSAWSHPQFEKGGGSGGGGSGGSAWSHPQFEK

Unresolved:  16-26, 68-81, 114-115, 144-185, 243-262, 443-489, 502, 621-640, 677-689, 812, 828-854, 1148-1288

**Name:** SC2**.**C1. 2P
**PDB:** 7CN9 (10)
QCVNLTTRTQLPPAYTNSFTRGVYYPDKVFRSSVLHSTQDLFLPFFSNVTWFHAIHVSGTNGTKRFDNPVLP
FNDGVYFASTEKSNIIRGWIFGTTLDSKTQSLLIVNNATNVVIKVCEFQFCNDPFLGVYYHKNNKSWMESEF
RVYSSANNCTFEYVSQPFLMDLEGKQGNFKNLREFVFKNIDGYFKIYSKHTPINLVRDLPQGFSALEPLVDL
PIGINITRFQTLLALHRSYLTPGDSSSGWTAGAAAYYVGYLQPRTFLLKYNENGTITDAVDCALDPLSETKC
TLKSFTVEKGIYQTSNFRVQPTESIVRFPNITNLCPFGEVFNATRFASVYAWNRKRISNCVADYSVLYNSASF
STFKCYGVSPTKLNDLCFTNVYADSFVIRGDEVRQIAPGQTGKIADYNYKLPDDFTGCVIAWNSNNLDSKV
GGNYNYLYRLFRKSNLKPFERDISTEIYQAGSTPCNGVEGFNCYFPLQSYGFQPTNGVGYQPYRVVVLSFEL
LHAPATVCGPKKSTNLVKNKCVNFNFNGLTGTGVLTESNKKFLPFQQFGRDIADTTDAVRDPQTLEILDITP
CSFGGVSVITPGTNTSNQVAVLYQDVNCTEVPVAIHADQLTPTWRVYSTGSNVFQTRAGCLIGAEHVNNSY
ECDIPIGAGICASYQTQTNSPGSAGSVASQSIIAYTMSLGAENSVAYSNNSIAIPTNFTISVTTEILPVSMTKTS
VDCTMYICGDSTECSNLLLQYGSFCTQLNRALTGIAVEQDKNTQEVFAQVKQIYKTPPIKDFGGFNFSQILP
DPSKPSKRSFIEDLLFNKVTLADAGFIKQYGDCLGDIAARDLICAQKFNGLTVLPPLLTDEMIAQYTSALLA
GTITSGWTFGAGAALQIPFAMQMAYRFNGIGVTQNVLYENQKLIANQFNSAIGKIQDSLSSTASALGKLQD
VVNQNAQALNTLVKQLSSNFGAISSVLNDILSRLDPPEAEVQIDRLITGRLQSLQTYVTQQLIRAAEIRASAN
LAATKMSECVLGQSKRVDFCGKGYHLMSFPQSAPHGVVFLHVTYVPAQEKNFTTAPAICHDGKAHFPREG
VFVSNGTHWFVTQRNFYEPQIITTDNTFVSGNCDVVIGIVNNTVYDP

Unresolved: 14-26, 180-182, 444-489, 622-640, 673-685, 812-852

**Name:** SC2**.**C1.2P.TM4
**PDB:** 6XM0 (11)
QCVNLTTRTQLPPAYTNSFTRGVYYPDKVFRSSVLHSTQDLFLPFFSNVTWFHAIHVSGTNGTKRFDNPVLP
FNDGVYFASTEKSNIIRGWIFGTTLDSKTQSLLIVNNATNVVIKVCEFQFCNDPFLGVYYHKNNKSWMESEF
RVYSSANNCTFEYVSQPFLMDLEGKQGNFKNLREFVFKNIDGYFKIYSKHTPINLVRDLPQGFSALEPLVDL
PIGINITRFQTLLALHRSYLTPGDSSSGWTAGAAAYYVGYLQPRTFLLKYNENGTITDAVDCALDPLSETKC
TLKSFTVEKGIYQTSNFRVQPTESIVRFPNITNLCPFGEVFNATRFASVYAWNRKRISNCVADYSVLYNSASF
STFKCYGVSPTKLNDLCFTNVYADSFVIRGDEVRQIAPGQTGKIADYNYKLPDDFTGCVIAWNSNNLDSKV
GGNYNYLYRLFRKSNLKPFERDISTEIYQAGSTPCNGVEGFNCYFPLQSYGFQPTNGVGYQPYRVVVLSFEL
LHAPATVCGPKKSTNLVKNKCVNFNFNGLTGTGVLTESNKKFLPFQQFGRDIADTTDAVRDPQTLEILDITP
CSFGGVSVITPGTNTSNQVAVLYQDVNCTEVPVAIHADQLTPTWRVYSTGSNVFQTRAGCLIGAEHVNNSY
ECDIPIGAGICASYQTQTNSPGSASSVASQSIIAYTMSLGAENSVAYSNNSIAIPTNFTISVTTEILPVSMTKTS
VDCTMYICGDSTECSNLLLQYGSFCTQLNRALTGIAVEQDKNTQEVFAQVKQIYKTPPIKDFGGFNFSQILP
DPSKPSKRSFIEDLLFNKVTLADAGFIKQYGDCLGDIAARDLICAQKFNGLTVLPPLLTDEMIAQYTSALLA
GTITSGWTFGAGAALQIPFAMQMAYRFNGIGVTQNVLYENQKLIANQFNSAIGKIQDSLSSTASALGKLQD
VVNQNAQALNTLVKQLSSNFGAISSVLNDILSRLDPPEAEVQIDRLITGRLQSLQTYVTQQLIRAAEIRASAN
LAATKMSECVLGQSKRVDFCGKGYHLMSFPQSAPHGVVFLHVTYVPAQEKNFTTAPAICHDGKAHFPREG
VFVSNGTHWFVTQRNFYEPQIITTDNTFVSGNCDVVIGIVNNTVYDPLQPELDSFKEELDKYFKNHTSPDVD

LGDISGINASVVNIQKEIDRLNEVAKNLNESLIDLQELGKYEQGSGYIPEAPRDGQAYVRKDGEWVLLSTFL
GRSLEVLFQGPGHHHHHHHHSAWSHPQFEKGGGSGGGGSGGSAWSHPQFEK

Unresolved: 14-26, 70-79, 144-158, 174-185, 251-263, 445-446, 677-688, 829-848, 1148-1288

**Name:** SC2. TM4-1
**PDB:** 6XR8 (12)
MFVFLVLLPLVSSQCVNLTTRTQLPPAYTNSFTRGVYYPDKVFRSSVLHSTQDLFLPFFSNVTWFHAIHVSG
TNGTKRFDNPVLPFNDGVYFASTEKSNIIRGWIFGTTLDSKTQSLLIVNNATNVVIKVCEFQFCNDPFLGVY
YHKNNKSWMESEFRVYSSANNCTFEYVSQPFLMDLEGKQGNFKNLREFVFKNIDGYFKIYSKHTPINLVRD
LPQGFSALEPLVDLPIGINITRFQTLLALHRSYLTPGDSSSGWTAGAAAYYVGYLQPRTFLLKYNENGTITD
AVDCALDPLSETKCTLKSFTVEKGIYQTSNFRVQPTESIVRFPNITNLCPFGEVFNATRFASVYAWNRKRISN
CVADYSVLYNSASFSTFKCYGVSPTKLNDLCFTNVYADSFVIRGDEVRQIAPGQTGKIADYNYKLPDDFTG
CVIAWNSNNLDSKVGGNYNYLYRLFRKSNLKPFERDISTEIYQAGSTPCNGVEGFNCYFPLQSYGFQPTNG
VGYQPYRVVVLSFELLHAPATVCGPKKSTNLVKNKCVNFNFNGLTGTGVLTESNKKFLPFQQFGRDIADTT
DAVRDPQTLEILDITPCSFGGVSVITPGTNTSNQVAVLYQDVNCTEVPVAIHADQLTPTWRVYSTGSNVFQT
RAGCLIGAEHVNNSYECDIPIGAGICASYQTQTNSPRRARSVASQSIIAYTMSLGAENSVAYSNNSIAIPTNFT
ISVTTEILPVSMTKTSVDCTMYICGDSTECSNLLLQYGSFCTQLNRALTGIAVEQDKNTQEVFAQVKQIYKT
PPIKDFGGFNFSQILPDPSKPSKRSFIEDLLFNKVTLADAGFIKQYGDCLGDIAARDLICAQKFNGLTVLPPLL
TDEMIAQYTSALLAGTITSGWTFGAGAALQIPFAMQMAYRFNGIGVTQNVLYENQKLIANQFNSAIGKIQD
SLSSTASALGKLQDVVNQNAQALNTLVKQLSSNFGAISSVLNDILSRLDKVEAEVQIDRLITGRLQSLQTYV
TQQLIRAAEIRASANLAATKMSECVLGQSKRVDFCGKGYHLMSFPQSAPHGVVFLHVTYVPAQEKNFTTA
PAICHDGKAHFPREGVFVSNGTHWFVTQRNFYEPQIITTDNTFVSGNCDVVIGIVNNTVYDPLQPELDSFKE
ELDKYFKNHTSPDVDLGDISGINASVVNIQKEIDRLNEVAKNLNESLIDLQELGKYEQYIKWPWYIWLGFIA
GLIAIVMVTIMLCCMTSCCSCLKGCCSCGSCCKFDEDDSEPVLKGVKLHYTLESGGGSAWSHPQFEKGGGS
GGGSGGSSAWSHPQFEK

**Name:** SC2.C1. 2P. TM4-2
**PDB:** 6XM4 (11)
QCVNLTTRTQLPPAYTNSFTRGVYYPDKVFRSSVLHSTQDLFLPFFSNVTWFHAIHVSGTNGTKRFDNPVLP
FNDGVYFASTEKSNIIRGWIFGTTLDSKTQSLLIVNNATNVVIKVCEFQFCNDPFLGVYYHKNNKSWMESEF
RVYSSANNCTFEYVSQPFLMDLEGKQGNFKNLREFVFKNIDGYFKIYSKHTPINLVRDLPQGFSALEPLVDL
PIGINITRFQTLLALHRSYLTPGDSSSGWTAGAAAYYVGYLQPRTFLLKYNENGTITDAVDCALDPLSETKC
TLKSFTVEKGIYQTSNFRVQPTESIVRFPNITNLCPFGEVFNATRFASVYAWNRKRISNCVADYSVLYNSASF
STFKCYGVSPTKLNDLCFTNVYADSFVIRGDEVRQIAPGQTGKIADYNYKLPDDFTGCVIAWNSNNLDSKV
GGNYNYLYRLFRKSNLKPFERDISTEIYQAGSTPCNGVEGFNCYFPLQSYGFQPTNGVGYQPYRVVVLSFEL
LHAPATVCGPKKSTNLVKNKCVNFNFNGLTGTGVLTESNKKFLPFQQFGRDIADTTDAVRDPQTLEILDITP
CSFGGVSVITPGTNTSNQVAVLYQDVNCTEVPVAIHADQLTPTWRVYSTGSNVFQTRAGCLIGAEHVNNSY
ECDIPIGAGICASYQTQTNSPGSASSVASQSIIAYTMSLGAENSVAYSNNSIAIPTNFTISVTTEILPVSMTKTS
VDCTMYICGDSTECSNLLLQYGSFCTQLNRALTGIAVEQDKNTQEVFAQVKQIYKTPPIKDFGGFNFSQILP
DPSKPSKRSFIEDLLFNKVTLADAGFIKQYGDCLGDIAARDLICAQKFNGLTVLPPLLTDEMIAQYTSALLA
GTITSGWTFGAGAALQIPFAMQMAYRFNGIGVTQNVLYENQKLIANQFNSAIGKIQDSLSSTASALGKLQD
VVNQNAQALNTLVKQLSSNFGAISSVLNDILSRLDPPEAEVQIDRLITGRLQSLQTYVTQQLIRAAEIRASAN
LAATKMSECVLGQSKRVDFCGKGYHLMSFPQSAPHGVVFLHVTYVPAQEKNFTTAPAICHDGKAHFPREG
VFVSNGTHWFVTQRNFYEPQIITTDNTFVSGNCDVVIGIVNNTVYDPLQPELDSFKEELDKYFKNHTSPDVD
LGDISGINASVVNIQKEIDRLNEVAKNLNESLIDLQELGKYEQGSGYIPEARDGQAYVRKDGEWVLLSTFLG
RSLEVLFQGPGHHHHHHHHSAWSHPQFEKGGGSGGGGSGGSAWSHPQFEK

Unresolved: 14-26, 70-79, 144-185, 251-263, 445-446, 677-688, 829-848, 1148-1288

**Name:** SC2.C1.TM4-2
**PDB:** 7KDH (13)
MFVFLVLLPLVSSQCVNLTTRTQLPPAYTNSFTRGVYYPDKVFRSSVLHSTQDLFLPFFSNVTWFHAIHVSG
TNGTKRFDNPVLPFNDGVYFASTEKSNIIRGWIFGTTLDSKTQSLLIVNNATNVVIKVCEFQFCNDPFLGVY
YHKNNKSWMESEFRVYSSANNCTFEYVSQPFLMDLEGKQGNFKNLREFVFKNIDGYFKIYSKHTPINLVRD

LPQGFSALEPLVDLPIGINITRFQTLLALHRSYLTPGDSSSGWTAGAAAYYVGYLQPRTFLLKYNENGTITD
AVDCALDPLSETKCTLKSFTVEKGIYQTSNFRVQPTESIVRFPNITNLCPFGEVFNATRFASVYAWNRKRISN
CVADYSVLYNSASFSTFKCYGVSPTKLNDLCFTNVYADSFVIRGDEVRQIAPGQTGKIADYNYKLPDDFTG
CVIAWNSNNLDSKVGGNYNYLYRLFRKSNLKPFERDISTEIYQAGSTPCNGVEGFNCYFPLQSYGFQPTNG
VGYQPYRVVVLSFELLHAPATVCGPKKSTNLVKNKCVNFNFNGLTGTGVLTESNKKFLPFQQFGRDIADTT
DAVRDPQTLEILDITPCSFGGVSVITPGTNTSNQVAVLYQDVNCTEVPVAIHADQLTPTWRVYSTGSNVFQT
RAGCLIGAEHVNNSYECDIPIGAGICASYQTQTNSPGSASSVASQSIIAYTMSLGAENSVAYSNNSIAIPTNFT
ISVTTEILPVSMTKTSVDCTMYICGDSTECSNLLLQYGSFCTQLNRALTGIAVEQDKNTQEVFAQVKQIYKT
PPIKDFGGFNFSQILPDPSKPSKRSFIEDLLFNKVTLADAGFIKQYGDCLGDIAARDLICAQKFNGLTVLPPLL
TDEMIAQYTSALLAGTITSGWTFGAGAALQIPFAMQMAYRFNGIGVTQNVLYENQKLIANQFNSAIGKIQD
SLSSTASALGKLQDVVNQNAQALNTLVKQLSSNFGAISSVLNDILSRLDKVEAEVQIDRLITGRLQSLQTYV
TQQLIRAAEIRASANLAATKMSECVLGQSKRVDFCGKGYHLMSFPQSAPHGVVFLHVTYVPAQEKNFTTA
PAICHDGKAHFPREGVFVSNGTHWFVTQRNFYEPQIITTDNTFVSGNCDVVIGIVNNTVYDPLQPELDSFKE
ELDKYFKNHTSPDVDLGDISGINASVVNIQKEIDRLNEVAKNLNESLIDLQELGKYEQGSGYIPEAPRDGQA
YVRKDGEWVLLSTFLGRSLEVLFQGPGHHHHHHHHSAWSHPQFEKGGGSGGGGSGGSAWSHPQFEK

Unresolved: 1-26, 70-81, 144-185, 243-262, 443-447, 471-489, 502, 621-640, 677-689, 812-852, 1148-1288

**Name:** BiPro-0
**PDB:** 6ZP7 (14)
MFVFLVLLPLVSSQCVNLTTRTQLPPAYTNSFTRGVYYPDKVFRSSVLHSTQDLFLPFFSNVTWFHAIHVSG
TNGTKRFDNPVLPFNDGVYFASTEKSNIIRGWIFGTTLDSKTQSLLIVNNATNVVIKVCEFQFCNDPFLGVY
YHKNNKSWMESEFRVYSSANNCTFEYVSQPFLMDLEGKQGNFKNLREFVFKNIDGYFKIYSKHTPINLVRD
LPQGFSALEPLVDLPIGINITRFQTLLALHRSYLTPGDSSSGWTAGAAAYYVGYLQPRTFLLKYNENGTITD
AVDCALDPLSETKCTLKSFTVEKGIYQTSNFRVQPTESIVRFPNITNLCPFGEVFNATRFASVYAWNRKRISN
CVADYSVLYNSASFSTFKCYGVSPTKLNDLCFTNVYADSFVIRGDEVRQIAPGQTGKIADYNYKLPDDFTG
CVIAWNSNNLDSKVGGNYNYLYRLFRKSNLKPFERDISTEIYQAGSTPCNGVEGFNCYFPLQSYGFQPTNG
VGYQPYRVVVLSFELLHAPATVCGPKKSTNLVKNKCVNFNFNGLTGTGVLTESNKKFLPFQQFGRDIADTT
DAVRDPQTLEILDITPCSFGGVSVITPGTNTSNQVAVLYQDVNCTEVPVAIHADQLTPTWRVYSTGSNVFQT
RAGCLIGAEHVNNSYECDIPIGAGICASYQTQTNSPRRARSVASQSIIAYTMSLGAENSVAYSNNSIAIPTNFT
ISVTTEILPVSMTKTSVDCTMYICGDSTECSNLLLQYGSFCTQLNRALTGIAVEQDKNTQEVFAQVKQIYKT
PPIKDFGGFNFSQILPDPSKPSKRSFIEDLLFNKVTLADAGFIKQYGDCLGDIAARDLICAQKFNGLTVLPPLL
TDEMIAQYTSALLAGTITSGWTFGAGAALQIPFAMQMAYRFNGIGVTQNVLYENQKLIANQFNSAIGKIQD
SLSSTASALGKLQDVVNQNAQALNTLVKQLSSNFGAISSVLNDILSRLDPPEAEVQIDRLITGRLQSLQTYVT
QQLIRAAEIRASANLAATKMSECVLGQSKRVDFCGKGYHLMSFPQSAPHGVVFLHVTYVPAQEKNFTTAP
AICHDGKAHFPREGVFVSNGTHWFVTQRNFYEPQIITTDNTFVSGNCDVVIGIVNNTVYDPLQPELDSFKEE
LDKYFKNHTSPDVDLGDISGINASVVNIQKEIDRLNEVAKNLNESLIDLQELGKYEQYIKWPWYIWLGFIAG
LIAIVMVTIMLCCMTSCCSCLKGCCSCGSCCKFDEDDSEPVLKGVKLHYT

Unresolved: 40-53, 105-134, 147-160, 216-234, 419-420, 594-612, 650-661, 802-824, 1122-1273

**Name**: SC2.TM5
**PDB ID**: 7C2L (35)
MFVFLVLLPLVSSQCVNLTTRTQLPPAYTNSFTRGVYYPDKVFRSSVLHSTQDLFLPFFSNVTWFHAIHVSG
TNGTKRFDNPVLPFNDGVYFASTEKSNIIRGWIFGTTLDSKTQSLLIVNNATNVVIKVCEFQFCNDPFLGVY
YHKNNKSWMESEFRVYSSANNCTFEYVSQPFLMDLEGKQGNFKNLREFVFKNIDGYFKIYSKHTPINLVRD
LPQGFSALEPLVDLPIGINITRFQTLLALHRSYLTPGDSSSGWTAGAAAYYVGYLQPRTFLLKYNENGTITD
AVDCALDPLSETKCTLKSFTVEKGIYQTSNFRVQPTESIVRFPNITNLCPFGEVFNATRFASVYAWNRKRISN
CVADYSVLYNSASFSTFKCYGVSPTKLNDLCFTNVYADSFVIRGDEVRQIAPGQTGKIADYNYKLPDDFTG
CVIAWNSNNLDSKVGGNYNYLYRLFRKSNLKPFERDISTEIYQAGSTPCNGVEGFNCYFPLQSYGFQPTNG
VGYQPYRVVVLSFELLHAPATVCGPKKSTNLVKNKCVNFNFNGLTGTGVLTESNKKFLPFQQFGRDIADTT
DAVRDPQTLEILDITPCSFGGVSVITPGTNTSNQVAVLYQDVNCTEVPVAIHADQLTPTWRVYSTGSNVFQT
RAGCLIGAEHVNNSYECDIPIGAGICASYQTQTNSPRGSASSVASQSIIAYTMSLGAENSVAYSNNSIAIPTNF
TISVTTEILPVSMTKTSVDCTMYICGDSTECSNLLLQYGSFCTQLNRALTGIAVEQDKNTQEVFAQVKQIYK
TPPIKDFGGFNFSQILPDPSKPSKRSFIEDLLFNKVTLADAGFIKQYGDCLGDIAARDLICAQKFNGLTVLPPL
LTDEMIAQYTSALLAGTITSGWTFGAGAALQIPFAMQMAYRFNGIGVTQNVLYENQKLIANQFNSAIGKIQ

DSLSSTASALGKLQDVVNQNAQALNTLVKQLSSNFGAISSVLNDILSRLDPPEAEVQIDRLITGRLQSLQTY
VTQQLIRAAEIRASANLAATKMSECVLGQSKRVDFCGKGYHLMSFPQSAPHGVVFLHVTYVPAQEKNFTT
APAICHDGKAHFPREGVFVSNGTHWFVTQRNFYEPQIITTDNTFVSGNCDVVIGIVNNTVYDPLQPELDSFK
EELDKYFKNHTSPDVDLGDISGINASVVNIQKEIDRLNEVAKNLNESLIDLQELGKYEQYIKWPWYIWLGFI
AGLIAIVMVTIMLCCMTSCCSCLKGCCSCGSCCKFDEDDSEPVLKGVKLHYTLEDYKDDDDK

### 2. SARS-CoV Protein Sequences

**SARS-CoV WT (Uniprot ID P59594) (15)**
MFIFLLFLTLTSGSDLDRCTTFDDVQAPNYTQHTSSMRGVYYPDEIFRSDTLYLTQDLFLPFYSNVTGFHTIN
HTFGNPVIPFKDGIYFAATEKSNVVRGWVFGSTMNNKSQSVIIINNSTNVVIRACNFELCDNPFFAVSKPMG
TQTHTMIFDNAFNCTFEYISDAFSLDVSEKSGNFKHLREFVFKNKDGFLYVYKGYQPIDVVRDLPSGFNTLK
PIFKLPLGINITNFRAILTAFSPAQDIWGTSAAAYFVGYLKPTTFMLKYDENGTITDAVDCSQNPLAELKCSV
KSFEIDKGIYQTSNFRVVPSGDVVRFPNITNLCPFGEVFNATKFPSVYAWERKKISNCVADYSVLYNSTFFST
FKCYGVSATKLNDLCFSNVYADSFVVKGDDVRQIAPGQTGVIADYNYKLPDDFMGCVLAWNTRNIDATS
TGNYNYKYRYLRHGKLRPFERDISNVPFSPDGKPCTPPALNCYWPLNDYGFYTTTGIGYQPYRVVVLSFEL
LNAPATVCGPKLSTDLIKNQCVNFNFNGLTGTGVLTPSSKRFQPFQQFGRDVSDFTDSVRDPKTSEILDISPC
SFGGVSVITPGTNASSEVAVLYQDVNCTDVSTAIHADQLTPAWRIYSTGNNVFQTQAGCLIGAEHVDTSYE
CDIPIGAGICASYHTVSLLRSTSQKSIVAYTMSLGADSSIAYSNNTIAIPTNFSISITTEVMPVSMAKTSVDCN
MYICGDSTECANLLLQYGSFCTQLNRALSGIAAEQDRNTREVFAQVKQMYKTPTLKYFGGFNFSQILPDPL
KPTKRSFIEDLLFNKVTLADAGFMKQYGECLGDINARDLICAQKFNGLTVLPPLLTDDMIAAYTAALVSGT
ATAGWTFGAGAALQIPFAMQMAYRFNGIGVTQNVLYENQKQIANQFNKAISQIQESLTTTSTALGKLQDV
VNQNAQALNTLVKQLSSNFGAISSVLNDILSRLDKVEAEVQIDRLITGRLQSLQTYVTQQLIRAAEIRASANL
AATKMSECVLGQSKRVDFCGKGYHLMSFPQAAPHGVVFLHVTYVPSQERNFTTAPAICHEGKAYFPREGV
FVFNGTSWFITQRNFFSPQIITTDNTFVSGNCDVVIGIINNTVYDPLQPELDSFKEELDKYFKNHTSPDVDLG
DISGINASVVNIQKEIDRLNEVAKNLNESLIDLQELGKYEQYIKWPWYVWLGFIAGLIAIVMVTILLCCMTS
CCSCLKGACSCGSCCKFDEDDSEPVLKGVKLHYT

**Name:** SC1. TM1
**PDB:** 6ACD (35)
MFIFLLFLTLTSGSDLDRCTTFDDVQAPNYTQHTSSMRGVYYPDEIFRSDTLYLTQDLFLPFYSNVTGFHTIN
HTFGNPVIPFKDGIYFAATEKSNVVRGWVFGSTMNNKSQSVIIINNSTNVVIRACNFELCDNPFFAVSKPMG
TQTHTMIFDNAFNCTFEYISDAFSLDVSEKSGNFKHLREFVFKNKDGFLYVYKGYQPIDVVRDLPSGFNTLK
PIFKLPLGINITNFRAILTAFSPAQDIWGTSAAAYFVGYLKPTTFMLKYDENGTITDAVDCSQNPLAELKCSV
KSFEIDKGIYQTSNFRVVPSGDVVRFPNITNLCPFGEVFNATKFPSVYAWERKKISNCVADYSVLYNSTFFST
FKCYGVSATKLNDLCFSNVYADSFVVKGDDVRQIAPGQTGVIADYNYKLPDDFMGCVLAWNTRNIDATS
TGNYNYKYRYLRHGKLRPFERDISNVPFSPDGKPCTPPALNCYWPLNDYGFYTTTGIGYQPYRVVVLSFEL
LNAPATVCGPKLSTDLIKNQCVNFNFNGLTGTGVLTPSSKRFQPFQQFGRDVSDFTDSVRDPKTSEILDISPC
SFGGVSVITPGTNASSEVAVLYQDVNCTDVSTAIHADQLTPAWRIYSTGNNVFQTQAGCLIGAEHVDTSYE
CDIPIGAGICASYHTVSLLRSTSQKSIVAYTMSLGADSSIAYSNNTIAIPTNFSISITTEVMPVSMAKTSVDCN
MYICGDSTECANLLLQYGSFCTQLNRALSGIAAEQDRNTREVFAQVKQMYKTPTLKYFGGFNFSQILPDPL
KPTKRSFIEDLLFNKVTLADAGFMKQYGECLGDINARDLICAQKFNGLTVLPPLLTDDMIAAYTAALVSGT
ATAGWTFGAGAALQIPFAMQMAYRFNGIGVTQNVLYENQKQIANQFNKAISQIQESLTTTSTALGKLQDV
VNQNAQALNTLVKQLSSNFGAISSVLNDILSRLDKVEAEVQIDRLITGRLQSLQTYVTQQLIRAAEIRASANL
AATKMSECVLGQSKRVDFCGKGYHLMSFPQAAPHGVVFLHVTYVPSQERNFTTAPAICHEGKAYFPREGV
FVFNGTSWFITQRNFFSPQIITTDNTFVSGNCDVVIGIINNTVYDPLQPELDSFKEELDKYFKNHTSPDVDLG
DISGINASVVNIQKEIDRLNEVAKNLNESLIDLQELGKYEQYIKWPWSHPQFEK

**Name:** SC1.S1.TM2
**PDB**: 6NB6 (35)
MGILPSPGMPALLSLVSLLSVLLMGCVAETGTSDLDRCTTFDDVQAPNYTQHTSSMRGVYYPDEIFRSDTL
YLTQDLFLPFYSNVTGFHTINHTFDNPVIPFKDGIYFAATEKSNVVRGWVFGSTMNNKSQSVIIINNSTNVVI
RACNFELCDNPFFAVSKPMGTQTHTMIFDNAFNCTFEYISDAFSLDVSEKSGNFKHLREFVFKNKDGFLYV
YKGYQPIDVVRDLPSGFNTLKPIFKLPLGINITNFRAILTAFSPAQDTWGTSAAAYFVGYLKPTTFMLKYDE
NGTITDAVDCSQNPLAELKCSVKSFEIDKGIYQTSNFRVVPSGDVVRFPNITNLCPFGEVFNATKFPSVYAW

ERKKISNCVADYSVLYNSTFFSTFKCYGVSATKLNDLCFSNVYADSFVVKGDDVRQIAPGQTGVIADYNYK
LPDDFMGCVLAWNTRNIDATSTGNYNYKYRYLRHGKLRPFERDISNVPFSPDGKPCTPPALNCYWPLNDY
GFYTTTGIGYQPYRVVVLSFELLNAPATVCGPKLSTDLIKNQCVNFNFNGLTGTGVLTPSSKRFQPFQQFGR
DVSDFTDSVRDPKTSEILDISPCSFGGVSVITPGTNASSEVAVLYQDVNCTDVSTAIHADQLTPAWRIYSTGN
NVFQTQAGCLIGAEHVDTSYECDIPIGAGICASYHTVSLLRSTSQKSIVAYTMSLGADSSIAYSNNTIAIPTNF
SISITTEVMPVSMAKTSVDCNMYICGDSTECANLLLQYGSFCTQLNRALSGIAAEQDRNTREVFAQVKQMY
KTPTLKYFGGFNFSQILPDPLKPTKRSFIEDLLFNKVTLADAGFMKQYGECLGDINARDLICAQKFNGLTVL
PPLLTDDMIAAYTAALVSGTATAGWTFGAGAALQIPFAMQMAYRFNGIGVTQNVLYENQKQIANQFNKAI
SQIQESLTTTSTALGKLQDVVNQNAQALNTLVKQLSSNFGAISSVLNDILSRLDPPEAEVQIDRLITGRLQSL
QTYVTQQLIRAAEIRASANLAATKMSECVLGQSKRVDFCGKGYHLMSFPQAAPHGVVFLHVTYVPSQERN
FTTAPAICHEGKAYFPREGVFVFNGTSWFITQRNFFSPQIITTDNTFVSGNCDVVIGIINNTVYDPLQPELDSF
KEELDKYFKNHTSPDVDLGDISGINASVVNIQKEIDRLNEVAKNLNESLIDLQELGKYEQYIKGSGRENLYF
QGGGGSGYIPEAPRDGQAYVRKDGEWVLLSTFLGHHHHHHHH

**3. MERS-CoV Protein Sequences**

**MERS-CoV WT (Uniprot ID K9N5Q8) (16)**
MIHSVFLLMFLLTPTESYVDVGPDSVKSACIEVDIQQTFFDKTWPRPIDVSKADGIIYPQGRTYSNITITYQGL
FPYQGDHGDMYVYSAGHATGTTPQKLFVANYSQDVKQFANGFVVRIGAAANSTGTVIISPSTSATIRKIYP
AFMLGSSVGNFSDGKMGRFFNHTLVLLPDGCGTLLRAFYCILEPRSGNHCPAGNSYTSFATYHTPATDCSD
GNYNRNASLNSFKEYFNLRNCTFMYTYNITEDEILEWFGITQTAQGVHLFSSRYVDLYGGNMFQFATLPVY
DTIKYYSIIPHSIRSIQSDRKAWAAFYVYKLQPLTFLLDFSVDGYIRRAIDCGFNDLSQLHCSYESFDVESGV
YSVSSFEAKPSGSVVEQAEGVECDFSPLLSGTPPQVYNFKRLVFTNCNYNLTKLLSLFSVNDFTCSQISPAAI
ASNCYSSLILDYFSYPLSMKSDLSVSSAGPISQFNYKQSFSNPTCLILATVPHNLTTITKPLKYSYINKCSRFLS
DDRTEVPQLVNANQYSPCVSIVPSTVWEDGDYYRKQLSPLEGGGWLVASGSTVAMTEQLQMGFGITVQY
GTDTNSVCPKLEFANDTKIASQLGNCVEYSLYGVSGRGVFQNCTAVGVRQQRFVYDAYQNLVGYYSDDG
NYYCLRACVSVPVSVIYDKETKTHATLFGSVACEHISSTMSQYSRSTRSMLKRRDSTYGPLQTPVGCVLGL
VNSSLFVEDCKLPLGQSLCALPDTPSTLTPRSVRSVPGEMRLASIAFNHPIQVDQLNSSYFKLSIPTNSFGVTQ
EYIQTTIQKVTVDCKQYVCNGFQKCEQLLREYGQFCSKINQALHGANLRQDDSVRNLFASVKSSQSSPIIPG
FGGDFNLTLLEPVSISTGSRSARSAIEDLLFDKVTIADPGYMQGYDDCMQQGPASARDLICAQYVAGYKVL
PPLMDVNMEAAYTSSLLGSIAGVGWTAGLSSFAAIPFAQSIFYRLNGVGITQQVLSENQKLIANKFNQALGA
MQTGFTTTNEAFHKVQDAVNNNAQALSKLASELSNTFGAISASIGDIIQRLDVLEQDAQIDRLINGRLTTLN
AFVAQQLVRSESAALSAQLAKDKVNECVKAQSKRSGFCGQGTHIVSFVVNAPNGLYFMHVGYYPSNHIEV
VSAYGLCDAANPTNCIAPVNGYFIKTNNTRIVDEWSYTGSSFYAPEPITSLNTKYVAPQVTYQNISTNLPPPL
LGNSTGIDFQDELDEFFKNVSTSIPNFGSLTQINTTLLDLTYEMLSLQQVVKALNESYIDLKELGNYTYYNK
WPWYIWLGFIAGLVALALCVFFILCCTGCGTNCMGK
LKCNRCCDRYEEYDLEPHKVHVH

**Name:** MC. SD. TM1
**PDB**: 5X5C & 5X5F (36)
YVDVGPDSVKSACIEVDIQQTFFDKTWPRPIDVSKADGIIYPQGRTYSNITITYQGLFPYQGDHGDMYVYSA
GHATGTTPQKLFVANYSQDVKQFANGFVVRIGAAANSTGTVIISPSTSATIRKIYPAFMLGSSVGNFSDGKM
GRFFNHTLVLLPDGCGTLLRAFYCILEPRSGNHCPAGNSYTSFATYHTPATDCSDGNYNRNASLNSFKEYF
NLRNCTFMYTYNITEDEILEWFGITQTAQGVHLFSSRYVDLYGGNMFQFATLPVYDTIKYYSIIPHSIRSIQS
DRKAWAAFYVYKLQPLTFLLDFSVDGYIRRAIDCGFNDLSQLHCSYESFDVESGVYSVSSFEAKPSGSVVE
QAEGVECDFSPLLSGTPPQVYNFKRLVFTNCNYNLTKLLSLFSVNDFTCSQISPAAIASNCYSSLILDYFSYPL
SMKSDLSVSSAGPISQFNYKQSFSNPTCLILATVPHNLTTITKPLKYSYINKCSRLLSDDRTEVPQLVNANQY
SPCVSIVPSTVWEDGDYYRKQLSPLEGGGWLVASGSTVAMTEQLQMGFGITVQYGTDTNSVCPKLEFAND
TKIASQLGNCVEYSLYGVSGRGVFQNCTAVGVRQQRFVYDAYQNLVGYYSDDGNYYCLRACVSVPVSVI
YDKETKTHATLFGSVACEHISSTMSQYSRSTRSMLKRRDSTYGPLQTPVGCVLGLVNSSLFVEDCKLPLGQ
SLCALPDTPSTLTPRSVSSVPGEMRLASIAFNHPIQVDQLNSSYFKLSIPTNFSFGVTQEYIQTTIQKVTVDCK
QYVCNGFQKCEQLLREYGQFCSKINQALHGANLRQDDSVRNLFASVKSSQSSPIIPGFGGDFNLTLLEPVSIS
TGSRSARSAIEDLLFDKVTIADPGYMQGYDDCMQQGPASARDLICAQYVAGYKVLPPLMDVNMEAAYTS
SLLGSIAGVGWTAGLSSFAAIPFAQSIFYRLNGVGITQQVLSENQKLIANKFNQALGAMQTGFTTTNEAFQK
VQDAVNNNAQALSKLASELSNTFGAISASIGDIIQRLDVLEQDAQIDRLINGRLTTLNAFVAQQLVRSESAA

LSAQLAKDKVNECVKAQSKRSGFCGQGTHIVSFVVNAPNGLYFMHVGYYPSNHIEVVSAYGLCDAANPT
NCIAPVNGYFIKTNNTRIVDEWSYTGSSFYAPEPITSLNTKYVAPQVTYQNISTNLPPPLLGNSTGIDFQDELD
EFFKNVSTSIPNFGSLTQINTTLLDLTYEMLSLQQVVKALNESYIDLKELGNYTYYNKEFRLVPRGSPGSGYI
PEAPRDGQAYVRKDGEWVLLSTFLGHHHHHH

**Name:** MC.TM2
**PDB**: 5W9K (37)
MIHSVFLLMFLLTPTESYVDVGPDSVKSACIEVDIQQTFFDKTWPRPIDVSKADGIIYPQGRTYSNITITYQGL
FPYQGDHGDMYVYSAGHATGTTPQKLFVANYSQDVKQFANGFVVRIGAAANSTGTVIISPSTSATIRKIYP
AFMLGSSVGNFSDGKMGRFFNHTLVLLPDGCGTLLRAFYCILEPRSGNHCPAGNSYTSFATYHTPATDCSD
GNYNRNASLNSFKEYFNLRNCTFMYTYNITEDEILEWFGITQTAQGVHLFSSRYVDLYGGNMFQFATLPVY
DTIKYYSIIPHSIRSIQSDRKAWAAFYVYKLQPLTFLLDFSVDGYIRRAIDCGFNDLSQLHCSYESFDVESGV
YSVSSFEAKPSGSVVEQAEGVECDFSPLLSGTPPQVYNFKRLVFTNCNYNLTKLLSLFSVNDFTCSQISPAAI
ASNCYSSLILDYFSYPLSMKSDLSVSSAGPISQFNYKQSFSNPTCLILATVPHNLTTITKPLKYSYINKCSRFLS
DDRTEVPQLVNANQYSPCVSIVPSTVWEDGDYYRKQLSPLEGGGWLVASGSTVAMTEQLQMGFGITVQY
GTDTNSVCPKLEFANDTKIASQLGNCVEYSLYGVSGRGVFQNCTAVGVRQQRFVYDAYQNLVGYYSDDG
NYYCLRACVSVPVSVIYDKETKTHATLFGSVACEHISSTMSQYSRSTRSMLKRRDSTYGPLQTPVGCVLGL
VNSSLFVEDCKLPLGQSLCALPDTPSTLTPASVGSVPGEMRLASIAFNHPIQVDQLNSSYFKLSIPTNFSFGVT
QEYIQTTIQKVTVDCKQYVCNGFQKCEQLLREYGQFCSKINQALHGANLRQDDSVRNLFASVKSSQSSPIIP
GFGGDFNLTLLEPVSISTGSRSARSAIEDLLFDKVTIADPGYMQGYDDCMQQGPASARDLICAQYVAGYKV
LPPLMDVNMEAAYTSSLLGSIAGVGWTAGLSSFAAIPFAQSIFYRLNGVGITQQVLSENQKLIANKFNQALG
AMQTGFTTTNEAFHKVQDAVNNNAQALSKLASELSNTFGAISASIGDIIQRLDPPEQDAQIDRLINGRLTTL
NAFVAQQLVRSESAALSAQLAKDKVNECVKAQSKRSGFCGQGTHIVSFVVNAPNGLYFMHVGYYPSNHIE
VVSAYGLCDAANPTNCIAPVNGYFIKTNNTRIVDEWSYTGSSFYAPEPITSLNTKYVAPQVTYQNISTNLPPP
LLGNSTGIDFQDELDEFFKNVSTSIPNFGSLTQINTTLLDLTYEMLSLQQVVKALNESYIDLKELGNYTYGSG
YIPEAPRDGQAYVRKDGEWVLLSTFLGRSLEVLFQ

### 4. Supplementary Table 1

See attached file titled Supplementary_Table_1.xlsx for extensive review of individual binding properties and epitopes of over 40 antibodies and their related pdb identifiers. References for the table are located in this document.

### 5. Protein Sequence Alignment and Comparison

Multiple sequence alignment of spike protein sequences are attached in file Spike_Protein_Alignment_Tarakanova.clustal_num

### 6. Dynamic Domain & NMA Videos

Like the Dynamic Domain index files, videos showing the NMA trajectory and highlighted dynamic domains are labeled [PDB ID].mp4

### 7. Results of artificial controls

The graphical results of the artificial controls used in **Results section 3.4** are listed below:

i. **SC2.S2.TM1-1'**
Combined Unresolved regions (WT'A)-: 1-26, 70-81, 114-115, 144-185, 243-262, 443-489, 502, 621-640, 677-689, 812, 828-854, 1148-end

<u>Combined unresolved regions (SC2.S2.TM1-1')</u>: 1-26, 70-81,114-115,144-185,243-262, 443-489,502,618-640, 677-698, 812, 828-859, 1148-end

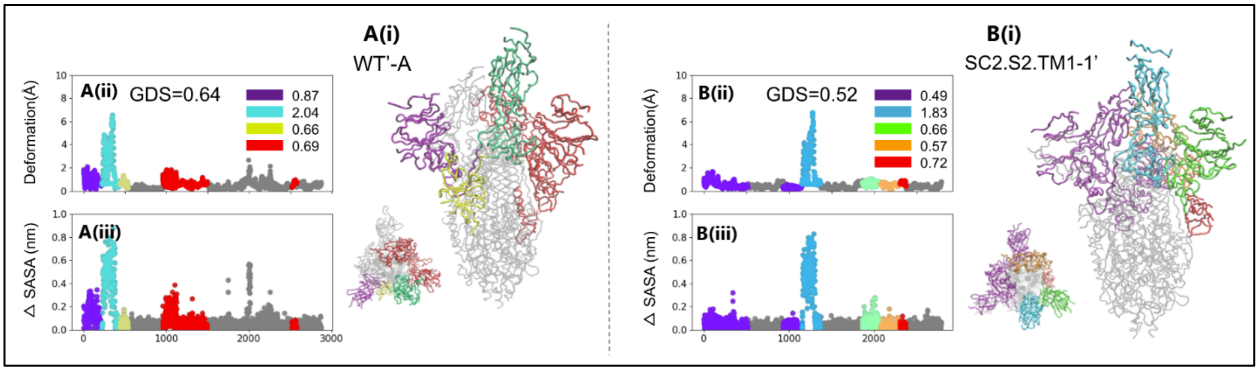


**SI Figure 1**: The domain dynamics associated with **(A)** WT'-A and **(B)** SC2.S2.TM1-1' ANMs. The PDB ID, global dynamics score (GDS), local dynamics scores (LDSs), deformation profile **(ii)**, and Δ SASA profile **(iii)** is listed for each structure. On each 3-D structure and profile, identified dynamic domains are labeled in different colors and their LDSs are listed in each legend. The WT dynamic domain breakdown is contained in **Figure 1** and SC2.S2.TM1-1 in **Figure 4.**

**ii. SC2.C1.2P' & SC2.C1.2P.TM4',**

<u>Combined unresolved regions</u>:12-26, 70-81, 114-115, 144-165, 173-185, 243-263, 445-446, 621-640, 613-640, 673-690, 812-854, 1148-end

### 8. Derived features used for thermal stability training

The features used to train the thermal stability predictor are those contained in the original combined data set and

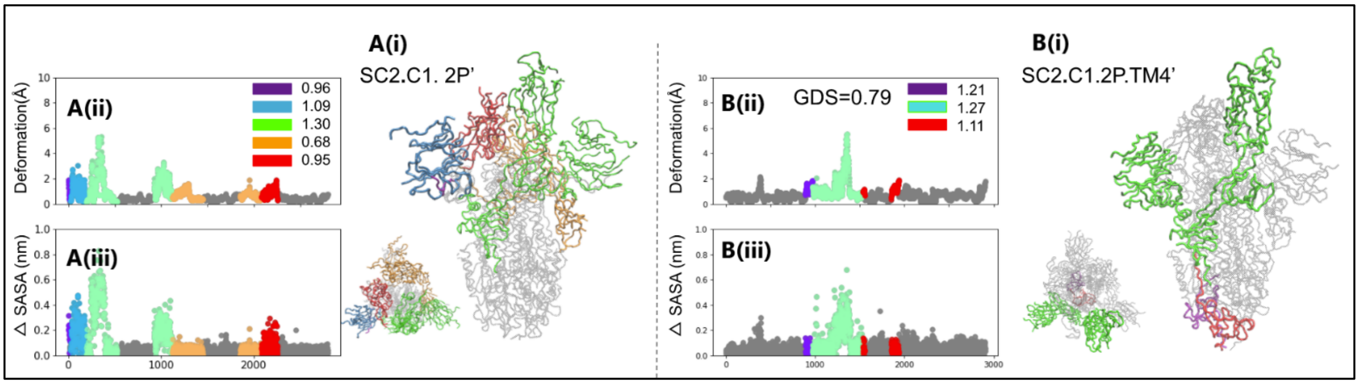


**SI Figure 2:** The domain dynamics associated with **(A)** SC2.C1.2P' and **(B)** SC2.C1.2P.TM4' ANMs. The PDB ID, global dynamics score (GDS), local dynamics scores (LDSs), deformation profile **(ii)**, and Δ SASA profile **(iii)** is listed for each structure. On each 3-D structure and profile, identified dynamic domains are labeled in different colors and their LDSs are listed in each legend. The SC2.C1.2P and SC2.C1.2P.TM4 dynamic domain breakdowns are located in **Figure 4.**

also those computed by the following resources. The Amber software (39) was used to find bond length, bond angle, dihedral angles, van der walls contributions, electrostatic contributions, polar solvation, total gas free energy, total solvation free energy, and total system energy. The FoldX software (40) was used to find solvation energy for polar groups, solvation energy for apolar groups, water bridge hydrogen bonding, intra-molecule hydrogen bonding, electrostatic interactions between charged groups, and atomic clash overlaps. The remaining features are outlined in **SI Table A**. Amino acid-related biological features are found using AAindex (42). Disorder and Relative Surface

Accessibility are found using the SCRATCH webserver (43). All remaining features in **SI Table A** are found using the Expasy ProtScale tool (41).

**SI Table A**: Summary of the biochemical, structural, and biological features used for thermal stability predictor training.

| **Biochemical Features** | **Structural features** | **Biological Features** |
|---|---|---|
| Molecular Weight | Solvent Accessibility | Mobility of Amino Acids |
| Hydrophobicity Index | 3-State Secondary Structure | Codon Amount per Amino Acid |
| Side Chain pKa | Bulkiness | |
| Frequencies of buried and exposed sequence | Buried Area from Standard to Folded State | |
| Electrostatic Charge | Disorder | |
| | Flexibility Index | |
| | Relative Surface Accessibility | |
| | Absolute Surface Accessibility | |
